# High-density peptide arrays detect tuberculosis through immune remodeling, not only antigen recognition alone

**DOI:** 10.64898/2026.04.16.718855

**Authors:** Dimitry Schmidt, Sergey Biniaminov, Nathalie Biniaminov, Clemens von Bojnicic-Kninski, Roman Popov, Josef Maier, Hubert Bernauer, Johanna Griesbaum, Nicole Schneiderhan-Marra, Alex Dulovic, Alexander Nesterov-Mueller

**Affiliations:** Institute of Microstructure Technology, Karlsruhe Institute of Technology, Hermann-von-Helmholtz-Platz 1, Eggenstein-Leopoldshafen, Germany; HS Analysis, Haid-und-Neu-Straße 7, 76131 Karlsruhe, Germany; axxelera, Karl-Floesser-Str. 14, Karlsruhe, Germany; ATG:biosynthetics GmbH, Weberstraße 40, Merzhausen, Germany; IstLS, Härlestr. 24/1, Oberndorf a.N., Germany; NMI Natural and Medical Sciences Institute at the University of Tübingen, Reutlingen, Germany

**Author notes:** Corresponding Authors: Alexander Nesterov-Mueller,; Alex Dulovic. shared last authors (Alex Dulovic, Alexander Nesterov-Mueller).

## Abstract

Serological diagnostics for tuberculosis rely on pathogen-derived antigens to detect infection-specific antibodies. Whether chronic TB infection also reshapes the global topology of the antibody repertoire remains largely unexplored. Here we profile serum antibody binding across 6,936 peptides in 105 individuals from three countries using two complementary libraries: *Mycobacterium tuberculosis* peptides (TBC) and a resemblance-ranking library representing the human self-proteome (RRL). We construct a five-dimensional immune state vector from distributional binding properties and map individual sera into an immune phase space. A remodeling classifier achieves virtually identical performance on pathogen-derived and host-derived peptides (AUC 0.63–0.73), demonstrating that the diagnostic signal arises from global repertoire restructuring rather than antigen-specific recognition. HIV co-infection partially masks this signal; restricting analysis to HIV-negative individuals increases AUC to 0.73 (permutation p = 0.005) and enables detection of smear-negative TB (AUC = 0.83, specificity 0.95 with three peptides). Phase-space projections reveal that TB severity maps onto a continuous remodeling gradient, with smear-negative patients occupying intermediate positions between healthy controls and smear-positive cases. These findings position high-density peptide arrays as sensors of antibody repertoire topology, enabling detection of chronic immune states beyond antigen-specific recognition.

## Introduction

Serological diagnostics have historically been built upon an antigen-centric view of immunity: disease-associated information is assumed to reside in antibodies recognizing discrete pathogen-derived epitopes, and assays are designed accordingly (1). While this framework has been highly successful for identifying exposure to infectious agents, it implicitly reduces the antibody repertoire to a collection of independent epitope-specific signals. Yet the adaptive immune system is not merely a catalog of specificities; it is a dynamic, high-dimensional system whose structure reflects clonal selection, affinity maturation, and immune history. Recent developments in systems serology have begun to capture this complexity by profiling polyclonal antibody features — including subclass, glycosylation, and Fc-mediated effector functions — as high-dimensional immune signatures (2-5). However, these approaches remain anchored to predefined antigen panels and do not directly interrogate the global topology of antibody binding distributions. A fundamental question therefore remains open: can disease states be detected not through what the immune system recognizes, but through how the recognition landscape is organized?

The antibody repertoire emerges from V(D)J recombination, somatic hypermutation, and germinal center selection — processes that collectively generate a structured distribution of B-cell clones (5, 6). Crucially, chronic antigen exposure can reshape this distribution through clonal expansion and selective focusing, reducing effective diversity while amplifying dominant clones (7). The distinction between acute and chronic infections is critical here. Acute infections produce transient immune perturbations that resolve within weeks; the antibody landscape returns to baseline after pathogen clearance. Chronic infections, by contrast, impose sustained selective pressure on the B-cell repertoire, potentially driving it into a new quasi-stationary state characterized by altered diversity, inequality, and geometric organization. Immune repertoire sequencing studies have confirmed that infections, vaccination, and autoimmunity induce measurable shifts in diversity, clonality, and network structure (8-10). These findings suggest that chronic disease states may be encoded not only in individual antigen-specific responses but also in the global topology of the repertoire itself. However, repertoire sequencing provides a structural readout of B-cell clones rather than a functional measurement of circulating antibody binding patterns. Whether serological profiling can detect global repertoire remodeling, or whether detection is possible without pathogen-derived antigens, has not been tested.

Tuberculosis, caused by *Mycobacterium tuberculosis*, offers a particularly compelling model for investigating this question. TB is characterized by prolonged antigen persistence within granulomatous lesions and sustained immune activation over months to years. These conditions can drive extended germinal center reactions (6, 7), promoting affinity maturation and selective expansion of dominant B-cell clones. Because affinity-matured antibodies frequently exhibit degrees of polyreactivity (11, 12), expansion of dominant clones can redistribute binding across broad regions of peptide sequence space — including host-like sequences — rather than remaining confined to single epitopes. Recent systems serology studies have demonstrated that antibodies play functional roles in TB immunity beyond classical neutralization (13), yet the possibility that TB induces measurable remodeling of the global antibody binding landscape has received limited attention. The clinical relevance of such remodeling is underscored by the limitations of current TB diagnostics. Sputum smear microscopy — the most widely available test in resource-limited settings — fails to detect smear-negative, culture-positive (S−C+) patients, who represent approximately 40% of pulmonary TB cases and remain a major source of ongoing transmission.

If immune remodeling signatures could detect these patients, this would address a critical gap in the TB diagnostic cascade. However, the challenge is compounded by HIV co-infection, which remains prevalent in TB-endemic regions. HIV profoundly disrupts B-cell homeostasis, inducing hypergammaglobulinemia, increased polyreactivity, and altered germinal center architecture (14-16). These perturbations can distort repertoire organization, potentially obscuring more subtle infection-specific immune signals. Disentangling TB-associated remodeling from HIV-driven immune disruption therefore provides a stringent test of whether systemic immune states can be resolved serologically.

High-density peptide arrays provide a unique experimental platform for this investigation. By simultaneously measuring antibody binding across thousands of peptides, these arrays generate high-dimensional binding vectors for individual serum samples. Traditionally used for epitope mapping and antigen discovery (17-20), peptide arrays have been interpreted as collections of discrete antigenic probes. However, high-density peptide arrays can also be interpreted as statistical samplers of antibody binding space rather than as collections of discrete antigenic probes. Under this interpretation, distributional properties of binding intensities — such as entropy, inequality, and multivariate displacement — become measurable surrogates of underlying repertoire topology. Immunosignaturing studies using random-sequence peptide microarrays have demonstrated that disease classification can be achieved from collective binding patterns without knowledge of specific epitopes (21-23), and phage immunoprecipitation approaches have enabled comprehensive serological profiling at population scale (24, 25). Critically, these approaches have not yet been combined with an explicit geometric framework in which individual sera are mapped into a low-dimensional immune state space and disease is detected as displacement within that space — an approach analogous to phase-space representations in dynamical systems.

To experimentally separate antigen-specific recognition from global repertoire remodeling, we employ a resemblance-ranking peptide library (RRL) designed to approximate the statistical structure of the human peptidome (26). Unlike random peptide collections, the RRL was constructed by an algorithm that selects 10-mer peptides maximally representing the k-mer fragment diversity of the entire human proteome. These peptides have no designed relationship to any pathogen; they probe the host-directed, self-reactive component of serum antibody binding. If TB classification succeeds on this host-proteome library at the same accuracy as on pathogen-derived peptides, this constitutes direct evidence that the diagnostic signal arises from global immune remodeling rather than antigen-specific recognition.

Here, we profile serum antibody binding using high-density peptide arrays comprising two complementary libraries: pathogen-derived peptides from *M. tuberculosis* and host-proteome-derived peptides (Figure 1). These two libraries provide fundamentally different views of the immune response: pathogen-derived peptides probe *pathogen-directed* antibody reactivity, while host-proteome-derived peptides probe *host-directed* (self-reactive) antibody patterns. If TB-associated immune changes are detectable through both libraries equally, this constitutes evidence for global remodeling beyond pathogen-specific immunity. We construct a five-dimensional immune state vector from distributional binding properties and show that a remodeling classifier achieves equivalent performance on both libraries.

**Figure 1.**
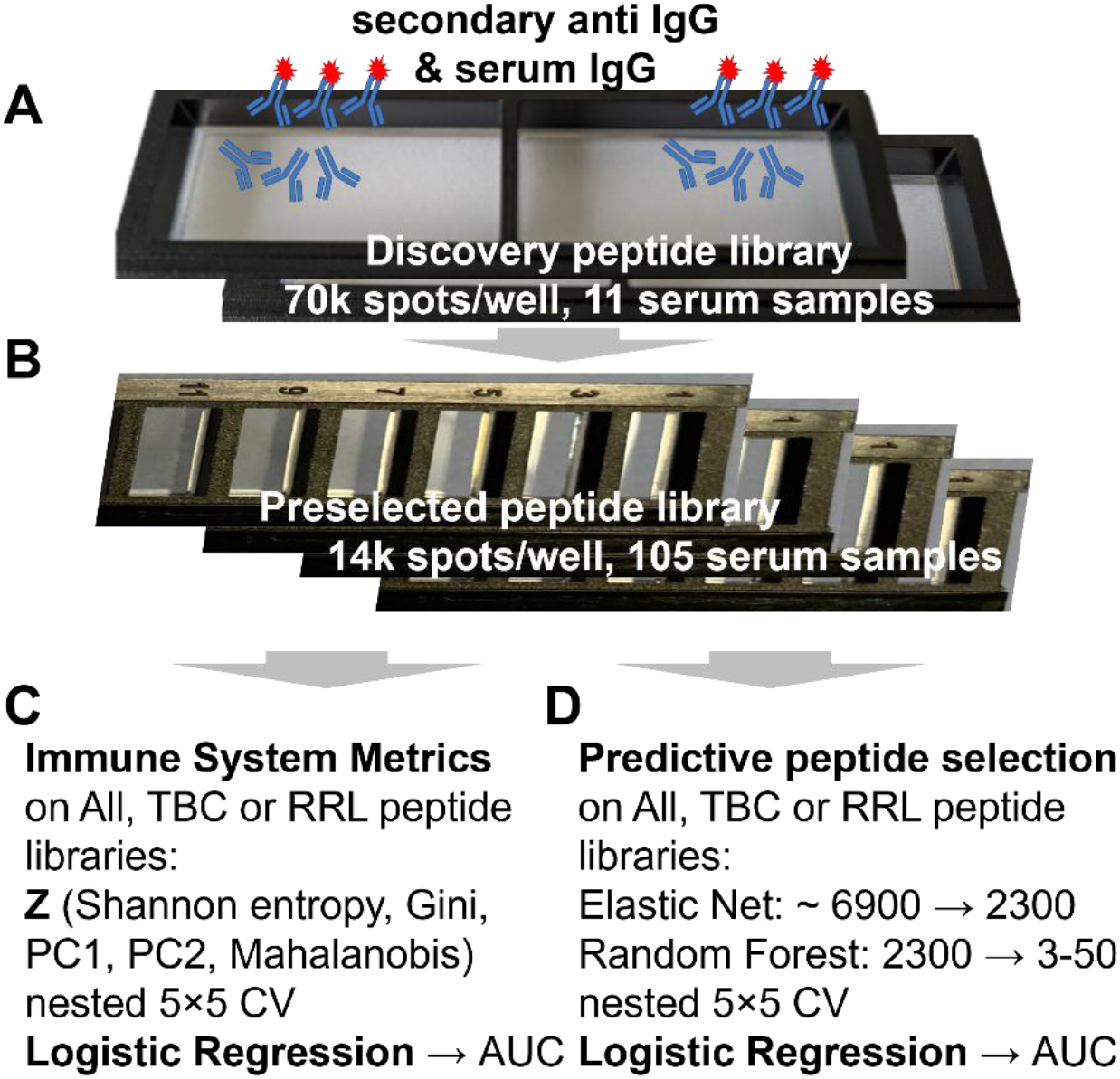
Study design. (A) Discovery peptide library (70k spots/well, 11 serum samples) for immunoreactivity-based preselection. (B) Preselected peptide library (14k spots/well, 105 serum samples). (C) Immune system metrics: **Z**-vector computed on ALL, TBC, or RRL libraries. (D) Predictive peptide selection: Elastic Net → Random Forest → Logistic Regression. Both pipelines evaluated by nested 5×5 cross-validation.

After removing the confounding effect of HIV co-infection, we detect a severity-dependent remodeling gradient and identify a three-peptide panel that detects smear-negative TB with AUC = 0.83 and specificity of 0.95. These findings demonstrate that high-density peptide arrays can function as topological sensors of antibody repertoire organization, enabling serological detection of chronic immune states beyond antigen-specific recognition.

## Results

### Study Cohort and Peptide Array

A total of 105 serum samples were profiled on high-density peptide microarrays containing 4,317 twelve-mer peptides derived from the *Mycobacterium tuberculosis* complex proteome (TBC; 4,241 protein-specific and 76 randomized internal controls), 2,087 ten-mer peptides from a resemblance-ranking library (RRL) designed to represent the human peptidome (26), and other peptides (overall 6,936 peptides; see Methods section). The RRL peptide panel was derived from a two-stage design (Figure 1A, B; see Methods section). The cohort comprised 58 TB-negative controls and 47 TB-positive patients (27 smear-positive/culture-positive [S+C+] and 20 smear-negative/culture-positive [S−C+]), recruited from South Africa, Peru, and Vietnam. 35 patients were HIV co-infected, 70 were HIV-negative.

### Immune Remodeling Is Detectable Through Distributional Metrics

Five immune state features (Shannon entropy, Gini coefficient, PC1, PC2, Mahalanobis distance) were computed from the full peptide intensity profile, forming the immune state vector **Z** (Figure 1C). Using **Z** as input for logistic regression (nested 5×5 CV), the remodeling classifier achieved AUC = 0.593 ± 0.154 on all peptides, 0.634 ± 0.159 on TBC peptides, and 0.615 ± 0.107 on RRL peptides in the full cohort (Table 1). The narrow range across libraries (ΔAUC = 0.041 between TBC and ALL) already suggests that the classification signal is not restricted to pathogen-derived peptides. The peptide-level classifier (EN→RF→LR, top 50) achieved AUC = 0.561 ± 0.114. All models showed moderate performance in the full cohort, reflecting the confounding effect of HIV co-infection on the antibody binding landscape, as demonstrated below.

**Table 1.** Model performance: Full cohort vs HIV-negative only.

| Model | Full ($n = 105$) | HIV- ($n = 70$) | ΔAUC |
| --- | --- | --- | --- |
| Remodeling (ALL, 6,936) | $0.593 \pm 0.154$ | **$0.728 \pm 0.086$** | +0.135 |
| Remodeling (TBC, 4,317) | $0.634 \pm 0.159$ | **$0.670 \pm 0.102$** | +0.036 |
| Remodeling (RRL, 2,087) | $0.615 \pm 0.107$ | **$0.691 \pm 0.131$** | +0.076 |
| Peptide LR (ALL, EN→RF→50) | $0.561 \pm 0.114$ | **$0.740 \pm 0.108$** | +0.179 |

### HIV Co-Infection Masks the TB Remodeling Signal

Exclusion of HIV-positive samples dramatically improved classification across all models and libraries (Table 1, Figure 2). For the remodeling classifier on all peptides, AUC increased from 0.593 to 0.728 (+0.135); for the peptide-level classifier, from 0.561 to 0.740 (+0.179) (Table 1). The improvement was not restricted to pathogen-derived peptides: on the host-proteome RRL library, remodeling AUC increased from 0.615 to 0.691 (+0.076), confirming that HIV masks a global immune signal rather than selectively affecting antigen-specific responses. In the Entropy– Gini phase space, group ellipses separated visibly after HIV exclusion (Cohen’s d: entropy 0.26 → 0.65; Gini 0.22 → 0.67; Figure 2, top vs bottom row). The TB-positive centroid shifted toward lower entropy and higher Gini values relative to TB-negative controls, and this shift was geometrically consistent across all three peptide libraries (Figure 2, columns). In the full cohort, HIV-positive samples (Figure 2, top row, X markers) occupied a dispersed region overlapping both TB-positive and TB-negative clouds, providing a geometric explanation for the reduced classification performance. This improvement was consistent across all three libraries.

**Figure 2.**
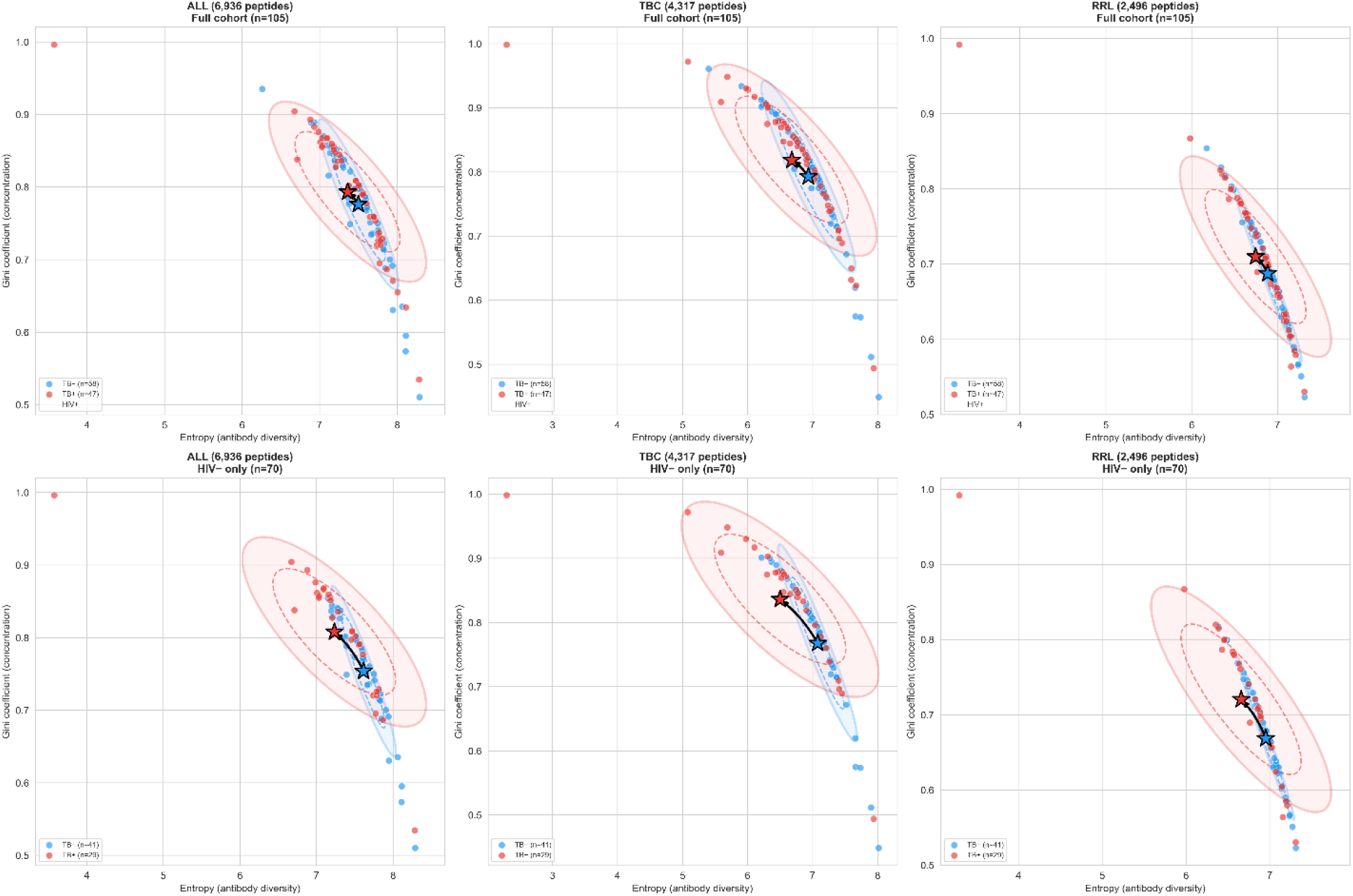
Entropy × Gini phase space with centroids and confidence ellipses. Top row: Full cohort (n = 105). Bottom row: HIV-negative only (n = 70). Columns: ALL, TBC, RRL peptide libraries. Stars = group centroids; dashed ellipses = 1σ; filled ellipses = 1.5σ; arrows = TB− → TB+ shift.

### Immune Remodeling Is Library-Independent

The immune state vector **Z**, computed independently from TBC, RRL, and ALL libraries, produced nearly identical classification performance (Table 1) and virtually identical trajectories in the Entropy–Gini phase space (Figure 2). In the HIV-negative subcohort, the remodeling AUC was 0.728 (ALL), 0.670 (TBC), and 0.691 (RRL) — a range of only 0.058 across libraries with fundamentally different biological origins. The effect size profiles confirmed the same pattern: entropy and Gini showed medium Cliff’s delta (|δ| > 0.32) consistently across all three libraries, while PC1 showed a library-dependent gradient with the largest effect on TBC (δ = −0.427) and the smallest on RRL (δ = −0.191) (Table 2). The direction of all five Z-features was concordant between TBC and RRL (5/5 agreement), and the rank order of effect sizes was strongly correlated (Spearman ρ = 0.900, p = 0.037). The near-equivalence of TBC and RRL demonstrates that TB infection induces a global redistribution of the antibody repertoire encompassing both anti-pathogen and anti-self reactivity.

**Table 2.** Effect sizes (Cliff’s delta) per **Z**-feature, HIV-negative only.

| Feature | ALL | TBC | RRL |
| --- | --- | --- | --- |
| Entropy | -0.435 | -0.450 | -0.353 |
| Gini | +0.422 | +0.428 | +0.326 |
| PC1 | -0.334 | -0.427 | -0.191 |
| PC2 | +0.229 | +0.041 | +0.230 |
| Mahalanobis | +0.201 | +0.118 | +0.159 |

### Two-Stage Feature Selection

With 6,936 features and only 105 samples (p >> n), direct feature ranking is unstable. Elastic Net pre-filtering (l1_ratio = 0.5, α = 0.001) consistently removed approximately two-thirds of peptides within each cross-validation training fold (ALL: 6,936 → 2,318; TBC: 4,317 → 1,844; RRL: 2,087 → 985), improving the feature-to-tree ratio from 0.07 to 0.22 (Supplementary Table S2). A sweep across panel sizes revealed that 3 peptides achieved AUC = 0.717 in the full cohort, outperforming 50 peptides (AUC = 0.561). In the HIV-negative subcohort, optimal performance was N = 20 (AUC = 0.739). This inverted relationship between panel size and performance reflects the high-dimensional noise inherent in peptide array data and provides empirical justification for the Elastic Net pre-filter (Supplementary Figure S1, Supplementary Table S3).

### Smear-Negative TB Detection

The analysis was stratified by TB severity to assess whether immune remodeling can detect the diagnostically most challenging group: smear-negative, culture-positive (S−C+) patients — those who carry TB but test negative by standard sputum microscopy (Figure 3).

**Figure 3.**
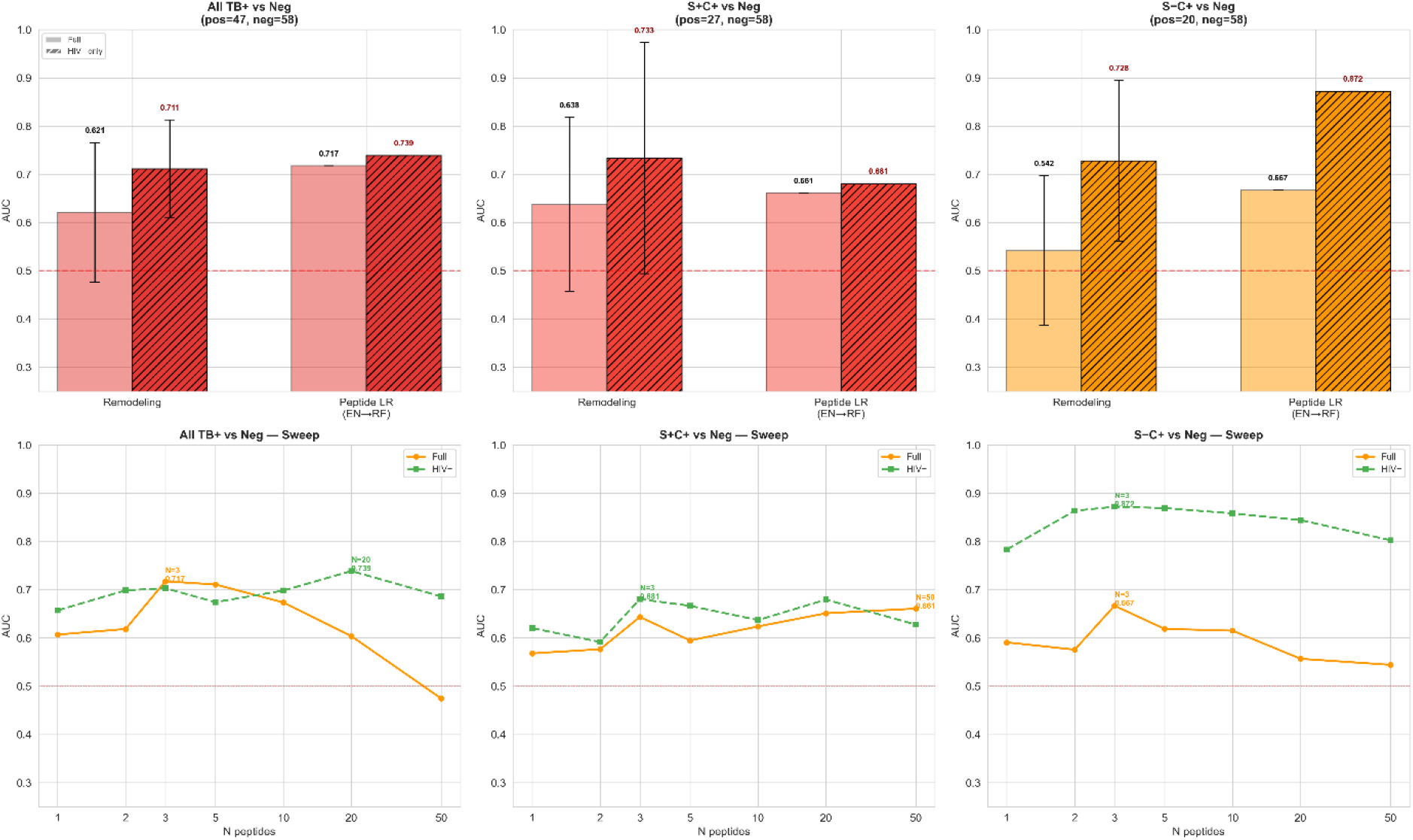
TB severity analysis: AUC bars and peptide number sweep curves for All TB+ / S+C+ / S−C+ vs Negative, Full cohort and HIV-negative only.

In the full cohort, smear-negative TB was undetectable by remodeling (AUC = 0.508, indistinguishable from chance). After HIV exclusion, AUC rose to 0.706 for remodeling and 0.831 [0.690–0.951] for the 3-peptide classifier — the largest HIV-exclusion effect (ΔAUC = +0.211). This result is visible in Figure 3 (bottom row, right panel): S−C+ classification jumps from chance level in the full cohort to the highest AUC observed in the entire study. The 3-peptide panel achieved specificity 0.951 [0.875–1.000] and sensitivity 0.643 [0.375–0.889] at the Youden-optimal threshold (Table 3). For comparison, smear-positive TB (S+C+) was detectable by remodeling alone (AUC = 0.700) but showed weaker peptide-level performance (AUC = 0.628), the opposite pattern to S−C+. This result suggests that different severity levels engage different aspects of the immune remodeling process (Figure 3, middle vs right columns).

**Table 3.** Classification by TB severity (HIV-negative only).

| Task | n<br>(pos/neg) | Remodeling AUC [95%<br>CI] | Peptide LR AUC [95%<br>CI] | Sens | Spec |
| --- | --- | --- | --- | --- | --- |
| All TB+ vs Neg | 70 (29/41) | 0.728 [0.524–0.796] | 0.740 | 0.448 | 0.902 |
| S+C+ vs Neg | 56 (15/41) | 0.700 [0.460–0.807] | 0.628 | — | — |
| S-C+ vs Neg | 55 (14/41) | **0.706 [0.522–0.867]** | **0.872 [0.690–0.951]** | **0.643** | **0.951** |

### Different Immune Signatures for Different Severity

The effect size analysis revealed that S+C+ and S−C+ disease are characterized by qualitatively different immune remodeling profiles in the HIV-negative subcohort (Table 4, Figure 4). S+C+ was detected through distributional metrics (entropy δ = −0.567, Gini δ = +0.545), consistent with broad immune activation and repertoire focusing in high-burden disease. S−C+ relied on structural features (PC1 δ = −0.521, PC2 δ = +0.364), suggesting a subtler reorganization that alters the principal axes of repertoire variation without substantially changing the overall diversity measures. In the Entropy–Gini projection (Figure 4, top row), the three groups arranged along a continuous trajectory Negative → S−C+ → S+C+, with S−C+ occupying an intermediate position — consistent with a dose-dependent remodeling process proportional to bacterial burden. The PC1 × PC2 projection (Figure 4, bottom row) revealed a distinct centroid geometry in which S−C+ patients separated from controls along a different axis than S+C+ patients, confirming that distributional and structural features capture complementary aspects of the severity gradient.

**Table 4.**
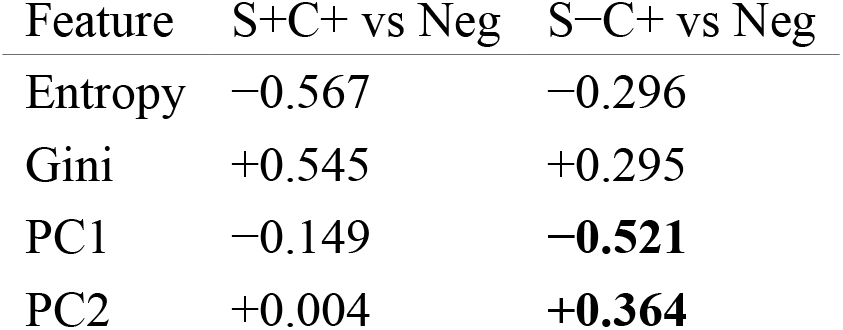
Cliff’s delta by severity (HIV-negative, ALL library).

**Figure 4.**
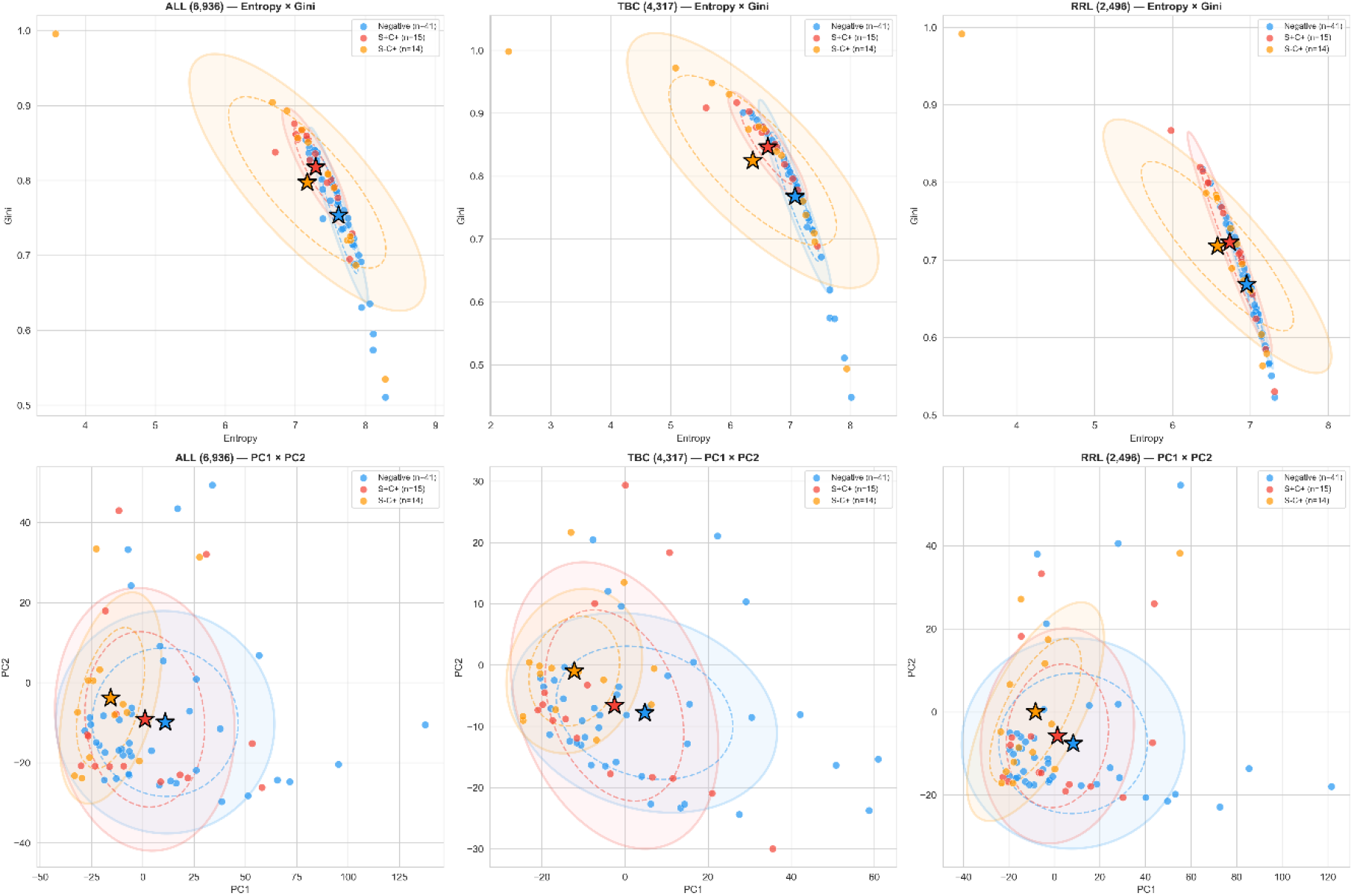
TB severity in phase space (HIV-negative only). Top row: Entropy × Gini. Bottom row: PC1 × PC2. Stars = centroids; ellipses = 1σ/1.5σ. Blue = Negative, Red = S+C+, Orange = S−C+.

### Diagnostic Peptides

#### Pathogen and Host-Proteome Converge

Analysis of the top peptides selected for S−C+ detection (HIV-negative) revealed a composition that directly reflects the dual nature of immune remodeling (Figure 5). 9 of the top 15 peptides originated from TBC antigens (Ag85A, PPE60, EchA3, GarA, Culp1, LprG, Mpt63), while 6 were RRL host-proteome peptides (Figure 5, left panel; red bars = TBC, blue bars = RRL). The TBC peptides mapped to established immunodominant *M. tuberculosis* proteins: Ag85A (the major secreted antigen used in vaccine development), and PPE60 (PE/PPE immune evasion family, highly expressed in extracellular vesicles) (1, 13, 27), EchA3 (28-31) (crotonase, a serodiagnostic candidate), GarA (13, 32, 33) (glycogen accumulation regulator, differentiates latent/acute TB in serodiagnosis), Culp1 (34) (cutinase, serodiagnostic antigen), LprG (28, 35-37) (conserved lipoprotein, serodiagnostic antigen) and Mpt63 (34, 37) (immunoprotective secreted protein, serodiagnostic antigen). The independent rediscovery of these known diagnostic targets provides external validation of the EN→RF selection pipeline. RRL peptides were enriched for hydrophobic residues (49% vs 38%) and alpha-helix motifs, suggesting that they sample antibody reactivity against structural host protein motifs. All 15 peptides showed reduced reactivity in S−C+ patients (all p < 0.001; Figure 5, right panel — all bars point left), consistent with attenuated antibody responses in paucibacillary disease across both pathogen-directed and host-directed targets (Supplementary Table S1). This parallel reduction across both compartments supports the model of a globally remodeled antibody repertoire in smear-negative TB.

**Figure 5.**
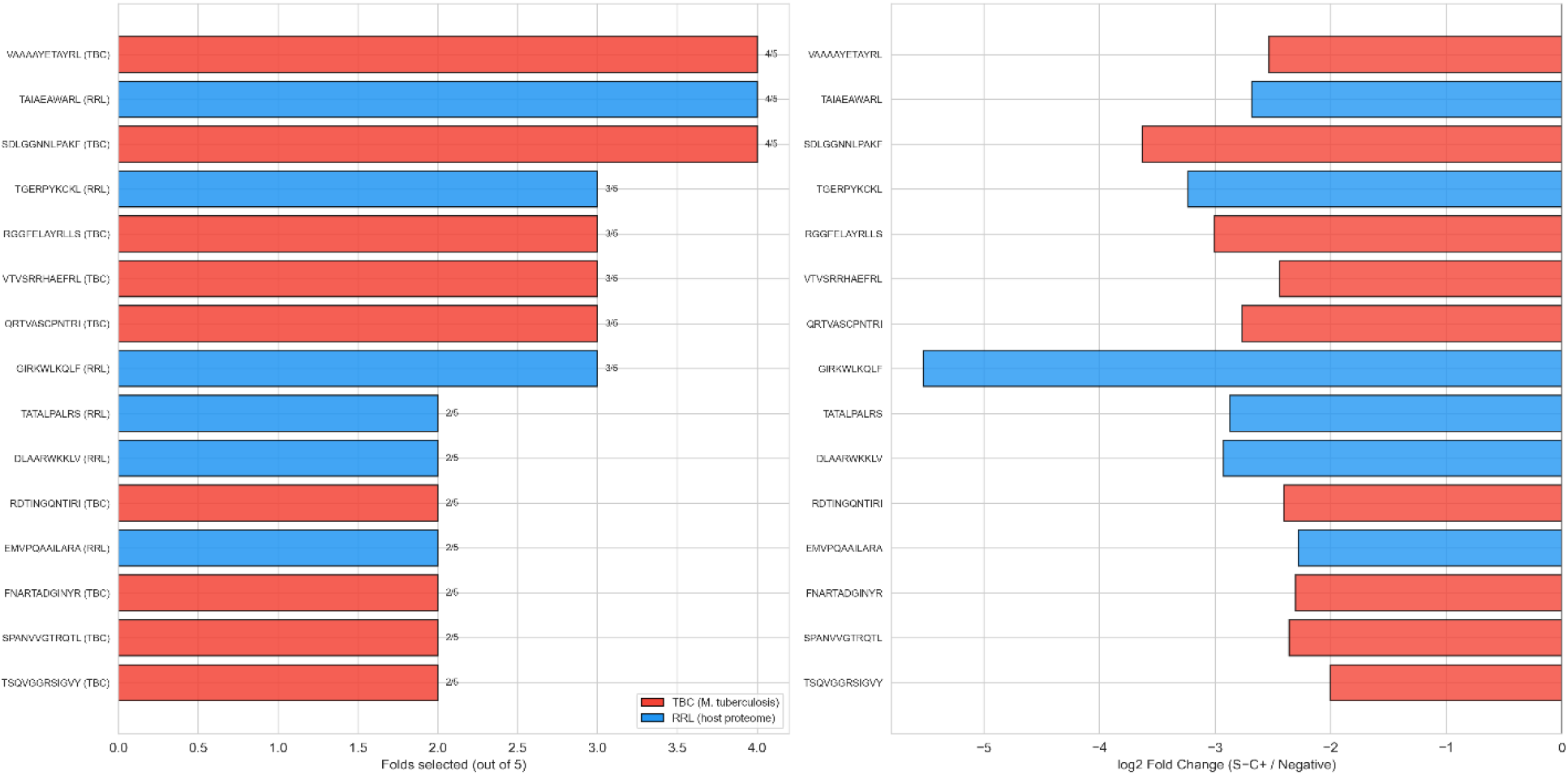
Top 15 peptides for S−C+ detection (HIV-negative). Left: Folds selected (red = TBC, blue = RRL). Right: log_2_ fold-change direction (all negative in S−C+).

### Remodeling and Peptide Classifiers: Same Biology, Different Resolution

We tested whether combining the 5 remodeling features (**Z**-vector) with the top EN→RF-selected peptides would improve classification. Across all tasks and cohorts, the combined model did not outperform the better individual model (Table 5). For all TB+ detection, the combined model (AUC = 0.650) performed worse than remodeling alone (0.711). For S+C+ detection, the combined model (0.706) was slightly below remodeling (0.733). For S−C+ the combined model was substantially worse (0.758 vs 0.872 for peptides alone — a loss of 0.114 AUC points). This pattern is consistent across all tasks: the combined model consistently falls between or below the two individual models, never exceeding the better one (Supplementary Figure S2). Both approaches measure the same immune perturbation at different analytical resolutions: the Z-vector summarizes the global distributional shape of the antibody binding landscape (macroscopic view), while individual peptides resolve specific antibody-epitope interactions (microscopic view). Because the **Z**-vector is computed from the same peptide intensities, combining both adds noise rather than new information.

**Table 5.**
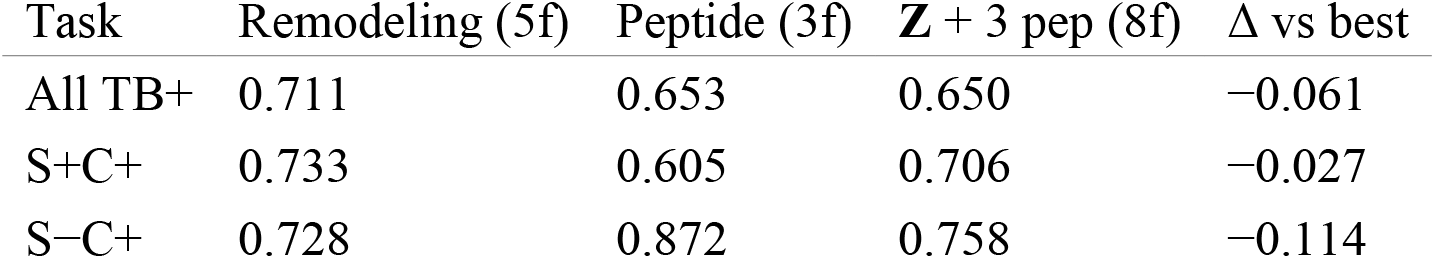
Combined model performance (HIV-negative).

| Task | Remodeling (5f) | Peptide (3f) | **Z** + 3 pep (8f) | $\Delta$ vs best |
| --- | --- | --- | --- | --- |
| All TB+ | 0.711 | 0.653 | 0.650 | −0.061 |
| S+C+ | 0.733 | 0.605 | 0.706 | −0.027 |
| S−C+ | 0.728 | 0.872 | 0.758 | −0.114 |

### Clinical Diagnostic Strategy for Smear-Negative TB

Although the remodeling and peptide classifiers capture the same underlying biology, their distinct operating characteristics enable a clinically meaningful two-stage diagnostic strategy for smear-negative TB in HIV-negative patients (Figure 6, Table 6). The remodeling score achieves high sensitivity (0.857) — it detects 12 of 14 smear-negative TB patients — suitable for screening; the 3-peptide panel achieves high specificity (0.951) — only 2 false positives in 41 negatives — suitable for confirmation (Figure 6). A parallel strategy (positive if either test is positive) missed 1 of 14 S−C+ cases in this cohort (sensitivity 0.949 assuming independence; NPV 0.952), at the cost of a higher false-positive rate (21/41) that would require subsequent culture confirmation — a trade-off acceptable in TB-endemic settings. Conversely, a sequential strategy (positive only if both tests agree) achieves specificity of 0.976 with only 1 false positive out of 41 negatives (Table 6).

**Table 6.** Two-stage diagnostic strategy (S−C+ vs Neg, HIV−, n = 55).

| Strategy | Sensitivity | Specificity | PPV | NPV | Missed | False+ |
| --- | --- | --- | --- | --- | --- | --- |
| Remodeling alone | **0.857** | 0.512 | 0.375 | 0.913 | 2/14 | 20/41 |
| 3-peptide alone | 0.643 | **0.951** | 0.818 | 0.886 | 5/14 | 2/41 |
| Sequential (AND) | 0.551 | **0.976** | **0.889** | 0.870 | 6/14 | **1/41** |
| Parallel (OR) | **0.949** | 0.487 | 0.382 | **0.952** | **1/14** | 21/41 |

**Figure 6.**
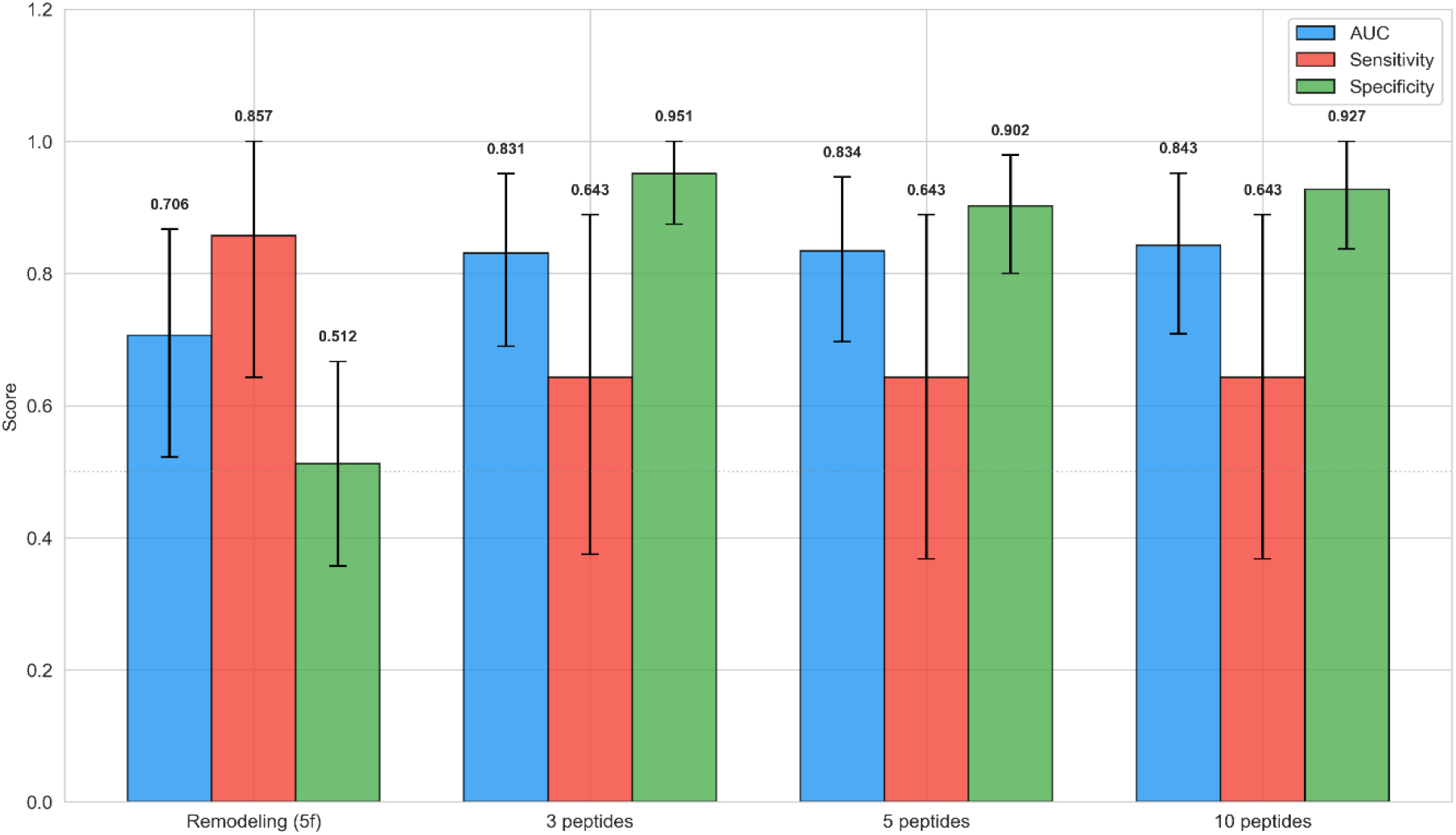
Sensitivity/Specificity with 95% Bootstrap CI for S−C+ vs Negative (HIV-negative, n = 55).

### Statistical Validation: Permutation Tests

To confirm that the observed classification performance is not an artifact of the analytical pipeline, we performed permutation testing: TB labels were randomly shuffled 200 times and the full remodeling pipeline was re-executed for each permutation (Figure 7, Table 7). The observed remodeling AUC exceeded the 95th percentile of the null distribution in both cohorts (Figure 7, red line clearly right of the gray null distribution). In the full cohort, the observed AUC (0.623) exceeded the null mean (0.496) with p = 0.035. In the HIV-negative subcohort, the result was stronger: observed AUC = 0.703, null mean = 0.503, p = 0.005 (Figure 7, right panel). The null distributions centered at approximately 0.50 in both cohorts, confirming that the nested cross-validation procedure is unbiased and does not inflate performance estimates.

**Table 7.**
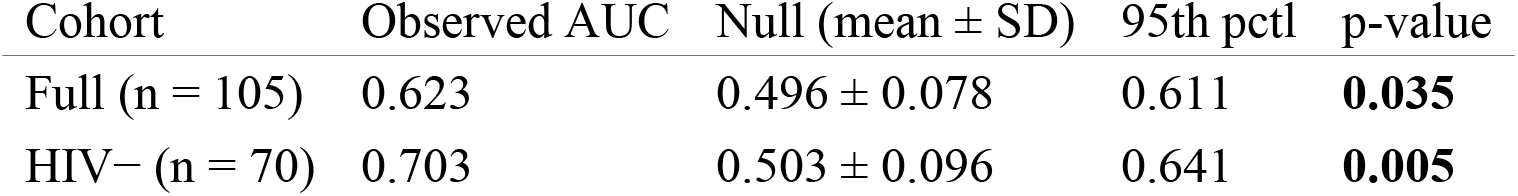
Permutation test results.

| Cohort | Observed AUC | Null (mean ± SD) | 95th pctl | p-value |
| --- | --- | --- | --- | --- |
| Full (n = 105) | 0.623 | 0.496 ± 0.078 | 0.611 | **0.035** |
| HIV- (n = 70) | 0.703 | 0.503 ± 0.096 | 0.641 | **0.005** |

**Figure 7.**
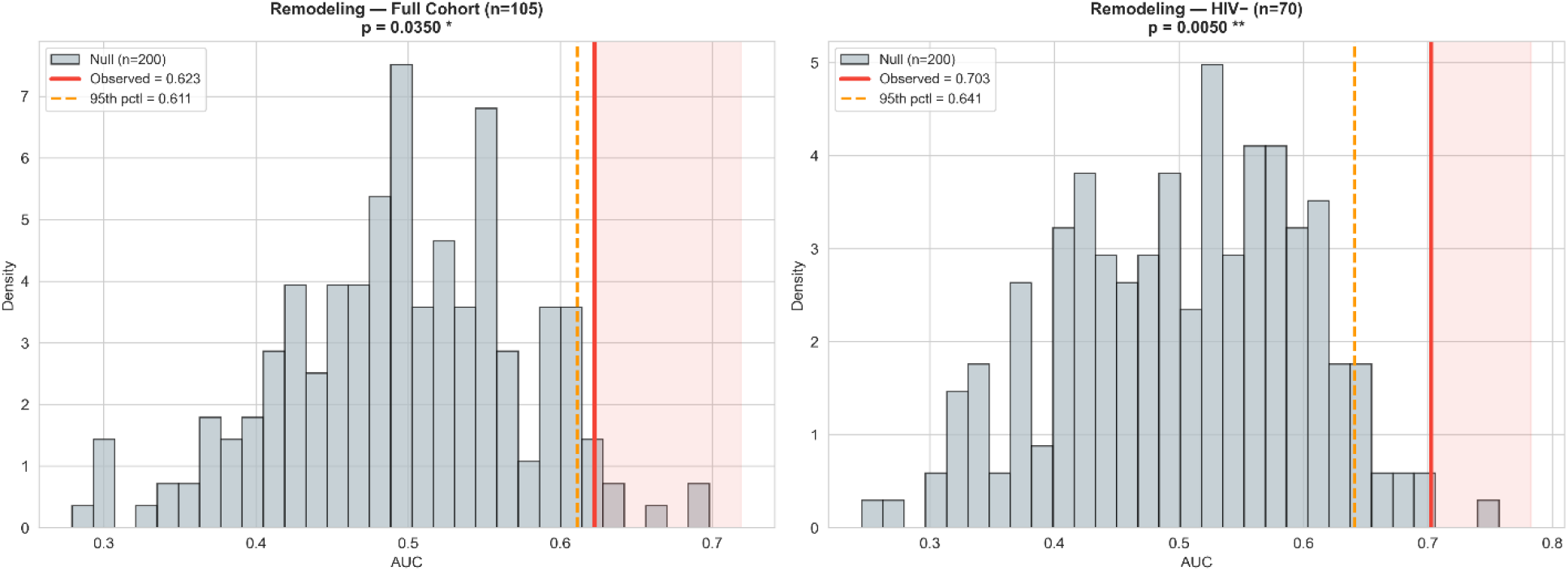
Permutation test histograms (200 iterations). Left: Full cohort. Right: HIV-negative. Red line = observed AUC; gray = null distribution.

### Cross-Population Generalization

To assess whether the immune remodeling signal generalizes across geographically distinct populations, we performed leave-one-country-out (LOCO) cross-validation: models were trained on samples from two countries and tested on the held-out country (Figure 8, Table 8).

**Table 8.**
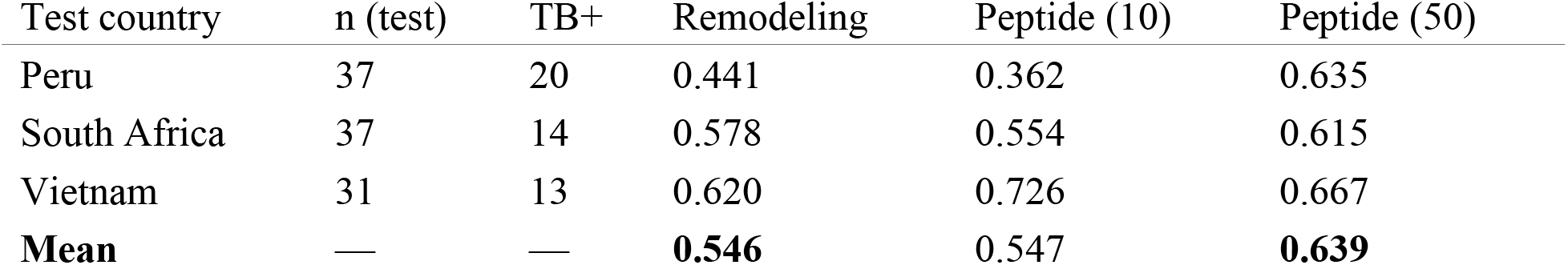
Leave-one-country-out results (full cohort).

**Figure 8.**
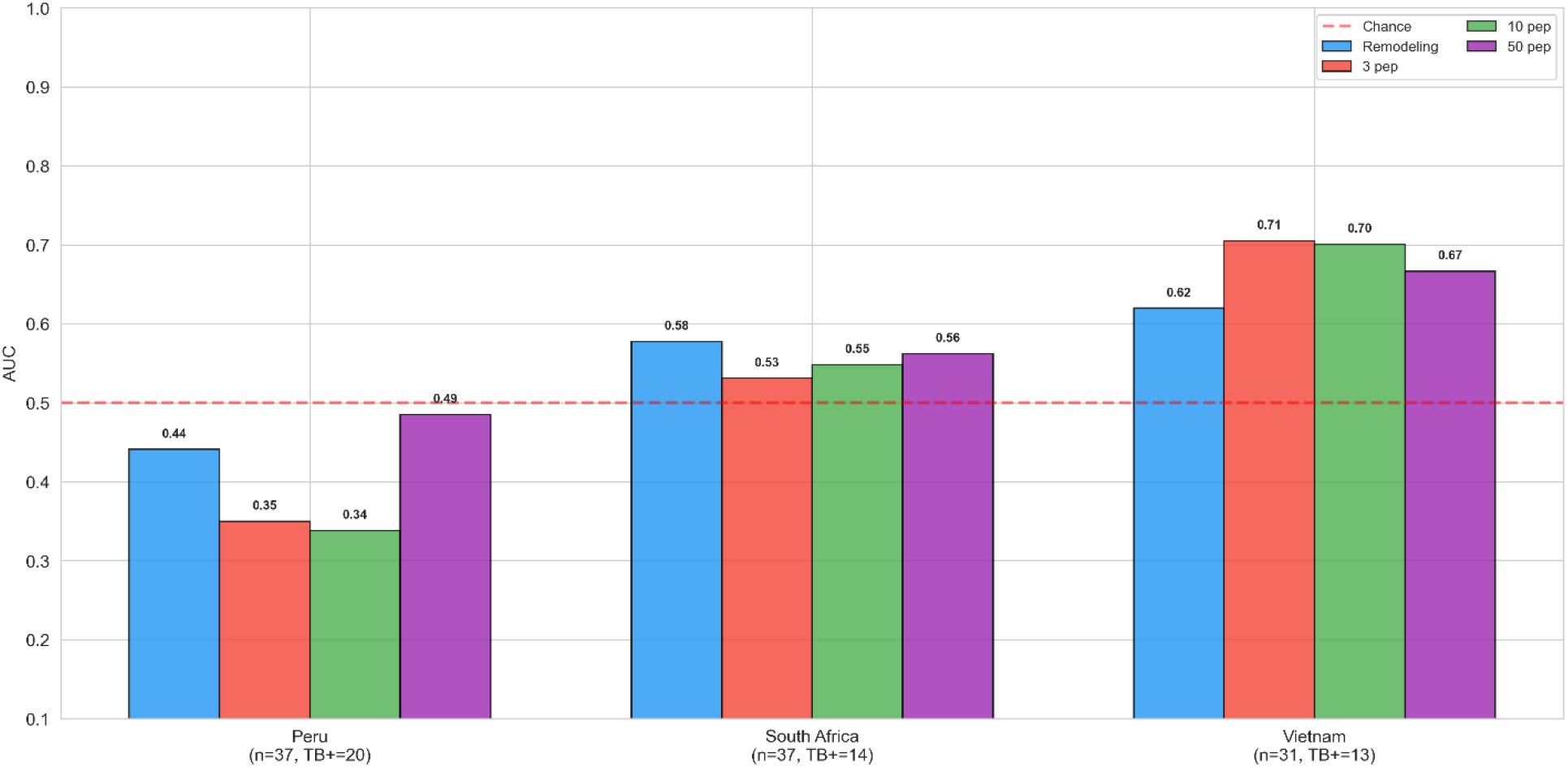
Leave-one-country-out validation: AUC by test country for Remodeling and Peptide models.

The remodeling classifier showed variable cross-population transfer (mean LOCO AUC = 0.546), with Vietnam showing the strongest generalization (AUC = 0.620), South Africa intermediate performance (AUC = 0.578), and Peru below-chance performance (AUC = 0.441) — consistent with the larger TB-positive fraction at this recruitment site (20/37). The peptide-level classifier with 50 features showed stronger and more uniform cross-population generalization (mean LOCO AUC = 0.639), maintaining above-chance performance in all three test populations (Figure 8). The performance gap between within-cohort cross-validation (AUC = 0.623) and LOCO (AUC = 0.546) reflects the population heterogeneity inherent in a three-continent cohort and provides a benchmark for future multicenter validation.

The LOCO results suggest that cross-population transfer of the remodeling signal is sensitive to cohort composition and sample size, whereas the peptide-level classifier generalizes more robustly across recruitment sites. The S−C+ analysis is based on 14 HIV-negative cases; validation in a larger, multicenter cohort is essential. Nevertheless, the finding that smear-negative TB becomes detectable only after HIV exclusion — confirmed by permutation testing (p = 0.005) — suggests the underlying immune signal is genuine.

## Discussion

Using high-density peptide arrays with two complementary libraries, pathogen-derived (TBC) and host-proteome-derived (RRL), we observed that a distributional classifier based on five immune state features achieved virtually identical classification performance on both libraries. This convergence — in which the remodeling classifier achieved comparable performance on both libraries while individual peptide selection was dominated by TBC-derived sequences (Figure 5) — demonstrates that TB infection induces a measurable restructuring of the antibody binding landscape that extends beyond pathogen-specific epitope recognition(1). The signal is subtle in the full cohort (AUC ∼0.60) but becomes clearly detectable after removing the confounding effect of HIV co-infection (AUC 0.73, permutation p = 0.005), and unexpectedly enables detection of smear-negative TB, the form of disease most resistant to conventional diagnostics. (Table 1, Table 7)

The equivalence of classification performance on TBC and RRL peptides is the central finding of this work. The RRL is not a random peptide collection; it was constructed by a resemblance-ranking algorithm to maximally represent the k-mer fragment diversity of the entire human proteome(26). Each RRL peptide approximates a sequence fragment occurring in human proteins. When TB-associated diagnostic information is detectable on these host-proteome peptides at the same accuracy as on *M. tuberculosis* antigens (Table 1, Table 2) this implies that TB infection measurably alters the antibody reactivity against self-derived epitopes. This observation extends the concept of immune remodeling beyond the pathogen-specific response: TB does not merely stimulate anti-mycobacterial antibodies but reshapes the self-reactive component of the antibody repertoire. Such reshaping is biologically plausible given the prolonged germinal center reactions (6, 7), sustained polyclonal B-cell activation, and affinity maturation that characterize chronic TB infection (13). Affinity-matured antibodies frequently exhibit degrees of polyreactivity (11), and clonal expansion of dominant B-cell populations can redistribute binding across broad regions of peptide sequence space, including host-like sequences (9, 12). The RRL library serves as a statistical sensor for such redistribution.

To provide a geometric interpretation of these findings, we mapped individual sera into an immune phase space defined by the five distributional coordinates of the Z-vector. This representation treats each serum sample as a point in a continuous immune state manifold, where disease is detected as displacement within this manifold. Similar concepts have emerged in immune repertoire analysis, where antibody and T-cell repertoires are studied as complex systems whose structure encodes information about immune history (8, 10). The Entropy–Gini projection revealed that TB-positive samples are displaced toward lower entropy and higher Gini coefficients, consistent with a narrowing and concentration of the antibody repertoire (Figure 2). Entropy has previously been proposed as an indicator of immune health status in the context of immunosignaturing (23), and our findings extend this concept to a multivariate phase-space framework. This displacement was virtually identical on TBC and RRL peptides, providing a geometric confirmation of the library-independence (Figure 2, Table 2). The phase-space framework is particularly suited to chronic infections such as tuberculosis, where prolonged antigen persistence drives progressive repertoire focusing over months to years (7). Unlike acute infections — where immune perturbation is transient and the antibody landscape rapidly returns to baseline — chronic TB produces a stable shift in repertoire topology that persists as a quasi-stationary state. Whether this approach can detect immune perturbations in acute infections remains an open question for future investigation.

The effect of HIV co-infection provides both a biological insight and a practical constraint. HIV-positive samples formed a dispersed cloud in the Entropy–Gini phase space that partially overlapped with the TB-positive region (Figure 2, top row). This overlap is consistent with both conditions independently driving repertoire focusing — TB through chronic antigen-specific stimulation(13), HIV through polyclonal B-cell activation, hypergammaglobulinemia, and contraction of functional B-cell subsets(14-16). The four-group centroid ordering (TB−/HIV− → TB−/HIV+ ≈ TB+/HIV− → TB+/HIV+) quantifies this geometric convergence and explains why HIV exclusion uniformly improved classification (Figure 2, Table 1). This finding has direct implications for diagnostic deployment: peptide-array-based immune remodeling assays may perform best in HIV-negative populations or require HIV-stratified interpretation algorithms.

Stratification by TB severity revealed an unexpected gradient in the immune phase space. Smear-positive patients (S+C+) showed the strongest distributional shift, driven primarily by entropy and Gini coefficient. Smear-negative patients (S−C+) exhibited a qualitatively different signature dominated by structural features (PC1, PC2) (Table 4, Figure 4), suggesting a subtler reorganization that alters the principal axes of repertoire variation without substantially changing the overall diversity. The three groups arranged along a continuous trajectory — Negative → S−C+ → S+C+ — consistent with a dose-dependent remodeling process proportional to bacterial burden. (Table 4, Figure 4)

The detection of smear-negative TB is the most clinically relevant finding. In the full cohort, S−C+ classification was at chance level (AUC = 0.508). After HIV exclusion, the peptide-level classifier achieved AUC = 0.831 with only three peptides (Table 3, Figure 3). The selected peptides included both TBC antigens (Ag85A, PPE60 (1, 13), EchA3 (28-31), GarA(13, 32, 33), Culp1 (34), LprG (28, 35-37), Mpt63(34, 37)) and RRL host-proteome peptides, and all showed reduced reactivity in S−C+ patients. This parallel attenuation across pathogen-directed and host-directed targets is consistent with a globally dampened immune remodeling signature in paucibacillary disease. (Figure 5)

An informative negative result emerged from the combined model analysis. Concatenating **Z**-vector features with top peptide intensities did not improve classification (Table 5). This confirms that both approaches measure the same underlying immune perturbation at different analytical resolutions: the **Z**-vector summarizes the global shape of the binding distribution (macroscopic), while individual peptides resolve specific antibody–epitope interactions (microscopic). This convergence of two methodologically independent approaches strengthens the evidence for a coherent, system-wide immune remodeling process.

Despite measuring the same underlying biology, the two classifiers exhibit complementary operating characteristics enabling a two-stage diagnostic strategy. The remodeling score achieves high sensitivity (0.857) suitable for screening; the three-peptide panel achieves high specificity (0.951) suitable for confirmation. A parallel deployment reduces missed smear-negative TB cases to 1 in 14 (sensitivity 0.949), while a sequential strategy achieves specificity of 0.976 (Figure 6, Table 6). These trade-offs are directly relevant to TB-endemic settings, where missed cases drive ongoing transmission.

Several limitations should be considered. The cohort comprises 105 individuals, and the smear-negative analysis is based on 14 HIV-negative S−C+ cases. While permutation tests (38) confirm the signal is genuine (p = 0.005) (Figure 7, Table 7), independent validation in a larger cohort is essential. Leave-one-country-out validation showed moderate cross-population generalization (mean LOCO AUC = 0.546) (Figure 8, Table 8), reflecting population heterogeneity. The study is cross-sectional; longitudinal studies could determine whether entropy-based metrics track disease progression or treatment response. Sensitivity and specificity were estimated at a post-hoc threshold and should be considered optimistic until prospectively validated.

These limitations notwithstanding, the present findings suggest a conceptual reorientation of peptide-array-based serology. Traditionally, peptide arrays serve as platforms for epitope mapping (17, 18, 20). Immunosignaturing studies have demonstrated classification from collective binding patterns (21, 22), and phage immunoprecipitation approaches have enabled comprehensive profiling (24, 25). Our results extend these observations by demonstrating that the diagnostic signal on host-proteome peptides is equivalent to that on pathogen-derived peptides, arising from distributional repertoire restructuring. High-density peptide arrays can function as instruments for measuring the topology of antibody repertoires. The phase-space framework, in which individual sera are mapped into a low-dimensional immune state manifold and disease is detected as geometric displacement, may provide a general analytical approach for high-dimensional serological data across infectious, autoimmune, and inflammatory conditions, complementing existing systems serology approaches (2-5, 39).

## Methods

### Ethics Statement

Serum samples were obtained from the biobank of the Foundation for Innovative New Diagnostics (FIND, Geneva, Switzerland), a WHO collaborating centre for laboratory strengthening and diagnostic technology evaluation.

### Study Cohort

The biobank provided clinically annotated specimens from 105 individuals across three recruitment sites: South Africa (n = 37), Peru (n = 37), and Vietnam (n = 31). Clinical annotations included microbiologically confirmed TB status, HIV serostatus, sputum smear microscopy results, culture results, age, gender, and country of origin. The cohort included 47 individuals with confirmed active TB and 58 TB-negative controls. HIV co-infection status was determined for all participants: 35 were HIV-seropositive and 70 were HIV-seronegative. Among TB-positive patients, 27 were smear-positive/culture-positive (S+C+) and 20 were smear-negative/culture-positive (S−C+). TB diagnosis was based on microbiological confirmation (sputum smear microscopy and/or culture). TB-negative controls had no clinical, radiological, or microbiological evidence of active or latent TB.

### Peptide Array Design

The RRL peptide panel was derived from a two-stage design (Figure 1A,B). In the first stage, the discovery library with ∼35,000 peptides in duplicate (∼70,000 spots per well), were screened with sera from 11 individuals (6 TB+, 5 TB−). The discovery peptide library contained 10-mer peptides from resemblance-ranking peptide library (RRL) with the highest resemblance ranking (26). Based on the signals from the first screen (RRL peptides were ranked by overall binding intensity across both TB-positive and TB-negative samples), the number of RRL peptides was reduced to 2,087 peptides; Of the 11 discovery samples, 10 were included in the main cohort. Because selection was by immunoreactivity rather than disease discrimination, this does not introduce classification leakage.

The final peptide microarray comprised 6,936 peptides from three main sources. The largest group consisted of 4,317 twelve-mer peptides derived from the *Mycobacterium tuberculosis* complex (TBC), with the exception of six eight-mer peptides. Of these, 4,241 were protein-specific sequences — 4,155 genomic peptides from the *M. tuberculosis* reference strain H37Rv and 86 additional known epitopes retrieved from the Immune Epitope Database (IEDB) — along with 76 randomized copies serving as internal negative controls. The second component was the Resemblance-Ranking Library (RRL), contributing 2,087 ten-mer peptides designed to represent the human peptidome through a resemblance-ranking approach. The remaining 532 peptides originated from other sources and were all twelve-mers. These included both protein-specific sequences and their randomized counterparts: HIV (46 + 5), Epstein–Barr virus (10 + 5), human coronavirus (10 + 4), SARS-CoV-2 (10 + 2), influenza A (8 + 2), poliovirus (9 + 1), measles virus (5 + 2), and human GPCR-derived peptides (2 + 1). An additional 410 twelve-mer peptides derived from human proteome sequences were included as host-derived reference controls. In total, the array contained 4,341 protein-specific peptides, 98 randomized internal negative controls, 2,087 resemblance-ranking library peptides, and 410 human-derived reference peptides. None of the randomized control peptides appeared among the top discriminating features in any analysis.

### Microarray Incubation and Data Acquisition

Peptide libraries were synthesized on glass slides (25 × 75 mm) using Axxelera-based peptide array technology (Karlsruhe, Germany). Arrays in six-window chambers were hydrated (500 µL PBS-T, 10 min). Serum (1:400 in PBS-T + 10% Rockland Blocking Buffer MB-070; 400 µL) was incubated 2 h at RT, 120 rpm, in the dark. Washing: 4× PBS-T, then 2× 10-min incubation washes (180 rpm). Detection: goat anti-human IgG (1 µg/mL; Biozol #109-605-008), 45 min, 120 rpm. Slides were rinsed with ddH_2_O, dried, stored in argon at 4 °C. Scanning: Innoscan 1100 AL (Innopsys), 2 µm resolution, PMT 4, 635 nm. Intensities quantified by MAPIX software. The resulting 105 × 6,936 matrix was used for all analyses.

### Nested Cross-Validation

All classification used nested CV(40, 41): outer stratified 5-fold for performance; inner stratified 5-fold for hyperparameter tuning. All transformations (scaling, PCA, EN, RF, Mahalanobis) fitted on training data only. Seeds fixed (RANDOM_STATE = 42).

### Immune Remodeling Classifier

A five-dimensional **Z**-vector was computed per training fold: (i) Shannon entropy; (ii) Gini coefficient; (iii–iv) PC1, PC2 from PCA on z-scored intensities; (v) Mahalanobis distance from TB-negative centroid (regularization λ = 10^−3^). L2-regularized logistic regression with C tuned via inner CV over {0.001, 0.01, 0.1, 1, 10, 100}.

### Peptide-Level Classifier with Elastic Net Pre-Filter

Within each training fold: (i) Elastic Net (SGDClassifier, log loss, l1_ratio = 0.5, α = 0.001, balanced weights, 5,000 iterations) reduced features from 6,936 to ∼2,300 (33.4%); (ii) Random Forest (500 trees, balanced weights) ranked survivors; (iii) top N peptides classified by L2-logistic regression. Sweep: N = 1–200.

### HIV Stratification and TB Severity

Both classifiers re-evaluated on HIV-negative subset (n = 70). Severity stratification: All TB+ vs Neg, S+C+ vs Neg, S−C+ vs Neg. Combined classifier (**Z** + peptides) tested for complementarity.

### Clinical Performance

Sensitivity and specificity at Youden-optimal threshold from pooled out-of-fold predictions. Bootstrap 95% CI (1,000 iterations). Sequential (AND) and parallel (OR) strategies evaluated. Threshold estimated post hoc; values are optimistic estimates.

### Effect Sizes

Cliff’s delta (42) within training folds, averaged across folds. Thresholds: |δ| < 0.15 negligible, 0.15–0.33 small, 0.33–0.47 medium, > 0.47 large (43). Profile similarity: Spearman ρ between TBC and RRL vectors.

### Permutation Testing

200 label shuffles, each with full nested CV. p = (b + 1)/(N + 1) per Phipson & Smyth (38).

### Leave-One-Country-Out Validation

Train on 2 countries, test on held-out country. Both classifiers evaluated with N = 3, 5, 10, 50.

### Software

Python 3.11; scikit-learn v1.3, NumPy v1.24, SciPy v1.11, pandas v2.0, matplotlib v3.7. All seeds = 42. Code available as Jupyter notebooks.

## Supporting information

experimental data

readme to use code

## Acknowledgements

The authors acknowledge financial support from the Ministerium für Wirtschaft, Arbeit und Tourismus Baden-Württemberg through the Invest BW innovation programme.

## Author Contributions

A.N.-M., N.S.-M., S.B. and A.D. conceived the project. A.N.-M and A.D. supervised the project. A.N.-M. developed the computational methodology and implemented the analysis pipeline. D.S. performed data preprocessing and statistical validation. C.v B.-K. and R.P. assisted with peptide chip design, sample incubation, and data interpretation. J.M. and H.B. contributed to the design of peptide libraries. J.G. performed all experimental work. All authors contributed to the interpretation of results. A.N.-M. and A.D. wrote the manuscript with input from all authors.

## Data Availability

The peptide intensity matrix and sample metadata are available from the corresponding authors upon reasonable request. Source data for all figures are provided with this paper.

## Code Availability

Complete analysis code is available as Jupyter notebooks in the Supplementary Information.

