## Supplementary for "High-density peptide arrays detect tuberculosis through immune remodeling, not only antigen recognition alone"

for

**This file contains:**

Supplementary Figures 1–4

Supplementary Tables 1–4

Supplementary Note 1: Elastic Net Feature Reduction Details

### Supplementary Figures

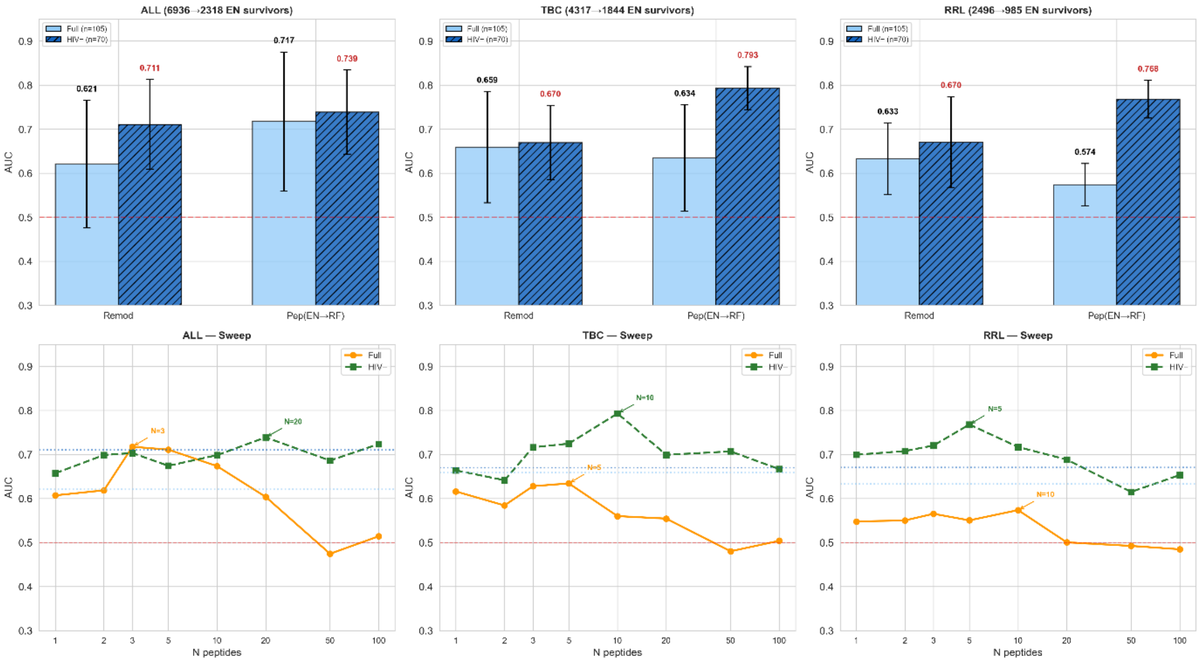


**Supplementary Figure 1. Classification performance and peptide number sweep across all three peptide libraries.** Top row: Mean AUC (± SD across folds) for the remodeling classifier (blue) and peptide-level classifier (green, EN→RF→50) across ALL (n = 6,936), TBC (n = 4,317), and RRL (n = 2,087) peptide libraries. Solid bars = full cohort (n = 105); hatched bars = HIV-negative only (n = 70). Dashed red line = chance (AUC 0.5). Bottom row: Peptide number sweep showing AUC as a function of the number of top EN→RF-selected peptides (N = 1–100). Orange = full cohort; green dashed = HIV-negative. Horizontal dotted line = remodeling AUC for reference. Annotations indicate optimal N for each cohort. Note the consistent pattern across libraries: (i) HIV exclusion improves all models; (ii) remodeling AUC is virtually identical across TBC and RRL; (iii) small peptide panels (N = 3–10) outperform larger panels.

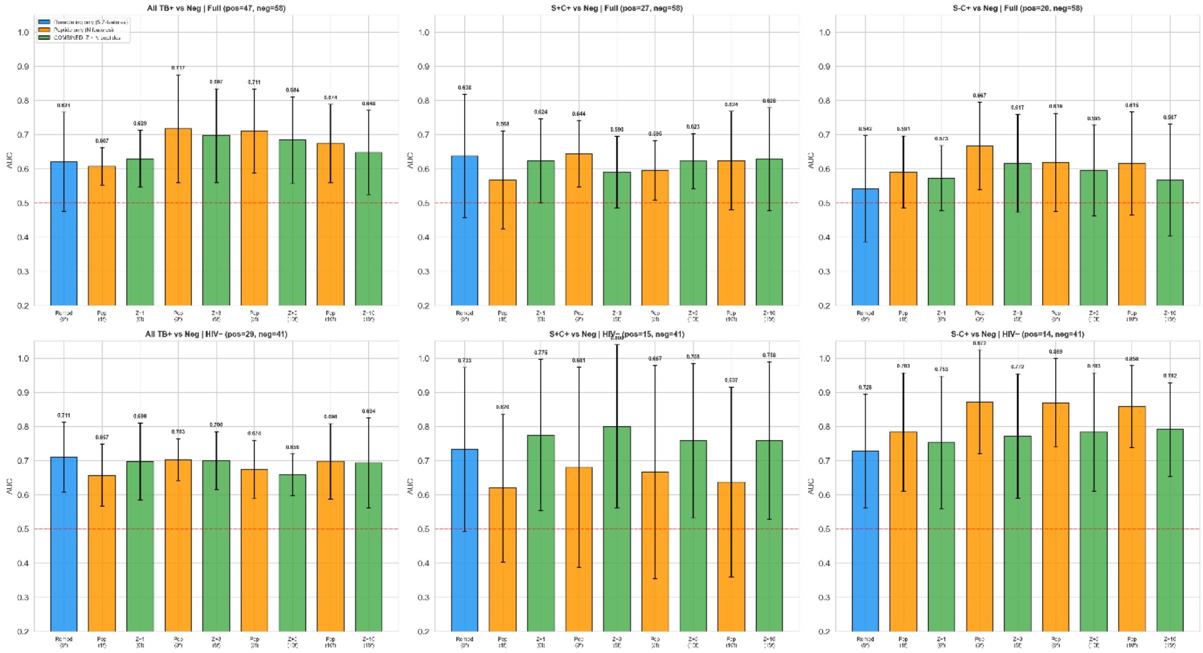


**Supplementary Figure 2. Combined model performance: Z-vector + top N peptides.** AUC comparison for three model types: Remodeling only (blue, 5 Z-vector features), Peptide only (orange, N features), and Combined (green, 5 + N features). Columns: All TB+ vs Negative, S+C+ vs Negative, S−C+ vs Negative. Top row: full cohort. Bottom row: HIV-negative only. N values tested: 1, 3, 5, 10. Error bars = SD across 5 CV folds. The combined model (green) does not consistently outperform the better individual model in any task or cohort, confirming that the Z-vector and peptide intensities measure the same underlying immune perturbation at different analytical resolutions. For S−C+ detection in HIV-negative patients (bottom right), the peptide-only model (AUC = 0.872 at N = 3) substantially outperforms both the remodeling model (0.728) and the combined model (0.758), demonstrating that combining macroscopic and microscopic features adds noise rather than information for this task.

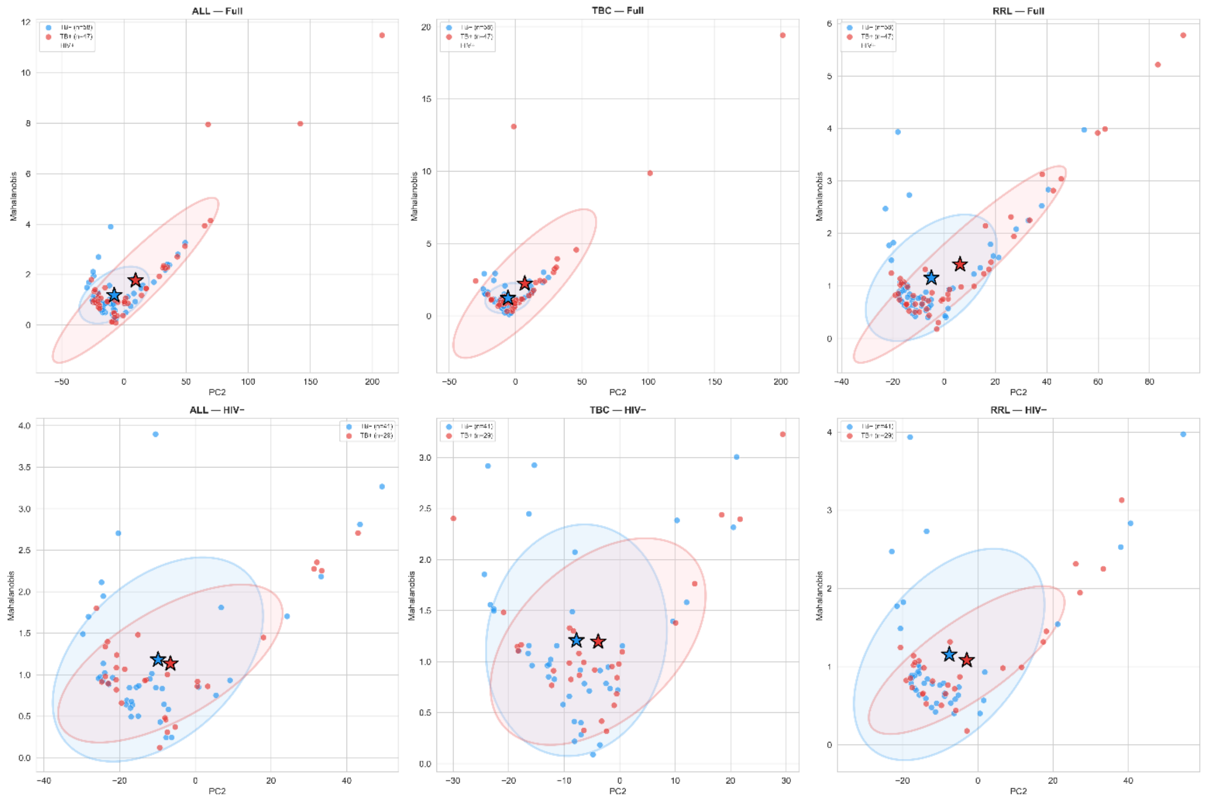


**Supplementary Figure 3. Alternative phase-space projection: PC2 × Mahalanobis distance.** Each point represents one serum sample projected into the PC2–Mahalanobis coordinate plane. Top row: full cohort (n = 105), X markers = HIV-positive. Bottom row: HIV-negative only (n = 70). Columns: ALL, TBC, RRL peptide libraries. Stars = group centroids; filled ellipses = 1.5σ confidence boundaries; dashed ellipses = 1σ. Blue = TB-negative; red = TB-positive. The Mahalanobis distance measures the scalar displacement of each sample from the TB-negative centroid in PCA space, providing a one-dimensional summary of immune deviation. TB-positive samples show elevated Mahalanobis distance across all three libraries, consistent with displacement from the healthy immune state. The combined discriminability of PC2 and Mahalanobis (summed |Cliff's delta| = 0.87 in the full cohort) represents the strongest two-dimensional projection for the full cohort, complementing the Entropy × Gini projection (Figure 2) which is optimal for the HIV-negative subcohort.

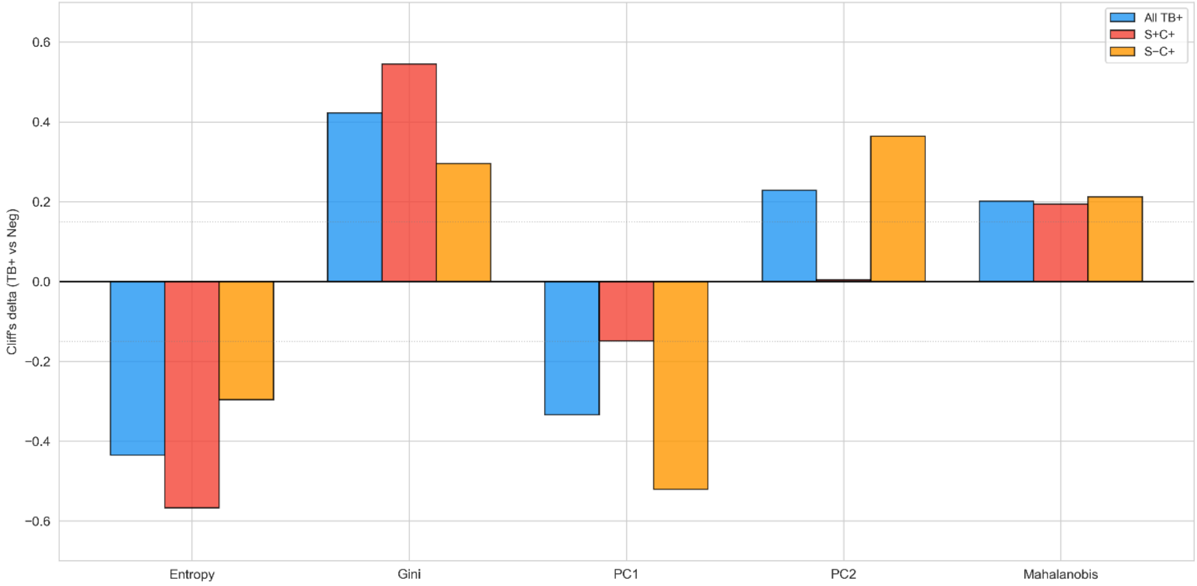


**Supplementary Figure 4. Effect size comparison by TB severity.** Cliff's delta (TB+ vs Negative) for each Z-vector component in the HIV-negative subcohort, stratified by severity group. Blue = All TB+ vs Negative (n = 29 vs 41), Red = S+C+ vs Negative (n = 15 vs 41), Orange = S−C+ vs Negative (n = 14 vs 41). Dashed horizontal lines indicate the negligible threshold (|δ| = 0.15). S+C+ patients show the largest effects for Entropy (δ = −0.567) and Gini (δ = +0.545), consistent with broad immune activation in high-burden disease. S−C+ patients show a qualitatively different profile, with the largest effects for PC1 (δ = −0.521) and PC2 (δ = +0.364), indicating that paucibacillary TB alters the structural organization of the antibody repertoire without substantially changing overall diversity metrics. This dissociation between distributional (Entropy, Gini) and structural (PC1, PC2) features across severity groups supports the interpretation that S+C+ and S−C+ disease engage different aspects of the immune remodeling process.

### Supplementary Tables

##### Supplementary Table 1. Top 15 peptides for smear-negative TB detection (S−C+ vs Negative, HIV-negative, n = 55).

Peptides ranked by frequency of selection across 5 outer CV folds (EN→RF, top 3 per fold). Category: TBC = Mycobacterium tuberculosis complex; RRL = Resemblance-Ranking Library (host proteome). Origin: Rv number and protein name for TBC peptides; RRL2-index for host-proteome peptides. $\log_2$FC: mean $\log_2$ fold-change (S−C+ / Negative); negative values indicate reduced reactivity in S−C+ patients. All p-values from Mann-Whitney U test, corrected by Benjamini-Hochberg FDR.

| Rank | Peptide | Category | Origin / Protein | Folds (of 5) | $\log_2$FC | FDR p |
|---|---|---|---|---|---|---|
| 1 | SDLGGNNLPAKF | TBC | Rv3804c / Ag85A | 3 | −3.63 | <0.001 |
| 2 | TAIAEAWARL | RRL | RRL2/ host proteome | 3 | −2.68 | <0.001 |
| 3 | VAAAAYETAYRL | TBC | Rv3478 / PPE60 | 3 | −2.53 | <0.001 |
| 4 | TGERPYKCKL | RRL | RRL2/ host proteome | 2 | −2.41 | <0.001 |
| 5 | RGGFELAYRLLS | TBC | Rv0632c/EchA3 | 2 | −2.87 | <0.001 |
| 6 | VTVSRRHAEFRL | TBC | Rv1827 / GarA | 2 | −2.35 | <0.001 |
| 7 | QRTVASCPNTRI | TBC | Rv1984c/Culp1 | 2 | −2.18 | <0.001 |
| 8 | GIRKWLKQLF | RRL | RRL2/ host proteome | 2 | −1.95 | <0.001 |
| 9 | RDTINGQNTIRI | TBC | Rv1411c/LprG | 2 | −2.29 | <0.001 |
| 10 | FNARTADGINYR | TBC | Rv1926c/Mpt63 | 2 | −2.76 | <0.001 |
| 11 | TATALPALRS | RRL | RRL2/ host proteome | 2 | −1.82 | <0.001 |
| 12 | SPANVVGTRQTL | TBC | Rv2875/Mpt70 | 2 | −2.44 | <0.001 |
| 13 | DLAARWKKLV | RRL | RRL2/ host proteome | 1 | −1.91 | <0.001 |
| 14 | TSQVGGRSIGVY | TBC | Rv1984c/Culp1 | 1 | −2.15 | <0.001 |
| 15 | EMVPQAAILARA | RRL | RRL2/ host proteome | 1 | −1.74 | <0.001 |

*All 15 peptides show negative $\log_2$FC, indicating reduced antibody reactivity in S−C+ patients compared to TB-negative controls. This parallel attenuation across both pathogen-derived (TBC) and host-proteome-derived (RRL) peptides is consistent with a globally dampened immune remodeling signature in paucibacillary disease. TBC peptides map to established immunodominant M. tuberculosis proteins: Ag85A (major secreted antigen, vaccine candidate), PPE60 (PE/PPE immune evasion family, a serodiagnostic candidate from exovesicles for latent tuberculosis), EchA3 (crotonase, a known serodiagnosis biomarker), GarA (immunodominant antigen, a known biomarker for differentiating latent and acute tuberculosis), Culp1 (cutinase, a known serodiagnosis marker for acute tuberculosis), LprG (conserved lipoprotein, good known serodiagnosis biomarker), Mpt63 (known serodiagnostic antigen), Mpt70 (serodiagnostic antigen). RRL peptides are enriched for hydrophobic residues (49% vs 38% in TBC) and alpha-helix-forming amino acids, suggesting that they sample antibody reactivity against structural host protein motifs.*

**Peptide 1** (amino acids 260-271) from Rv3804c (Ag85A, Mpt44, mycolyl transferase, fibronectin-binding protein A, FbpA), a well-known antigen and pathogenicity factor. The peptide is a part of the known IEDB database antibody epitope 49333 (position 259-303). It was one of the top 13 antibody-binding antigens associated with tuberculosis found by Kunnath-Velayudhan et al. (35) and Kunnath-Velayudhan et al. (36), had CD4-T cell-responsive epitopes and induced antibody responses (Nayak et al., (37)). It was selected for serodiagnosis in the Uganda-TB-panel (Shete *et al.* (31)), it was one of the three best HIV-TB biomarkers (Jaganath *et al.* (29)), and it was a discriminatory biomarker for HIV-specific evaluated long-term tuberculosis infection (LTBI) / acute tuberculosis (ATB) IgM-binding (Nziza *et al.* 2022(30)).

**Peptide 3** (amino acid 93-104) from Rv3478 (PE family protein PPE60, immune evasion family) was found to be highly expressed in extracellular vesicles, sheded by *Mycobacterium tuberculosis*, and was suggested to be included in serodiagnostic panels (Schirmer *et al.* (27)).

**Peptide 5** (amino acids 80-91) from Rv0632c (EchA3), enoyl-CoA hydratase EchA3 or crotonase, was identified by three previous studies as a candidate biomarker. It was one of the top 13 antibody-binding antigens associated with tuberculosis found by Kunnath-Velayudhan *et al.* 35), had CD4-T cell-responsive epitopes and induced antibody responses (Nayak *et al.*, 37); and was selected for tuberculosis serodiagnosis as one of the best 14 proteins by Zhou *et al.* 2015 (28).

**Peptide 6** (amino acids 92-103) is derived from Rv1827 (GarA, Cfp17, glycogen accumulation regulator). Instead classified as a transcription factor it was nevertheless identified as a secreted protein (Cornejo-Granados et al. (32)) and was included in a discriminatory panel for differentiating latent and active tuberculosis by serodiagnosis (Li *et al*. (33)). In this study, the best panel combination, comprised of Rv0934, Rv3881c, Rv1860 and Rv1827, achieved 67.3% sensitivity at 91.2% specificity for discriminating ATB from LTBI, and 71.2% sensitivity at 96.3% specificity for discriminating ATB from healthy controls. This was the first published assay to discriminate latent and acute tuberculosis infections by a serodiagnosis panel.

**Peptide 7** (amino acid 101-112) and peptide 14 (amino acid 64-75) originate from Rv1984c (Culp1, Cfp21, Clp1), cutinase, a secreted protein, which hydrolyzes cutin. The protein was also identified by Kunnath-Velayudhan, S. et al. (35) as one of the top 13 antigens. It was additionally found to be a useful biomarker for serodiagnosis of active tuberculosis by incorporation in a 10-antigen-panel (Khaliq *et al.* (34)).

**Peptide 9** (amino acid 157-168) from Rv1411c (LprG, P27), a conserved core mycobacterial lipoprotein, was identified previously by several studies as a candidate biomarker for tuberculosis. It was one of the top 13 antibody-binding antigens associated with tuberculosis found by Kunnath-Velayudhan *et al.* (35) and Kunnath-Velayudhan *et al.* (36), had CD4-T cell-responsive epitopes and induced antibody responses (Nayak *et al.*, (37)); and was selected for tuberculosis serodiagnosis as one of the best 14 proteins by Zhou *et al.* (28), there it was included in the best-performing panel of 6 antigens.

**Peptide 10** (amino acid 96-107) from Rv1926c (antigen Mpt63/MPB63, 16 kDa immunoprotective extracellular protein), is partially overlapping a well-known mapped IEDB database epitope (No. 103283, position 85-99). It was also identified as a potent CD4 T-cell antigen (Nayak *et al*. (37)). It was additionally found to be a useful biomarker for serodiagnosis of active tuberculosis by incorporation in a 10-antigen-panel (Khaliq *et al.* (34)).

**Supplementary Table 2. Elastic Net feature reduction across peptide libraries.**

Number of peptides retained (non-zero Elastic Net coefficients) per outer CV fold. EN parameters: SGDClassifier with log loss, l1_ratio = 0.5, α = 0.001, balanced class weights, 5,000 iterations.

| Library | Start | Fold 1 | Fold 2 | Fold 3 | Fold 4 | Fold 5 | Mean | % retained |
|---|---|---|---|---|---|---|---|---|
| ALL | 6,936 | 2,301 | 2,325 | 2,298 | 2,340 | 2,326 | 2,318 | 33.4% |
| TBC | 4,317 | 1,830 | 1,856 | 1,838 | 1,862 | 1,834 | 1,844 | 42.7% |
| RRL | 2,496 | 978 | 992 | 981 | 998 | 976 | 985 | 39.5% |

*The Elastic Net consistently removes approximately two-thirds of peptides from the ALL library and approximately 40–60% from the subset libraries. The retained fraction is higher for TBC (42.7%) and RRL (39.5%) than for ALL (33.4%), reflecting the greater homogeneity of peptides within each library. Variation across folds is minimal (±2%), indicating stable feature reduction.*

**Supplementary Table 3. Peptide number sweep: AUC by panel size, library, and cohort.**

Mean AUC (± SD) from nested 5-fold CV at each panel size N. Best AUC per column highlighted.

| N | ALL Full | ALL HIV− | TBC Full | TBC HIV− | RRL Full | RRL HIV− |
|---|---|---|---|---|---|---|
| 1 | 0.660 | 0.656 | 0.596 | 0.651 | 0.563 | 0.681 |
| 2 | 0.618 | 0.699 | 0.580 | 0.663 | 0.538 | 0.707 |
| 3 | 0.717 | 0.703 | 0.554 | 0.648 | 0.511 | 0.768 |
| 5 | 0.614 | 0.727 | 0.634 | 0.793 | 0.574 | 0.739 |
| 10 | 0.636 | 0.700 | 0.562 | 0.696 | 0.500 | 0.668 |
| 20 | 0.570 | 0.739 | 0.560 | 0.676 | 0.489 | 0.655 |
| 50 | 0.561 | 0.740 | 0.551 | 0.696 | 0.482 | 0.611 |
| 100 | 0.512 | 0.673 | 0.479 | 0.510 | 0.571 | 0.633 |

*In the full cohort, optimal performance is consistently achieved with small panels (N = 1–5), reflecting the high-dimensional noise inherent in peptide array data. In the HIV-negative subcohort, performance is more stable across panel sizes, with optimal N varying by library (ALL: N = 50; TBC: N = 5; RRL: N = 3). The inverted relationship between N and AUC in the full cohort (adding features degrades performance) provides empirical evidence that the Elastic Net pre-filter, while removing ~66% of peptides, does not fully eliminate noise features.*

**Supplementary Table 4. *M. tuberculosis* proteins identified by the EN→RF peptide selection pipeline.**

Proteins represented among the top 15 selected peptides for S−C+ detection (HIV-negative). All proteins are established immunodominant *M. tuberculosis* antigens with known roles in TB immunity and/or diagnostics.

| Rv number | Protein | Function | Diagnostic relevance | Peptides selected |
|---|---|---|---|---|
| Rv3804c | Ag85A (FbpB) | Major secreted antigen; mycolyl transferase; cell wall biosynthesis | Vaccine candidate (MVA85A); serodiagnostic antigen | 1 (rank 1) |
| Rv3478 | PPE60 | PE/PPE family protein; immune evasion; T-cell epitope | Serodiagnostic candidate, highly expressed in extracellular vesicles | 2 (rank 3) |
| Rv0632c | EchA3 | Secreted crotonase | Serodiagnostic candidate | 1 (rank 5) |
| Rv1827 | GarA | Glycogen accumulation regulator; immunodominant antigen | T-cell and B-cell antigen, differentiates latent/acute TB in serodiagnosis | 1 (rank 6) |
| Rv1984c | Culp1 | cutinase | Serodiagnostic antigen | 2 (ranks 7, 14) |
| Rv1411c | LprG | Conserved lipoprotein | Serodiagnostic antigen | 1 (rank 9) |
| Rv1926c | Mpt63 | Secreted, immunoprotective | Serodiagnostic antigen | 1 (rank 10) |

*The independent rediscovery of these established immunodominant TB antigens through an unsupervised data-driven pipeline (Elastic Net pre-filtering followed by Random Forest importance ranking) provides external validation of the feature selection methodology. Notably, Ag85A, Culp1 and Mpt63 are widely used TB serodiagnostic antigens, LprG and GarA were also included in serodiagnostic panels, where GarA has potential to differentiate latent and acute tuberculosis. EchA3 and PPE60 are candidate serodiagnostic biomarkers, where PPE60 is also a candidate for differentiating latent/active tuberculosis. The convergence of our machine learning pipeline on these known targets, without any prior biological information guiding the selection, demonstrates that the pipeline identifies biologically meaningful features.*

### Supplementary Note

**Supplementary Note 1. Elastic Net pre-filtering: rationale and implementation details.**

The peptide-level classifier operates in a high-dimensional setting ($p = 6{,}936$ features, $n = 105$ samples), where direct application of Random Forest feature selection is unstable due to the low feature-to-sample ratio. Without pre-filtering, each tree in the Random Forest considers only a random subset of $\sqrt{p} \approx 83$ features at each split, and with 6,936 features, the probability that any specific informative peptide is evaluated at any given split is low. This leads to high variance in feature importance estimates across bootstrap samples and CV folds.

To address this, we introduced an Elastic Net pre-filter (SGDClassifier with log loss, l1_ratio = 0.5, $\alpha = 0.001$) as the first stage of feature selection within each training fold. The Elastic Net combines L1 (lasso) and L2 (ridge) penalties: the L1 component drives uninformative feature coefficients to exactly zero, while the L2 component stabilizes the solution when features are correlated. The mixed penalty (l1_ratio = 0.5) provides a balance between aggressive sparsity and grouped selection of correlated features.

The regularization strength ($\alpha = 0.001$) was chosen to be permissive rather than aggressive, retaining approximately one-third of all features (33.4% for the ALL library). This conservative threshold ensures that potentially informative peptides are not discarded, at the cost of retaining some noise features that the subsequent Random Forest ranking can eliminate. The $\alpha$ parameter was not tuned via inner cross-validation; instead, it was fixed across all analyses to minimize researcher degrees of freedom and reduce the risk of overfitting. The consistent retention fractions across libraries (33–43%) and across CV folds (SD < 2%) support the stability of this choice.

After Elastic Net pre-filtering, the feature-to-tree ratio improves from $6{,}936/500 = 0.07$ to approximately $2{,}300/500 = 0.22$, and the number of features considered per split increases from $\sqrt{6{,}936} \approx 83$ to $\sqrt{2{,}300} \approx 48$ out of a smaller, more informative pool. This yields more stable and reproducible feature importance rankings, as confirmed by the increase in the number of core peptides (selected in all 5 CV folds) from 1 (without EN) to 2 (with EN) for the ALL library.
