## Supplementary material for "High-density peptide arrays detect tuberculosis through immune remodeling, not only antigen recognition alone": code: TB_Pipeline_All_ElasticNet.ipynb

```
# %% [markdown]
# TB Immune Remodeling — Full Master Pipeline (Steps 4–10)
## With Elastic Net Pre-Filter for Feature Reduction

**Continues from:** `TB_Immune_Remodeling_Pipeline.ipynb` (Steps 0–3)

| Step | Content |
|------|---------|
| 4 | Peptide-level classifier (**Elastic Net → RF → LR**) |
| 5 | Meta-model (stacked remodeling + peptide) |
| 6 | HIV-stratified analysis |
| 7 | Effect size analysis (Cliff's delta per Z-feature) |
| 8 | Volcano plot (visual only, no leakage) |
| 9 | Final outputs with bootstrap CI + DeLong test |
| 10 | Publication figures |

**Feature reduction:** 6,936 peptides → Elastic Net (~2,300 survivors) → RF ranking → top N → LR

**Design principles:** Strictly leakage-free. All fitting on discovery fold only.
Nested 5-fold CV. Focus: peptide selection for improved diagnosis.

# %% [markdown]
## 0 — Imports & Data Loading

*(Same as Steps 0–3 notebook — run this first)*

# %% [code]
import numpy as np
import pandas as pd
import matplotlib.pyplot as plt
import matplotlib.gridspec as gridspec
import seaborn as sns
import warnings
warnings.filterwarnings('ignore')

from scipy.stats import mannwhitneyu, norm
from scipy.stats import false_discovery_control   # Python ≥3.11 / scipy ≥1.11
from statsmodels.stats.multitest import multipletests

from sklearn.ensemble import RandomForestClassifier
from sklearn.linear_model import LogisticRegression
from sklearn.linear_model import SGDClassifier  # Elastic Net pre-filter
from sklearn.preprocessing import StandardScaler
from sklearn.decomposition import PCA
from sklearn.model_selection import StratifiedKFold, GridSearchCV, cross_val_predict
from sklearn.metrics import roc_auc_score, roc_curve

RANDOM_STATE = 42
np.random.seed(RANDOM_STATE)

plt.rcParams.update({'figure.dpi': 150, 'font.size': 11,
                     'axes.titlesize': 12, 'axes.labelsize': 11})
sns.set_style('whitegrid')

print('Libraries loaded ✓')

# %% [code]
# ── Load peptide intensity matrix ──────────────────────────────────────────
DATA_PATH = r"D:/Users/ha6057/Desktop/TBC_AI/Files for Simulation/all_signals_TBC.csv"      # peptide signals
META_PATH = r"D:/Users/ha6057/Desktop/TBC_AI/Files for Simulation/df_samples_all.xls"
PEP_CAT_PATH = r"D:/Users/ha6057/Desktop/TBC_AI/Files for Simulation/df_peptide_origin_with_Category.xlsx"

df_raw = pd.read_csv(DATA_PATH, index_col=0)
sample_ids   = df_raw.index.tolist()
all_peptides = df_raw.columns.tolist()
X_all        = df_raw.values.astype(np.float64)
n_samples, n_peptides = X_all.shape
print(f'Peptide matrix: {n_samples} samples × {n_peptides} peptides')

# ── Load metadata ──────────────────────────────────────────────────────────
meta = pd.read_csv(META_PATH).set_index('ID').loc[sample_ids]

def map_tb(v):
    return 0 if v == 'Negative' else 1

y_tb       = np.array([map_tb(v) for v in meta['TB Status']], dtype=int)
hiv_status = np.array([1 if v == 'HIV+' else 0
                        for v in meta['HIV_status']], dtype=int)
country    = meta['Country'].apply(
    lambda x: 'Viet Nam' if 'iet' in str(x) else x).values
age        = meta['Age'].values
gender     = meta['Gender'].values

print(f'TB+: {y_tb.sum()}  TB-: {(y_tb==0).sum()}')
print(f'HIV+: {hiv_status.sum()}  HIV-: {(hiv_status==0).sum()}')
print(f'Countries: {pd.Series(country).value_counts().to_dict()}')

# ── Peptide subsets ────────────────────────────────────────────────────────
# TBC = all peptides in this file (replace idx_tbc if you have a sub-list)
idx_tbc = np.arange(n_peptides)   # ← update if you have a specific TBC list
X_tbc   = X_all[:, idx_tbc]
tbc_peptide_names = [all_peptides[i] for i in idx_tbc]

print(f'TBC peptides for analysis: {len(idx_tbc)}')

# %% [markdown]
## Shared Helper Functions (Steps 0–3 reused here)

# %% [code]
def shannon_entropy(X):
    s = X.sum(axis=1, keepdims=True)
    s = np.where(s == 0, 1, s)
    p = np.clip(X / s, 1e-12, None)
    return -np.sum(p * np.log(p), axis=1)

def gini_coefficient(X):
    g = []
    for row in X:
        x  = np.sort(row)
        cx = np.cumsum(x)
        g.append((len(x)+1 - 2*cx.sum()/cx[-1])/len(x) if cx[-1] > 0 else 0)
    return np.array(g)

def build_Z(X_tr, X_te, y_tr, reg_lambda=1e-3):
    """Build immune state vector Z. Returns Z_tr, Z_te (scaled), Z_tr_raw, Z_te_raw."""
    # Distributional metrics
    ent_tr = shannon_entropy(X_tr);  gin_tr = gini_coefficient(X_tr)
    ent_te = shannon_entropy(X_te);  gin_te = gini_coefficient(X_te)
    # PCA
    sc = StandardScaler()
    Xtr_sc = sc.fit_transform(X_tr);  Xte_sc = sc.transform(X_te)
    pca = PCA(n_components=2, random_state=RANDOM_STATE)
    pcs_tr = pca.fit_transform(Xtr_sc);  pcs_te = pca.transform(Xte_sc)
    # Mahalanobis from TB- training centroid
    ref  = pcs_tr[y_tr == 0]
    mu   = ref.mean(axis=0)
    cov  = np.cov(ref, rowvar=False) + reg_lambda * np.eye(2)
    ci   = np.linalg.inv(cov)
    mah  = lambda P: np.array([np.sqrt((x-mu)@ci@(x-mu)) for x in P])
    Z_tr_raw = np.column_stack([ent_tr, gin_tr, pcs_tr[:,0], pcs_tr[:,1], mah(pcs_tr)])
    Z_te_raw = np.column_stack([ent_te, gin_te, pcs_te[:,0], pcs_te[:,1], mah(pcs_te)])
    szZ = StandardScaler()
    return szZ.fit_transform(Z_tr_raw), szZ.transform(Z_te_raw), Z_tr_raw, Z_te_raw

def tune_and_fit_lr(X_tr, y_tr, inner_cv, C_GRID):
    """Inner-CV tuning + final LR fit. Returns fitted model."""
    gs = GridSearchCV(
        LogisticRegression(max_iter=2000, random_state=RANDOM_STATE, solver='lbfgs'),
        C_GRID, cv=inner_cv, scoring='roc_auc', n_jobs=-1)
    gs.fit(X_tr, y_tr)
    lr = LogisticRegression(C=gs.best_params_['C'], max_iter=2000,
                             random_state=RANDOM_STATE, solver='lbfgs')
    lr.fit(X_tr, y_tr)
    return lr, gs.best_params_['C']

def cliffs_delta(a, b):
    """Cliff's delta effect size between arrays a and b."""
    n1, n2 = len(a), len(b)
    more = sum(1 for x in a for y in b if x > y)
    less = sum(1 for x in a for y in b if x < y)
    return (more - less) / (n1 * n2)

def bootstrap_auc_ci(y_true, y_prob, n_boot=1000, ci=0.95, seed=42):
    """Bootstrap confidence interval for AUC."""
    rng  = np.random.default_rng(seed)
    aucs = []
    n    = len(y_true)
    for _ in range(n_boot):
        idx  = rng.integers(0, n, size=n)
        if len(np.unique(y_true[idx])) < 2:
            continue
        aucs.append(roc_auc_score(y_true[idx], y_prob[idx]))
    lo = np.percentile(aucs, (1-ci)/2*100)
    hi = np.percentile(aucs, (1-(1-ci)/2)*100)
    return lo, hi

def delong_test(y_true, prob_a, prob_b):
    """
    Simplified DeLong-style paired AUC comparison using
    the Hanley-McNeil correlation method.
    Returns z-statistic and p-value (two-tailed).
    """
    def compute_Vxy(p_pos, p_neg):
        return np.array([np.mean(px > p_neg) for px in p_pos])
    def compute_Vyx(p_pos, p_neg):
        return np.array([np.mean(pn < p_pos) for pn in p_neg])

    y = np.asarray(y_true)
    pos_mask = y == 1;  neg_mask = y == 0
    n1, n0   = pos_mask.sum(), neg_mask.sum()

    def auc_vars(probs):
        pp, pn = probs[pos_mask], probs[neg_mask]
        V10    = compute_Vxy(pp, pn)
        V01    = compute_Vyx(pp, pn)
        auc    = V10.mean()
        s10    = np.var(V10, ddof=1) / n1
        s01    = np.var(V01, ddof=1) / n0
        return auc, s10, s01, V10, V01

    auc_a, s10a, s01a, V10a, V01a = auc_vars(prob_a)
    auc_b, s10b, s01b, V10b, V01b = auc_vars(prob_b)

    r_10 = np.corrcoef(V10a, V10b)[0,1] if n1 > 2 else 0
    r_01 = np.corrcoef(V01a, V01b)[0,1] if n0 > 2 else 0

    var_diff = s10a + s01a + s10b + s01b - 2*r_10*np.sqrt(s10a*s10b) - 2*r_01*np.sqrt(s01a*s01b)
    se       = np.sqrt(max(var_diff, 1e-12))
    z        = (auc_a - auc_b) / se
    p        = 2 * (1 - norm.cdf(abs(z)))
    return float(z), float(p)

C_GRID   = {'C': [0.001, 0.01, 0.1, 1, 10, 100]}
outer_cv = StratifiedKFold(n_splits=5, shuffle=True, random_state=RANDOM_STATE)
inner_cv = StratifiedKFold(n_splits=5, shuffle=True, random_state=RANDOM_STATE)
TOP_N_PEPTIDES = 50

print('Helper functions defined ✓')

# %% [markdown]
## STEP 3 — Model 1: Remodeling Classifier (Recap)
*(Re-run here so both models share the same CV splits)*

# %% [code]
remod = dict(y_true=[], y_prob=[], fold_aucs=[], Z_train=[], Z_test=[],
             train_indices=[], test_indices=[])

for fold_i, (train_idx, test_idx) in enumerate(outer_cv.split(X_tbc, y_tb)):
    X_tr, X_te = X_tbc[train_idx], X_tbc[test_idx]
    y_tr, y_te = y_tb[train_idx],   y_tb[test_idx]

    Z_tr, Z_te, Z_tr_raw, _ = build_Z(X_tr, X_te, y_tr)
    lr, best_C = tune_and_fit_lr(Z_tr, y_tr, inner_cv, C_GRID)
    y_prob = lr.predict_proba(Z_te)[:, 1]

    remod['y_true'].extend(y_te.tolist())
    remod['y_prob'].extend(y_prob.tolist())
    remod['fold_aucs'].append(roc_auc_score(y_te, y_prob))
    remod['Z_train'].append(Z_tr_raw)
    remod['Z_test'].append(Z_te)
    remod['train_indices'].append(train_idx)
    remod['test_indices'].append(test_idx)
    print(f'  Remodeling Fold {fold_i+1}: AUC={remod["fold_aucs"][-1]:.3f}  C={best_C}')

print(f'\nRemodeling AUC = {np.mean(remod["fold_aucs"]):.3f} ± {np.std(remod["fold_aucs"]):.3f}')

# %% [markdown]
## STEP 4 — Model 2: Peptide-Level Classifier (Elastic Net → RF → LR)

### Two-stage feature reduction pipeline per fold:
1. **Standardize** intensities (training scaler only)
2. **Elastic Net pre-filter** (SGDClassifier, l1_ratio=0.5, α=0.001) → removes ~67% noise peptides
3. **Random Forest** feature importance on EN survivors → rank peptides
4. **Select top 50** peptides by RF importance
5. **Inner CV** to tune LR `C`
6. **Predict** on test set

### Why two-stage selection?
- With p=6,936 >> n=105, direct RF ranking is unstable (feature/tree ratio = 0.07)
- **Elastic Net** (L1+L2) efficiently sets uninformative coefficients to zero
- After EN: ~2,300 survivors → RF feature/tree ratio improves to ~0.22 (3× better)
- Both EN and RF fitted on training fold only → no leakage

# %% [code]
peptide = dict(y_true=[], y_prob=[], fold_aucs=[], fold_top_peptides=[],
               fold_rf_importance=[], fold_best_C=[], fold_en_survivors=[])

# RF hyperparameters — fixed (no inner tuning for RF to save compute)
RF_PARAMS = dict(n_estimators=500, max_depth=None, min_samples_leaf=3,
                 class_weight='balanced', random_state=RANDOM_STATE, n_jobs=-1)

for fold_i, (train_idx, test_idx) in enumerate(outer_cv.split(X_tbc, y_tb)):
    X_tr_raw, X_te_raw = X_tbc[train_idx], X_tbc[test_idx]
    y_tr, y_te         = y_tb[train_idx],   y_tb[test_idx]

    # ── 4.1 Standardize ──────────────────────────────────────────────────
    scaler = StandardScaler()
    X_tr   = scaler.fit_transform(X_tr_raw)
    X_te   = scaler.transform(X_te_raw)

    # ── 4.2 Elastic Net pre-filter ───────────────────────────────────────
    en = SGDClassifier(loss='log_loss', penalty='elasticnet',
                       l1_ratio=0.5, alpha=0.001, max_iter=5000,
                       random_state=RANDOM_STATE, class_weight='balanced')
    en.fit(X_tr, y_tr)
    en_mask     = np.abs(en.coef_.ravel()) > 0
    n_survivors = en_mask.sum()
    peptide['fold_en_survivors'].append(n_survivors)

    X_tr_en = X_tr[:, en_mask]
    X_te_en = X_te[:, en_mask]

    # ── 4.3 Random Forest feature selection (on EN survivors) ─────────
    rf = RandomForestClassifier(**RF_PARAMS)
    rf.fit(X_tr_en, y_tr)

    importance    = rf.feature_importances_
    ranked_idx    = np.argsort(importance)[::-1]
    top50_idx     = ranked_idx[:TOP_N_PEPTIDES]

    # Map back to original peptide names
    survivor_indices = np.where(en_mask)[0]
    top50_names = [tbc_peptide_names[survivor_indices[i]] for i in top50_idx]
    top50_imp   = importance[top50_idx]

    peptide['fold_top_peptides'].append(top50_names)
    peptide['fold_rf_importance'].append(
        {tbc_peptide_names[survivor_indices[i]]: importance[i]
         for i in range(len(survivor_indices))})

    # ── 4.4 Reduce to top 50 ─────────────────────────────────────────────
    X_tr_sel = X_tr_en[:, top50_idx]
    X_te_sel = X_te_en[:, top50_idx]

    # ── 4.5 Inner CV + 4.6 Train LR ──────────────────────────────────────
    lr, best_C = tune_and_fit_lr(X_tr_sel, y_tr, inner_cv, C_GRID)
    y_prob     = lr.predict_proba(X_te_sel)[:, 1]
    fold_auc   = roc_auc_score(y_te, y_prob)

    peptide['y_true'].extend(y_te.tolist())
    peptide['y_prob'].extend(y_prob.tolist())
    peptide['fold_aucs'].append(fold_auc)
    peptide['fold_best_C'].append(best_C)

    print(f'  Peptide  Fold {fold_i+1}: EN surv={n_survivors}/{X_tr.shape[1]} '
          f'({100*n_survivors/X_tr.shape[1]:.1f}%)  AUC={fold_auc:.3f}  '
          f'C={best_C}  top3={top50_names[:3]}')

print(f'\nPeptide AUC = {np.mean(peptide["fold_aucs"]):.3f} ± {np.std(peptide["fold_aucs"]):.3f}')
print(f'EN survivors per fold: {peptide["fold_en_survivors"]}')
print(f'Mean reduction: {n_peptides} → {np.mean(peptide["fold_en_survivors"]):.0f} '
      f'({100*np.mean(peptide["fold_en_survivors"])/n_peptides:.1f}%)')

# %% [markdown]
### 4B — Peptide Selection Consensus
Which peptides are selected **consistently** across all 5 folds?  
Consistent selection = higher confidence that the peptide is genuinely informative.

# %% [code]
from collections import Counter

# Count how many folds each peptide was selected in
all_selected = [p for fold in peptide['fold_top_peptides'] for p in fold]
selection_freq = Counter(all_selected)

df_consensus = pd.DataFrame([
    {'Peptide': pep, 'Folds_Selected': cnt,
     'Mean_RF_Importance': np.mean([
         peptide['fold_rf_importance'][f].get(pep, 0)
         for f in range(5)])}
    for pep, cnt in selection_freq.most_common()
]).sort_values(['Folds_Selected', 'Mean_RF_Importance'], ascending=[False, False])

# Consensus tiers
core_peptides    = df_consensus[df_consensus['Folds_Selected'] == 5]['Peptide'].tolist()
robust_peptides  = df_consensus[df_consensus['Folds_Selected'] >= 4]['Peptide'].tolist()
stable_peptides  = df_consensus[df_consensus['Folds_Selected'] >= 3]['Peptide'].tolist()

print(f'Total unique peptides ever selected: {len(df_consensus)}')
print(f'Core     (5/5 folds): {len(core_peptides)}')
print(f'Robust   (≥4/5 folds): {len(robust_peptides)}')
print(f'Stable   (≥3/5 folds): {len(stable_peptides)}')
print(f'\nTop 20 most consistently selected peptides:')
print(df_consensus.head(20).to_string(index=False, float_format='%.5f'))

# Save consensus table
df_consensus.to_csv('peptide_selection_consensus.csv', index=False)
print('\nSaved: peptide_selection_consensus.csv')

# %% [markdown]
### 4C — Peptide Number Sweep
How many top peptides give the best AUC?  
Test N = 1, 2, 3, ... 200 — **using same EN → RF folds, no leakage.**

# %% [code]
SWEEP_NS = [1, 2, 3, 4, 5, 10, 15, 20, 30, 50, 75, 100, 150, 200]
sweep_results = {n: [] for n in SWEEP_NS}

print('Running peptide number sweep (EN → RF → LR)...')
for fold_i, (train_idx, test_idx) in enumerate(outer_cv.split(X_tbc, y_tb)):
    X_tr_raw, X_te_raw = X_tbc[train_idx], X_tbc[test_idx]
    y_tr, y_te         = y_tb[train_idx],   y_tb[test_idx]

    scaler = StandardScaler()
    X_tr   = scaler.fit_transform(X_tr_raw)
    X_te   = scaler.transform(X_te_raw)

    # Elastic Net pre-filter
    en = SGDClassifier(loss='log_loss', penalty='elasticnet',
                       l1_ratio=0.5, alpha=0.001, max_iter=5000,
                       random_state=RANDOM_STATE, class_weight='balanced')
    en.fit(X_tr, y_tr)
    en_mask = np.abs(en.coef_.ravel()) > 0
    X_tr_en = X_tr[:, en_mask]
    X_te_en = X_te[:, en_mask]

    # Fit RF once per fold on EN survivors
    rf = RandomForestClassifier(**RF_PARAMS)
    rf.fit(X_tr_en, y_tr)
    ranked = np.argsort(rf.feature_importances_)[::-1]

    for N in SWEEP_NS:
        top_idx  = ranked[:min(N, len(ranked))]
        gs = GridSearchCV(
            LogisticRegression(max_iter=2000, random_state=RANDOM_STATE, solver='lbfgs'),
            C_GRID, cv=inner_cv, scoring='roc_auc', n_jobs=-1)
        gs.fit(X_tr_en[:, top_idx], y_tr)
        lr = LogisticRegression(C=gs.best_params_['C'], max_iter=2000,
                                 random_state=RANDOM_STATE, solver='lbfgs')
        lr.fit(X_tr_en[:, top_idx], y_tr)
        auc = roc_auc_score(y_te, lr.predict_proba(X_te_en[:, top_idx])[:,1])
        sweep_results[N].append(auc)

    print(f'  Fold {fold_i+1} done')

sweep_means = [np.mean(sweep_results[n]) for n in SWEEP_NS]
sweep_stds  = [np.std(sweep_results[n])  for n in SWEEP_NS]
best_N      = SWEEP_NS[np.argmax(sweep_means)]
best_auc    = max(sweep_means)

print(f'\nOptimal N = {best_N} peptides (AUC = {best_auc:.3f})')
for n, m, s in zip(SWEEP_NS, sweep_means, sweep_stds):
    marker = ' ← BEST' if n == best_N else ''
    print(f'  N={n:4d}: {m:.3f} ± {s:.3f}{marker}')

# %% [markdown]
## STEP 5 — Meta-Model (Stacked Classifier)

**Logic:** If remodeling and peptide models capture different information,  
stacking them should improve AUC.  
**Keep meta-model only if ΔAUC > 0.02** over the better individual model.

# %% [code]
meta_model = dict(y_true=[], y_prob=[], fold_aucs=[])

# Rebuild per-fold predictions for stacking (must be out-of-fold)
fold_splits = list(outer_cv.split(X_tbc, y_tb))

for fold_i, (train_idx, test_idx) in enumerate(fold_splits):
    y_tr, y_te = y_tb[train_idx], y_tb[test_idx]

    # ── Generate OOF predictions on TRAINING set via inner CV ────────────
    # This ensures stacking inputs are themselves out-of-fold

    # Remodeling OOF (training)
    Z_tr, Z_te, _, _ = build_Z(X_tbc[train_idx], X_tbc[test_idx], y_tr)
    oof_remod_tr = cross_val_predict(
        LogisticRegression(C=1, max_iter=2000, random_state=RANDOM_STATE, solver='lbfgs'),
        Z_tr, y_tr, cv=inner_cv, method='predict_proba')[:, 1]

    # Peptide OOF (training) — with Elastic Net pre-filter
    scaler = StandardScaler()
    X_tr_sc = scaler.fit_transform(X_tbc[train_idx])
    X_te_sc = scaler.transform(X_tbc[test_idx])
    en = SGDClassifier(loss='log_loss', penalty='elasticnet',
                       l1_ratio=0.5, alpha=0.001, max_iter=5000,
                       random_state=RANDOM_STATE, class_weight='balanced')
    en.fit(X_tr_sc, y_tr)
    en_mask = np.abs(en.coef_.ravel()) > 0
    X_tr_sc_en = X_tr_sc[:, en_mask]
    X_te_sc_en = X_te_sc[:, en_mask]
    rf = RandomForestClassifier(**RF_PARAMS)
    rf.fit(X_tr_sc_en, y_tr)
    top_idx = np.argsort(rf.feature_importances_)[::-1][:TOP_N_PEPTIDES]
    oof_pep_tr = cross_val_predict(
        LogisticRegression(C=1, max_iter=2000, random_state=RANDOM_STATE, solver='lbfgs'),
        X_tr_sc_en[:, top_idx], y_tr, cv=inner_cv, method='predict_proba')[:, 1]

    # ── Stack on training OOF predictions ────────────────────────────────
    S_tr = np.column_stack([oof_remod_tr, oof_pep_tr])
    meta_lr, best_C = tune_and_fit_lr(S_tr, y_tr, inner_cv,
                                       {'C': [0.01, 0.1, 1, 10]})

    # ── Test predictions for both base models ────────────────────────────
    lr_remod, _ = tune_and_fit_lr(Z_tr, y_tr, inner_cv, C_GRID)
    p_remod_te  = lr_remod.predict_proba(Z_te)[:, 1]

    lr_pep, _   = tune_and_fit_lr(X_tr_sc_en[:, top_idx], y_tr, inner_cv, C_GRID)
    p_pep_te    = lr_pep.predict_proba(X_te_sc_en[:, top_idx])[:, 1]

    S_te    = np.column_stack([p_remod_te, p_pep_te])
    p_meta  = meta_lr.predict_proba(S_te)[:, 1]
    fold_auc = roc_auc_score(y_te, p_meta)

    meta_model['y_true'].extend(y_te.tolist())
    meta_model['y_prob'].extend(p_meta.tolist())
    meta_model['fold_aucs'].append(fold_auc)
    print(f'  Meta Fold {fold_i+1}: AUC={fold_auc:.3f}')

# ── Decision: keep meta-model? ────────────────────────────────────────────
auc_remod = np.mean(remod['fold_aucs'])
auc_pep   = np.mean(peptide['fold_aucs'])
auc_meta  = np.mean(meta_model['fold_aucs'])
best_base = max(auc_remod, auc_pep)
delta_meta = auc_meta - best_base

KEEP_META = delta_meta >= 0.02
print(f'\nRemodeling AUC : {auc_remod:.3f}')
print(f'Peptide    AUC : {auc_pep:.3f}')
print(f'Meta       AUC : {auc_meta:.3f}  (Δ = {delta_meta:+.3f})')
print(f'Keep meta-model: {"YES ✓" if KEEP_META else "NO — improvement < 0.02"}')


# %% [markdown]
## STEP 6 — HIV-Stratified Analysis

**Questions:**
1. Does the classifier work equally well in HIV− patients only?
2. Does HIV status *modify* the TB classification signal?
3. Does adding HIV as a covariate improve classification?

# %% [code]
# ── 6A: HIV- only pipeline ────────────────────────────────────────────────
hiv_neg_mask = (hiv_status == 0)
X_hiv_neg    = X_tbc[hiv_neg_mask]
y_hiv_neg    = y_tb[hiv_neg_mask]

print(f'HIV- cohort: {hiv_neg_mask.sum()} samples  '
      f'(TB+={y_hiv_neg.sum()}, TB-={(y_hiv_neg==0).sum()})')

hiv_neg_remod  = dict(fold_aucs=[], y_true=[], y_prob=[])
hiv_neg_peptide = dict(fold_aucs=[], y_true=[], y_prob=[])

cv_hiv = StratifiedKFold(n_splits=5, shuffle=True, random_state=RANDOM_STATE)

for fold_i, (tr_idx, te_idx) in enumerate(cv_hiv.split(X_hiv_neg, y_hiv_neg)):
    Xtr, Xte = X_hiv_neg[tr_idx], X_hiv_neg[te_idx]
    ytr, yte = y_hiv_neg[tr_idx], y_hiv_neg[te_idx]

    # Remodeling
    Ztr, Zte, _, _ = build_Z(Xtr, Xte, ytr)
    lr, _ = tune_and_fit_lr(Ztr, ytr, inner_cv, C_GRID)
    p = lr.predict_proba(Zte)[:, 1]
    hiv_neg_remod['fold_aucs'].append(roc_auc_score(yte, p))
    hiv_neg_remod['y_true'].extend(yte.tolist())
    hiv_neg_remod['y_prob'].extend(p.tolist())

    # Peptide — with Elastic Net pre-filter
    sc = StandardScaler()
    Xtr_sc = sc.fit_transform(Xtr); Xte_sc = sc.transform(Xte)
    en = SGDClassifier(loss='log_loss', penalty='elasticnet',
                       l1_ratio=0.5, alpha=0.001, max_iter=5000,
                       random_state=RANDOM_STATE, class_weight='balanced')
    en.fit(Xtr_sc, ytr)
    en_mask = np.abs(en.coef_.ravel()) > 0
    Xtr_en = Xtr_sc[:, en_mask]; Xte_en = Xte_sc[:, en_mask]
    rf = RandomForestClassifier(**RF_PARAMS)
    rf.fit(Xtr_en, ytr)
    top_idx = np.argsort(rf.feature_importances_)[::-1][:TOP_N_PEPTIDES]
    lr2, _ = tune_and_fit_lr(Xtr_en[:, top_idx], ytr, inner_cv, C_GRID)
    p2 = lr2.predict_proba(Xte_en[:, top_idx])[:, 1]
    hiv_neg_peptide['fold_aucs'].append(roc_auc_score(yte, p2))
    hiv_neg_peptide['y_true'].extend(yte.tolist())
    hiv_neg_peptide['y_prob'].extend(p2.tolist())

# ── 6B: HIV as covariate ──────────────────────────────────────────────────
covar_aucs = []
for fold_i, (train_idx, test_idx) in enumerate(outer_cv.split(X_tbc, y_tb)):
    y_tr, y_te = y_tb[train_idx], y_tb[test_idx]
    Z_tr, Z_te, _, _ = build_Z(X_tbc[train_idx], X_tbc[test_idx], y_tr)
    hiv_tr = hiv_status[train_idx].reshape(-1,1)
    hiv_te = hiv_status[test_idx].reshape(-1,1)
    Z_tr_h = np.hstack([Z_tr, hiv_tr])
    Z_te_h = np.hstack([Z_te, hiv_te])
    lr, _ = tune_and_fit_lr(Z_tr_h, y_tr, inner_cv, C_GRID)
    covar_aucs.append(roc_auc_score(y_te, lr.predict_proba(Z_te_h)[:,1]))

print('\n── HIV Analysis Results ──────────────────────────────')
print(f'Full cohort   | Remodeling: {np.mean(remod["fold_aucs"]):.3f}  '
      f'Peptide: {np.mean(peptide["fold_aucs"]):.3f}')
print(f'HIV- only     | Remodeling: {np.mean(hiv_neg_remod["fold_aucs"]):.3f}  '
      f'Peptide: {np.mean(hiv_neg_peptide["fold_aucs"]):.3f}')
print(f'HIV covariate | Remodeling+HIV: {np.mean(covar_aucs):.3f}')
print(f'\nΔ AUC (HIV- vs Full) | Remodeling: '
      f'{np.mean(hiv_neg_remod["fold_aucs"]) - np.mean(remod["fold_aucs"]):+.3f}')
print(f'Δ AUC (HIV- vs Full) | Peptide   : '
      f'{np.mean(hiv_neg_peptide["fold_aucs"]) - np.mean(peptide["fold_aucs"]):+.3f}')


# %% [markdown]
## STEP 7 — Effect Size Analysis (Cliff's Delta per Z-Feature)

**Within training folds only** — no test data used.  
Cliff's delta: −1 = TB− always higher, +1 = TB+ always higher, 0 = no difference.  
|δ| interpretation: <0.15 negligible, 0.15–0.33 small, 0.33–0.47 medium, >0.47 large.

# %% [code]
Z_LABELS = ['Entropy', 'Gini', 'PC1', 'PC2', 'Mahalanobis']

fold_cliffs  = {z: [] for z in Z_LABELS}
fold_mwu_p   = {z: [] for z in Z_LABELS}
fold_mwu_stat = {z: [] for z in Z_LABELS}

for fold_i, (train_idx, _) in enumerate(outer_cv.split(X_tbc, y_tb)):
    y_tr   = y_tb[train_idx]
    _, _, Z_tr_raw, _ = build_Z(X_tbc[train_idx],
                                  X_tbc[train_idx[:5]],  # dummy test
                                  y_tr)

    for fi, fname in enumerate(Z_LABELS):
        vals_neg = Z_tr_raw[y_tr == 0, fi]
        vals_pos = Z_tr_raw[y_tr == 1, fi]

        # Mann-Whitney U
        stat, p = mannwhitneyu(vals_pos, vals_neg, alternative='two-sided')
        fold_mwu_stat[fname].append(stat)
        fold_mwu_p[fname].append(p)

        # Cliff's delta (TB+ vs TB-)
        cd = cliffs_delta(vals_pos.tolist(), vals_neg.tolist())
        fold_cliffs[fname].append(cd)

# Summary table
def cliff_label(d):
    a = abs(d)
    if a < 0.15: return 'negligible'
    elif a < 0.33: return 'small'
    elif a < 0.47: return 'medium'
    else: return 'large'

rows = []
for fname in Z_LABELS:
    cd_mean = np.mean(fold_cliffs[fname])
    cd_std  = np.std(fold_cliffs[fname])
    p_mean  = np.mean(fold_mwu_p[fname])
    rows.append({'Feature': fname,
                 "Cliff's δ (mean)": cd_mean,
                 "Cliff's δ (std)": cd_std,
                 '|δ|': abs(cd_mean),
                 'Magnitude': cliff_label(cd_mean),
                 'MWU p (mean)': p_mean})

df_effect = pd.DataFrame(rows).sort_values('|δ|', ascending=False)
print("Effect Sizes (Cliff's Delta, TB+ vs TB−, training folds only):")
print(df_effect.to_string(index=False, float_format='%.4f'))

df_effect.to_csv('results_effect_sizes.csv', index=False)
print('\nSaved: results_effect_sizes.csv')

# %% [markdown]
## STEP 8 — Volcano Plot (Visual Only — No Classifier Use)

**Per-peptide comparison TB+ vs TB−.**  
Computed within training folds only, then averaged.  
FDR correction (Benjamini-Hochberg).  
⚠️ Volcano is **visualization only** — not used for classification.

# %% [code]
# Compute volcano stats within training folds (leakage-free)
fold_log2fc = []
fold_pvals  = []

for train_idx, _ in outer_cv.split(X_tbc, y_tb):
    X_tr = X_tbc[train_idx]
    y_tr = y_tb[train_idx]
    pos_tr = X_tr[y_tr == 1]
    neg_tr = X_tr[y_tr == 0]

    # log2 fold change: mean(TB+) / mean(TB-)
    mean_pos = pos_tr.mean(axis=0) + 1e-9
    mean_neg = neg_tr.mean(axis=0) + 1e-9
    log2fc   = np.log2(mean_pos / mean_neg)
    fold_log2fc.append(log2fc)

    # Mann-Whitney per peptide
    pvals = []
    for j in range(X_tr.shape[1]):
        _, p = mannwhitneyu(pos_tr[:, j], neg_tr[:, j], alternative='two-sided')
        pvals.append(p)
    fold_pvals.append(pvals)

# Average across folds
mean_log2fc = np.mean(fold_log2fc, axis=0)
mean_pvals  = np.mean(fold_pvals, axis=0)

# FDR correction (Benjamini-Hochberg)
_, fdr_pvals, _, _ = multipletests(mean_pvals, alpha=0.05, method='fdr_bh')
neg_log10_fdr = -np.log10(fdr_pvals + 1e-300)

# Build volcano dataframe
df_volcano = pd.DataFrame({
    'Peptide': tbc_peptide_names,
    'log2FC': mean_log2fc,
    'FDR_p': fdr_pvals,
    'neg_log10_FDR': neg_log10_fdr,
})
df_volcano['Significant'] = (fdr_pvals < 0.05) & (np.abs(mean_log2fc) > 0.5)
df_volcano['Direction']   = np.where(mean_log2fc > 0, 'Up in TB+', 'Up in TB-')

n_sig = df_volcano['Significant'].sum()
n_up  = ((df_volcano['Significant']) & (mean_log2fc > 0)).sum()
n_dn  = ((df_volcano['Significant']) & (mean_log2fc < 0)).sum()

print(f'Significant peptides (FDR<0.05, |log2FC|>0.5): {n_sig}')
print(f'  Up in TB+: {n_up}  |  Up in TB-: {n_dn}')
print(f'\nTop 15 upregulated in TB+:')
print(df_volcano[df_volcano['log2FC']>0].sort_values('neg_log10_FDR',
    ascending=False).head(15)[['Peptide','log2FC','FDR_p']].to_string(
    index=False, float_format='%.4f'))

df_volcano.to_csv('peptide_volcano_data.csv', index=False)
print('\nSaved: peptide_volcano_data.csv')

# %% [markdown]
## STEP 9 — Final Output: Bootstrap CI + DeLong Test

# %% [code]
y_true_arr = np.array(remod['y_true'])
p_remod    = np.array(remod['y_prob'])
p_peptide  = np.array(peptide['y_prob'])
p_meta_arr = np.array(meta_model['y_prob'])

# Bootstrap CI
ci_remod   = bootstrap_auc_ci(y_true_arr, p_remod)
ci_peptide = bootstrap_auc_ci(y_true_arr, p_peptide)
ci_meta    = bootstrap_auc_ci(y_true_arr, p_meta_arr)

# DeLong paired test: remodeling vs peptide
z_rp, p_rp = delong_test(y_true_arr, p_remod, p_peptide)
# DeLong: meta vs best base
best_base_prob = p_remod if auc_remod >= auc_pep else p_peptide
z_mb, p_mb     = delong_test(y_true_arr, p_meta_arr, best_base_prob)

# HIV-stratified bootstrap
y_hivneg_arr    = np.array(hiv_neg_remod['y_true'])
p_hivneg_remod  = np.array(hiv_neg_remod['y_prob'])
p_hivneg_pep    = np.array(hiv_neg_peptide['y_prob'])
ci_hivneg_remod = bootstrap_auc_ci(y_hivneg_arr, p_hivneg_remod)
ci_hivneg_pep   = bootstrap_auc_ci(y_hivneg_arr, p_hivneg_pep)

print('='*65)
print('FINAL RESULTS SUMMARY')
print('='*65)
print(f"\n{'Model':<30} {'Mean AUC':>10} {'95% CI':>20} {'Fold SD':>10}")
print('-'*65)

rows_out = [
    ('Remodeling (full)',       np.mean(remod['fold_aucs']),         ci_remod,   np.std(remod['fold_aucs'])),
    ('Peptide LR (full)',       np.mean(peptide['fold_aucs']),       ci_peptide, np.std(peptide['fold_aucs'])),
    ('Meta-model',              np.mean(meta_model['fold_aucs']),    ci_meta,    np.std(meta_model['fold_aucs'])),
    ('Remodeling (HIV- only)',  np.mean(hiv_neg_remod['fold_aucs']), ci_hivneg_remod, np.std(hiv_neg_remod['fold_aucs'])),
    ('Peptide LR (HIV- only)',  np.mean(hiv_neg_peptide['fold_aucs']),ci_hivneg_pep,  np.std(hiv_neg_peptide['fold_aucs'])),
    ('Remodeling + HIV covar.', np.mean(covar_aucs), ('—','—'), np.std(covar_aucs)),
]

df_final = []
for name, auc, ci, sd in rows_out:
    ci_str = f'{ci[0]:.3f}–{ci[1]:.3f}' if ci[0] != '—' else '—'
    print(f'{name:<30} {auc:>10.3f} {ci_str:>20} {sd:>10.3f}')
    df_final.append({'Model': name, 'AUC': auc, 'CI_lo': ci[0] if ci[0]!='—' else np.nan,
                     'CI_hi': ci[1] if ci[1]!='—' else np.nan, 'SD': sd})

print(f'\nDeLong test: Remodeling vs Peptide  z={z_rp:.3f}  p={p_rp:.4f}')
print(f'DeLong test: Meta vs Best-base      z={z_mb:.3f}  p={p_mb:.4f}')
print(f'Meta-model decision: {"KEEP (ΔAUC ≥ 0.02)" if KEEP_META else "DISCARD (ΔAUC < 0.02)"}')

pd.DataFrame(df_final).to_csv('results_final_summary.csv', index=False)
print('\nSaved: results_final_summary.csv')

# %% [markdown]
## STEP 10 — Publication Figures

| Figure | Content |
|--------|----------|
| Fig 1  | Study design + CV diagram |
| Fig 2  | Remodeling feature separation (Z-space) |
| Fig 3  | AUC comparison: remodeling vs peptide |
| Fig 4  | HIV suppression effect |
| Fig 5  | Volcano + peptide consensus |

# %% [code]
# ═══════════════════════════════════════════════════════════════════════════
# FIGURE 1 — Study Design & CV Diagram
# ═══════════════════════════════════════════════════════════════════════════
fig, axes = plt.subplots(1, 2, figsize=(16, 6))

# Left: cohort overview
ax = axes[0]
ax.axis('off')
countries = pd.Series(country).value_counts()
x_pos = {'South Africa': 0.2, 'Peru': 0.5, 'Viet Nam': 0.8}
colors_c = {'South Africa': '#9C27B0', 'Peru': '#FF9800', 'Viet Nam': '#009688'}

# Draw cohort boxes
for ctry, n_c in countries.items():
    xp = x_pos.get(ctry, 0.5)
    rect = plt.Rectangle((xp-0.12, 0.55), 0.24, 0.25,
                          facecolor=colors_c.get(ctry,'grey'), alpha=0.3,
                          edgecolor='black', transform=ax.transAxes)
    ax.add_patch(rect)
    ax.text(xp, 0.71, ctry, ha='center', va='center',
            transform=ax.transAxes, fontweight='bold', fontsize=10)
    tb_pos = ((country == ctry) & (y_tb == 1)).sum()
    tb_neg = ((country == ctry) & (y_tb == 0)).sum()
    ax.text(xp, 0.62, f'n={n_c}\nTB+={tb_pos} TB-={tb_neg}',
            ha='center', va='center', transform=ax.transAxes, fontsize=9)

ax.text(0.5, 0.92, f'Multi-Country TB Cohort  (n={n_samples})',
        ha='center', va='center', transform=ax.transAxes,
        fontsize=13, fontweight='bold')
ax.text(0.5, 0.46, f'Peptide library: {n_peptides} TBC peptides',
        ha='center', va='center', transform=ax.transAxes, fontsize=10, style='italic')

# Pipeline summary boxes
steps = [
    ('Model 1: Remodeling', 'Entropy + Gini + PC1\n+ PC2 + Mahalanobis\n→ Logistic Regression', '#2196F3'),
    ('Model 2: Peptide LR', f'RF: Top {TOP_N_PEPTIDES} peptides\n(TBC library)\n→ Logistic Regression', '#4CAF50'),
    ('Meta-model', 'Stack probabilities\nfrom Model 1+2\n→ Logistic Regression', '#FF9800'),
]
for si, (title, desc, color) in enumerate(steps):
    xp = 0.18 + si * 0.32
    rect = plt.Rectangle((xp-0.13, 0.05), 0.26, 0.28,
                          facecolor=color, alpha=0.15,
                          edgecolor=color, linewidth=2, transform=ax.transAxes)
    ax.add_patch(rect)
    ax.text(xp, 0.28, title, ha='center', va='center',
            transform=ax.transAxes, fontweight='bold', fontsize=9, color=color)
    ax.text(xp, 0.15, desc, ha='center', va='center',
            transform=ax.transAxes, fontsize=8)
ax.set_title('A  |  Study Design', fontweight='bold', pad=10)

# Right: CV diagram
ax2 = axes[1]
ax2.axis('off')
ax2.set_title('B  |  Nested Cross-Validation Scheme', fontweight='bold', pad=10)

fold_colors = ['#BBDEFB','#C8E6C9','#FFE0B2','#F8BBD0','#E1BEE7']
for fi in range(5):
    for bi in range(5):
        color = '#F44336' if bi == fi else fold_colors[fi]
        label = 'Test' if bi == fi else 'Train'
        rect = plt.Rectangle((bi*0.17+0.09, 0.72-fi*0.13), 0.15, 0.1,
                              facecolor=color, alpha=0.7, edgecolor='white',
                              linewidth=2, transform=ax2.transAxes)
        ax2.add_patch(rect)
        ax2.text(bi*0.17+0.165, 0.775-fi*0.13, label,
                 ha='center', va='center', transform=ax2.transAxes,
                 fontsize=7, color='white' if bi==fi else 'black')

for fi in range(5):
    ax2.text(0.07, 0.775-fi*0.13, f'Fold {fi+1}',
             ha='right', va='center', transform=ax2.transAxes, fontsize=9)

ax2.text(0.5, 0.25, 'Within each Train set: inner 5-fold CV\nfor LR hyperparameter C tuning',
         ha='center', transform=ax2.transAxes, fontsize=10,
         bbox=dict(boxstyle='round', facecolor='wheat', alpha=0.5))
ax2.text(0.5, 0.10, '⚠  All transformations (PCA, scaler, RF selection)\nfitted on Train only — Test never seen during fitting',
         ha='center', transform=ax2.transAxes, fontsize=9, color='#B71C1C')

plt.tight_layout()
plt.savefig('Fig1_study_design.png', bbox_inches='tight', dpi=180)
plt.show()
print('Fig 1 saved.')

# %% [code]
# ═══════════════════════════════════════════════════════════════════════════
# FIGURE 2 — Remodeling Feature Separation
# ═══════════════════════════════════════════════════════════════════════════

# Collect all test-fold Z_raw values
all_Z_raw, all_y_vis, all_hiv_vis, all_country_vis = [], [], [], []
for fold_i, (train_idx, test_idx) in enumerate(outer_cv.split(X_tbc, y_tb)):
    X_tr, X_te = X_tbc[train_idx], X_tbc[test_idx]
    y_tr, y_te = y_tb[train_idx],   y_tb[test_idx]
    _, _, Z_tr_raw, Z_te_raw = build_Z(X_tr, X_te, y_tr)
    all_Z_raw.extend(Z_te_raw.tolist())
    all_y_vis.extend(y_te.tolist())
    all_hiv_vis.extend(hiv_status[test_idx].tolist())
    all_country_vis.extend(country[test_idx].tolist())

Z_mat     = np.array(all_Z_raw)
y_vis     = np.array(all_y_vis)
hiv_vis   = np.array(all_hiv_vis)
ctry_vis  = np.array(all_country_vis)

fig, axes = plt.subplots(2, 3, figsize=(16, 10))
TB_COLOR  = {0: '#2196F3', 1: '#F44336'}
TB_NAME   = {0: 'TB−', 1: 'TB+'}

for ai, (fi, fname) in enumerate(enumerate(Z_LABELS)):
    ax = axes[ai // 3, ai % 3]
    for label in [0, 1]:
        vals = Z_mat[y_vis == label, fi]
        ax.hist(vals, bins=20, alpha=0.55, color=TB_COLOR[label],
                label=TB_NAME[label], density=True)
    stat, p = mannwhitneyu(Z_mat[y_vis==0,fi], Z_mat[y_vis==1,fi],
                            alternative='two-sided')
    cd = cliffs_delta(Z_mat[y_vis==1,fi].tolist(), Z_mat[y_vis==0,fi].tolist())
    sig = '***' if p<0.001 else '**' if p<0.01 else '*' if p<0.05 else 'ns'
    ax.set_title(f'{fname}\nMWU p={p:.3f} {sig}  δ={cd:+.3f} ({cliff_label(cd)})',
                 fontsize=10)
    ax.set_xlabel(fname); ax.set_ylabel('Density')
    ax.legend(fontsize=9)

# 6th panel: PC1 vs PC2 scatter
ax6 = axes[1, 2]
for label in [0, 1]:
    m = y_vis == label
    ax6.scatter(Z_mat[m, 2], Z_mat[m, 3],
                c=TB_COLOR[label], label=TB_NAME[label],
                alpha=0.65, s=40, edgecolors='white', linewidth=0.4)
ax6.set_xlabel('PC1'); ax6.set_ylabel('PC2')
ax6.set_title('PC1 vs PC2\n(immune manifold)')
ax6.legend()

fig.suptitle('Figure 2 — Immune Remodeling Feature Separation (TB+ vs TB−)',
             fontsize=13, fontweight='bold')
plt.tight_layout()
plt.savefig('Fig2_remodeling_separation.png', bbox_inches='tight', dpi=180)
plt.show()
print('Fig 2 saved.')

# %% [code]
# ═══════════════════════════════════════════════════════════════════════════
# FIGURE 3 — AUC Comparison: Remodeling vs Peptide
# ═══════════════════════════════════════════════════════════════════════════
fig, axes = plt.subplots(1, 3, figsize=(17, 6))

COLORS_MODEL = {'Remodeling': '#2196F3', 'Peptide LR': '#4CAF50', 'Meta-model': '#FF9800'}

# Left: Fold AUC swarm/box
ax = axes[0]
models_data = [
    ('Remodeling',  remod['fold_aucs']),
    ('Peptide LR',  peptide['fold_aucs']),
    ('Meta-model',  meta_model['fold_aucs']),
]
for xi, (name, aucs) in enumerate(models_data):
    color = COLORS_MODEL[name]
    bp = ax.boxplot(aucs, positions=[xi], patch_artist=True, widths=0.4,
                    medianprops={'color':'black','linewidth':2})
    bp['boxes'][0].set_facecolor(color); bp['boxes'][0].set_alpha(0.6)
    jitter = np.random.normal(0, 0.04, len(aucs))
    ax.scatter(np.full(5, xi)+jitter, aucs, color=color, s=50, zorder=5,
               edgecolors='white')
ax.axhline(0.5, color='red', ls='--', alpha=0.4, lw=1.5)
ax.set_xticks([0,1,2]); ax.set_xticklabels([m[0] for m in models_data])
ax.set_ylabel('ROC-AUC'); ax.set_ylim(0.2, 1.0)
ax.set_title('A  |  Fold AUC Distribution', fontweight='bold')

# Middle: ROC curves
ax2 = axes[1]
for name, y_t, y_p, color in [
    ('Remodeling', remod['y_true'],      remod['y_prob'],       '#2196F3'),
    ('Peptide LR', peptide['y_true'],    peptide['y_prob'],     '#4CAF50'),
    ('Meta-model', meta_model['y_true'], meta_model['y_prob'],  '#FF9800'),
]:
    yt = np.array(y_t); yp = np.array(y_p)
    fpr, tpr, _ = roc_curve(yt, yp)
    auc_val = roc_auc_score(yt, yp)
    ci_lo, ci_hi = bootstrap_auc_ci(yt, yp, n_boot=500)
    ax2.plot(fpr, tpr, color=color, lw=2.5,
             label=f'{name}\nAUC={auc_val:.3f} [{ci_lo:.3f}–{ci_hi:.3f}]')
ax2.plot([0,1],[0,1],'k--',alpha=0.3, lw=1.5)
ax2.set_xlabel('False Positive Rate'); ax2.set_ylabel('True Positive Rate')
ax2.set_title('B  |  ROC Curves (pooled, bootstrap CI)',fontweight='bold')
ax2.legend(loc='lower right', fontsize=8)

# Right: Peptide sweep curve
ax3 = axes[2]
ax3.errorbar(SWEEP_NS, sweep_means, yerr=sweep_stds, fmt='o-',
             color='#4CAF50', linewidth=2, markersize=7, capsize=5,
             label='Peptide LR AUC')
ax3.axhline(np.mean(remod['fold_aucs']), color='#2196F3', ls='--',
            lw=2, label=f'Remodeling AUC = {np.mean(remod["fold_aucs"]):.3f}')
ax3.axvline(best_N, color='red', ls=':', lw=1.5, label=f'Optimal N={best_N}')
ax3.scatter([best_N], [best_auc], color='red', s=120, zorder=6)
ax3.set_xlabel('Number of Top Peptides Selected')
ax3.set_ylabel('ROC-AUC')
ax3.set_title(f'C  |  Peptide Selection Sweep\nOptimal N = {best_N} (AUC={best_auc:.3f})',
              fontweight='bold')
ax3.legend(fontsize=9)
ax3.set_ylim(0.35, 1.0)

fig.suptitle('Figure 3 — Model Performance Comparison', fontsize=13, fontweight='bold')
plt.tight_layout()
plt.savefig('Fig3_auc_comparison.png', bbox_inches='tight', dpi=180)
plt.show()
print('Fig 3 saved.')

# %% [code]
# ═══════════════════════════════════════════════════════════════════════════
# FIGURE 4 — HIV Suppression / Modification Effect
# ═══════════════════════════════════════════════════════════════════════════
fig, axes = plt.subplots(1, 3, figsize=(16, 6))

# Left: AUC full vs HIV- bars
ax = axes[0]
bar_data = [
    ('Remodeling\n(Full)', np.mean(remod['fold_aucs']),         np.std(remod['fold_aucs']),         '#2196F3'),
    ('Remodeling\n(HIV−)', np.mean(hiv_neg_remod['fold_aucs']), np.std(hiv_neg_remod['fold_aucs']), '#90CAF9'),
    ('Peptide LR\n(Full)', np.mean(peptide['fold_aucs']),       np.std(peptide['fold_aucs']),       '#4CAF50'),
    ('Peptide LR\n(HIV−)', np.mean(hiv_neg_peptide['fold_aucs']),np.std(hiv_neg_peptide['fold_aucs']),'#A5D6A7'),
    ('Remod\n+HIV covar',  np.mean(covar_aucs),                 np.std(covar_aucs),                 '#FF9800'),
]
for xi, (name, m, s, color) in enumerate(bar_data):
    ax.bar(xi, m, yerr=s, color=color, edgecolor='black', alpha=0.8, capsize=6)
    ax.text(xi, m+s+0.01, f'{m:.3f}', ha='center', va='bottom', fontsize=9, fontweight='bold')
ax.axhline(0.5, color='red', ls='--', alpha=0.4)
ax.set_xticks(range(len(bar_data)))
ax.set_xticklabels([b[0] for b in bar_data], fontsize=8)
ax.set_ylabel('Mean AUC ± SD'); ax.set_ylim(0.3, 1.0)
ax.set_title('A  |  Full Cohort vs HIV− Only', fontweight='bold')

# Middle: ROC curves HIV+ vs HIV- subsets (remodeling model)
ax2 = axes[1]
fpr_f, tpr_f, _ = roc_curve(remod['y_true'], remod['y_prob'])
auc_f = roc_auc_score(remod['y_true'], remod['y_prob'])
ax2.plot(fpr_f, tpr_f, color='#2196F3', lw=2.5,
         label=f'Full cohort (AUC={auc_f:.3f})')
fpr_h, tpr_h, _ = roc_curve(hiv_neg_remod['y_true'], hiv_neg_remod['y_prob'])
auc_h = roc_auc_score(hiv_neg_remod['y_true'], hiv_neg_remod['y_prob'])
ax2.plot(fpr_h, tpr_h, color='#90CAF9', lw=2.5, ls='--',
         label=f'HIV− only (AUC={auc_h:.3f})')
ax2.plot([0,1],[0,1],'k--',alpha=0.3)
ax2.set_xlabel('FPR'); ax2.set_ylabel('TPR')
ax2.set_title('B  |  ROC: Remodeling Model\n(Full vs HIV−)', fontweight='bold')
ax2.legend(fontsize=9)

# Right: Z-feature distributions split by HIV
ax3 = axes[2]
# Show Mahalanobis by TB x HIV group
mah_vals = Z_mat[:, 4]
groups = [
    (mah_vals[(y_vis==0)&(hiv_vis==0)], 'TB−/HIV−', '#2196F3'),
    (mah_vals[(y_vis==0)&(hiv_vis==1)], 'TB−/HIV+', '#90CAF9'),
    (mah_vals[(y_vis==1)&(hiv_vis==0)], 'TB+/HIV−', '#F44336'),
    (mah_vals[(y_vis==1)&(hiv_vis==1)], 'TB+/HIV+', '#EF9A9A'),
]
for i, (vals, label, color) in enumerate(groups):
    if len(vals) > 0:
        vp = ax3.violinplot(vals, positions=[i], showmedians=True, widths=0.65)
        for pc in vp['bodies']: pc.set_facecolor(color); pc.set_alpha(0.7)
        vp['cmedians'].set_color('black'); vp['cmedians'].set_linewidth(2)
        jitter = np.random.normal(0, 0.07, len(vals))
        ax3.scatter(np.full(len(vals),i)+jitter, vals, color=color,
                    s=25, alpha=0.6, zorder=5, edgecolors='white', lw=0.3)
ax3.set_xticks(range(4))
ax3.set_xticklabels([g[1] for g in groups], fontsize=9)
ax3.set_ylabel('Mahalanobis Distance')
ax3.set_title('C  |  Immune Displacement\nby TB × HIV group', fontweight='bold')

fig.suptitle('Figure 4 — HIV Effect on TB Immune Classification', fontsize=13, fontweight='bold')
plt.tight_layout()
plt.savefig('Fig4_HIV_effect.png', bbox_inches='tight', dpi=180)
plt.show()
print('Fig 4 saved.')

# %% [code]
# ═══════════════════════════════════════════════════════════════════════════
# FIGURE 5 — Volcano + Peptide Consensus
# ═══════════════════════════════════════════════════════════════════════════
fig, axes = plt.subplots(1, 2, figsize=(17, 7))

# Left: Volcano
ax = axes[0]

# Colors by significance and direction
def volcano_color(row):
    if not row['Significant']: return '#B0BEC5'
    return '#F44336' if row['log2FC'] > 0 else '#2196F3'

df_volcano['color'] = df_volcano.apply(volcano_color, axis=1)
df_volcano['selected_consensus'] = df_volcano['Peptide'].isin(core_peptides)

# Plot non-significant
mask_ns = ~df_volcano['Significant']
ax.scatter(df_volcano.loc[mask_ns,'log2FC'],
           df_volcano.loc[mask_ns,'neg_log10_FDR'],
           c='#B0BEC5', s=12, alpha=0.4, label='Not significant')

# Plot significant
for direction, color, label in [('Up in TB+','#F44336','Up in TB+'),
                                  ('Up in TB-','#2196F3','Up in TB−')]:
    mask = df_volcano['Significant'] & (df_volcano['Direction'] == direction)
    ax.scatter(df_volcano.loc[mask,'log2FC'],
               df_volcano.loc[mask,'neg_log10_FDR'],
               c=color, s=30, alpha=0.75, label=label)

# Highlight consensus peptides
mask_core = df_volcano['Significant'] & df_volcano['selected_consensus']
if mask_core.any():
    ax.scatter(df_volcano.loc[mask_core,'log2FC'],
               df_volcano.loc[mask_core,'neg_log10_FDR'],
               c='gold', s=80, edgecolors='black', linewidths=0.8,
               zorder=6, label=f'Core consensus (5/5 folds, n={len(core_peptides)})')

# Threshold lines
fdr_thresh = -np.log10(0.05)
ax.axhline(fdr_thresh, color='gray', ls='--', lw=1, alpha=0.7,
           label=f'FDR=0.05 (−log10={fdr_thresh:.1f})')
ax.axvline(0.5,  color='gray', ls=':', lw=1, alpha=0.7)
ax.axvline(-0.5, color='gray', ls=':', lw=1, alpha=0.7)

# Label top 10 peptides
top_label = df_volcano[df_volcano['Significant']].nlargest(10,'neg_log10_FDR')
for _, row in top_label.iterrows():
    ax.annotate(row['Peptide'], (row['log2FC'], row['neg_log10_FDR']),
                xytext=(5, 3), textcoords='offset points', fontsize=6.5,
                color='black', arrowprops=dict(arrowstyle='-', lw=0.5, color='gray'))

ax.set_xlabel('log₂ Fold Change (TB+ / TB−)')
ax.set_ylabel('−log₁₀(FDR p-value)')
ax.set_title(f'A  |  Volcano: {n_sig} significant peptides\n'
             f'(FDR<0.05, |log₂FC|>0.5) — VISUALIZATION ONLY',
             fontweight='bold')
ax.legend(fontsize=8, loc='upper left')

# Right: Peptide consensus bar chart
ax2 = axes[1]
df_top_con = df_consensus.head(30).copy()
bar_colors_con = []
for _, row in df_top_con.iterrows():
    n_folds = row['Folds_Selected']
    if n_folds == 5:   bar_colors_con.append('#F44336')
    elif n_folds == 4: bar_colors_con.append('#FF9800')
    elif n_folds == 3: bar_colors_con.append('#FFC107')
    else:              bar_colors_con.append('#90CAF9')

bars = ax2.barh(range(len(df_top_con)),
                df_top_con['Mean_RF_Importance'],
                color=bar_colors_con, edgecolor='black', alpha=0.85)
ax2.set_yticks(range(len(df_top_con)))
ax2.set_yticklabels(df_top_con['Peptide'], fontsize=8)
ax2.invert_yaxis()
ax2.set_xlabel('Mean RF Importance (across folds)')
ax2.set_title(f'B  |  Top 30 Peptides by RF Consensus\n'
              f'(color = folds selected: red=5/5, orange=4/5, yellow=3/5)',
              fontweight='bold')

# Annotate folds
for i, (_, row) in enumerate(df_top_con.iterrows()):
    ax2.text(row['Mean_RF_Importance'] * 1.01, i,
             f"{int(row['Folds_Selected'])}/5",
             va='center', fontsize=7.5)

# Legend
from matplotlib.patches import Patch
legend_els = [Patch(facecolor='#F44336', label='5/5 folds (core)'),
              Patch(facecolor='#FF9800', label='4/5 folds (robust)'),
              Patch(facecolor='#FFC107', label='3/5 folds (stable)'),
              Patch(facecolor='#90CAF9', label='≤2/5 folds')]
ax2.legend(handles=legend_els, fontsize=8, loc='lower right')

fig.suptitle('Figure 5 — Peptide Signatures: Volcano & Selection Consensus',
             fontsize=13, fontweight='bold')
plt.tight_layout()
plt.savefig('Fig5_volcano_consensus.png', bbox_inches='tight', dpi=180)
plt.show()
print('Fig 5 saved.')

# %% [markdown]
## Final Comprehensive Summary + Export

# %% [code]
print('='*70)
print('MASTER PIPELINE — FINAL SUMMARY')
print('='*70)

print(f"""
COHORT
  Samples  : {n_samples}  (TB+={y_tb.sum()}, TB-={(y_tb==0).sum()})
  HIV+     : {hiv_status.sum()}  |  HIV-: {(hiv_status==0).sum()}
  Countries: South Africa / Peru / Viet Nam
  Peptides : {n_peptides} TBC

MODEL PERFORMANCE (nested 5-fold CV, mean AUC ± SD)
  Model 1 — Remodeling     : {np.mean(remod['fold_aucs']):.3f} ± {np.std(remod['fold_aucs']):.3f}
  Model 2 — Peptide LR     : {np.mean(peptide['fold_aucs']):.3f} ± {np.std(peptide['fold_aucs']):.3f}
  Meta-model               : {np.mean(meta_model['fold_aucs']):.3f} ± {np.std(meta_model['fold_aucs']):.3f}
  Meta-model kept          : {'YES' if KEEP_META else 'NO (ΔAUC < 0.02)'}

PEPTIDE SELECTION (EN pre-filter → top {TOP_N_PEPTIDES} per fold via RF)\n"
"  EN survivors (mean)  : {np.mean(peptide['fold_en_survivors']):.0f} / {n_peptides} ({100*np.mean(peptide['fold_en_survivors'])/n_peptides:.1f}%)
  Core peptides (5/5 folds): {len(core_peptides)}
  Robust      (≥4/5 folds) : {len(robust_peptides)}
  Stable      (≥3/5 folds) : {len(stable_peptides)}
  Optimal N (sweep)        : {best_N} peptides (AUC={best_auc:.3f})

HIV ANALYSIS
  Remodeling HIV- only     : {np.mean(hiv_neg_remod['fold_aucs']):.3f} ± {np.std(hiv_neg_remod['fold_aucs']):.3f}
  Peptide LR HIV- only     : {np.mean(hiv_neg_peptide['fold_aucs']):.3f} ± {np.std(hiv_neg_peptide['fold_aucs']):.3f}
  Remodeling + HIV covar.  : {np.mean(covar_aucs):.3f} ± {np.std(covar_aucs):.3f}

STATISTICAL COMPARISONS
  DeLong: Remodeling vs Peptide  z={z_rp:.3f}  p={p_rp:.4f}
  DeLong: Meta vs Best-base      z={z_mb:.3f}  p={p_mb:.4f}
""")

print('EFFECT SIZES (Cliff\'s delta, training folds):')
print(df_effect[['Feature',"Cliff's δ (mean)",'Magnitude','MWU p (mean)']].to_string(
    index=False, float_format='%.4f'))

print(f"""
VOLCANO SUMMARY
  Significant peptides     : {n_sig}  (FDR<0.05, |log2FC|>0.5)
  Up in TB+                : {n_up}
  Up in TB-                : {n_dn}

FILES SAVED
  peptide_selection_consensus.csv
  peptide_volcano_data.csv
  results_effect_sizes.csv
  results_final_summary.csv
  Fig1_study_design.png
  Fig2_remodeling_separation.png
  Fig3_auc_comparison.png
  Fig4_HIV_effect.png
  Fig5_volcano_consensus.png
""")

print('='*70)
print('✓ Pipeline complete.')


# %% [code]


# %% [code]


# %% [code]


# %% [code]
```
