## Supplementary material for "High-density peptide arrays detect tuberculosis through immune remodeling, not only antigen recognition alone": code: TB_Pipeline_RRL_ElasticNet.ipynb

```
# %% [markdown]
# TB Immune Remodeling — RRL Library Analysis
## Applying Steps 4–10 Pipeline to Random Reference Library (RRL2) Only

**Purpose:** Replicate the full TB_Pipeline_Steps4to10 workflow using **only the 2,496 RRL peptides**
instead of all 6,936 peptides, and evaluate how AUC changes when restricting to HIV-negative samples.

| Step | Content |
|------|---------|
| 0 | Data loading & RRL subset extraction |
| 3 | Remodeling classifier (recap on RRL) |
| 4 | Peptide-level classifier (RF → top-N → LR) |
| 4B | Peptide selection consensus |
| 4C | Peptide number sweep |
| 5 | Meta-model (stacked) |
| 6 | HIV-stratified analysis ← **KEY COMPARISON** |
| 7 | Effect size analysis |
| 8 | Volcano plot |
| 9 | Bootstrap CI + DeLong |
| 10 | Publication figures |

# %% [markdown]
## 0 — Imports & Configuration

# %% [code]
import numpy as np
import pandas as pd
import matplotlib.pyplot as plt
import matplotlib.gridspec as gridspec
import matplotlib.patches as mpatches
import seaborn as sns
import warnings
warnings.filterwarnings('ignore')

from scipy.stats import mannwhitneyu, norm
from statsmodels.stats.multitest import multipletests
from collections import Counter

from sklearn.ensemble import RandomForestClassifier
from sklearn.linear_model import LogisticRegression
from sklearn.linear_model import SGDClassifier  # Elastic Net pre-filter
from sklearn.preprocessing import StandardScaler
from sklearn.decomposition import PCA
from sklearn.model_selection import StratifiedKFold, GridSearchCV, cross_val_predict
from sklearn.metrics import roc_auc_score, roc_curve

RANDOM_STATE = 42
np.random.seed(RANDOM_STATE)

plt.rcParams.update({'figure.dpi': 150, 'font.size': 11,
                     'axes.titlesize': 12, 'axes.labelsize': 11})
sns.set_style('whitegrid')

print('Libraries loaded')

# %% [markdown]
## 0B — Load Data & Extract RRL Subset

Load the full signal matrix + metadata, then **filter columns to RRL peptides only**
using the peptide category file.

# %% [code]
# ── Paths (adjust to your environment) ──────────────────────────────────────
DATA_PATH = r"D:/Users/ha6057/Desktop/TBC_AI/Files for Simulation/all_signals_TBC.csv"
META_PATH = r"D:/Users/ha6057/Desktop/TBC_AI/Files for Simulation/df_samples_all.xls"
PEP_CAT_PATH = r"D:/Users/ha6057/Desktop/TBC_AI/Files for Simulation/df_peptide_origin_with_Category.xlsx"

# ── Load peptide matrix ────────────────────────────────────────────────────
df_raw = pd.read_csv(DATA_PATH, index_col=0)
sample_ids   = df_raw.index.tolist()
all_peptides = df_raw.columns.tolist()
n_samples_total = df_raw.shape[0]
print(f'Full matrix: {df_raw.shape[0]} samples x {df_raw.shape[1]} peptides')

# ── Load metadata ──────────────────────────────────────────────────────────
meta = pd.read_csv(META_PATH).set_index('ID').loc[sample_ids]

def map_tb(v):
    return 0 if v == 'Negative' else 1

y_tb       = np.array([map_tb(v) for v in meta['TB Status']], dtype=int)
hiv_status = np.array([1 if v == 'HIV+' else 0 for v in meta['HIV_status']], dtype=int)
country    = meta['Country'].apply(lambda x: 'Viet Nam' if 'iet' in str(x) else x).values
age        = meta['Age'].values
gender     = meta['Gender'].values

# ── Load peptide categories & extract RRL ──────────────────────────────────
df_pep_cat = pd.read_excel(PEP_CAT_PATH)
rrl_sequences = set(
    df_pep_cat[df_pep_cat['Category'] == 'Random Reference Library (RRL2)']['Sequence']
)

# Filter signal matrix to RRL columns only
rrl_cols = [col for col in all_peptides if col in rrl_sequences]
idx_rrl  = np.array([all_peptides.index(c) for c in rrl_cols])

X_rrl = df_raw[rrl_cols].values.astype(np.float64)
rrl_peptide_names = rrl_cols
n_samples, n_rrl = X_rrl.shape

# Also keep mapping to origin names for annotation
seq2origin = dict(zip(df_pep_cat['Sequence'], df_pep_cat['Origin']))

print(f'\nRRL peptides found in signal matrix: {n_rrl}')
print(f'Samples: {n_samples}  (TB+={y_tb.sum()}, TB-={(y_tb==0).sum()})')
print(f'HIV+: {hiv_status.sum()}  HIV-: {(hiv_status==0).sum()}')
print(f'Countries: {pd.Series(country).value_counts().to_dict()}')
print(f'\nThis analysis uses ONLY the {n_rrl} RRL peptides (no TBC peptides).')

# %% [markdown]
## Shared Helper Functions

# %% [code]
def shannon_entropy(X):
    s = X.sum(axis=1, keepdims=True)
    s = np.where(s == 0, 1, s)
    p = np.clip(X / s, 1e-12, None)
    return -np.sum(p * np.log(p), axis=1)

def gini_coefficient(X):
    g = []
    for row in X:
        x  = np.sort(row)
        cx = np.cumsum(x)
        g.append((len(x)+1 - 2*cx.sum()/cx[-1])/len(x) if cx[-1] > 0 else 0)
    return np.array(g)

def build_Z(X_tr, X_te, y_tr, reg_lambda=1e-3):
    ent_tr = shannon_entropy(X_tr);  gin_tr = gini_coefficient(X_tr)
    ent_te = shannon_entropy(X_te);  gin_te = gini_coefficient(X_te)
    sc = StandardScaler()
    Xtr_sc = sc.fit_transform(X_tr);  Xte_sc = sc.transform(X_te)
    pca = PCA(n_components=2, random_state=RANDOM_STATE)
    pcs_tr = pca.fit_transform(Xtr_sc);  pcs_te = pca.transform(Xte_sc)
    ref  = pcs_tr[y_tr == 0]
    mu   = ref.mean(axis=0)
    cov  = np.cov(ref, rowvar=False) + reg_lambda * np.eye(2)
    ci   = np.linalg.inv(cov)
    mah  = lambda P: np.array([np.sqrt((x-mu)@ci@(x-mu)) for x in P])
    Z_tr_raw = np.column_stack([ent_tr, gin_tr, pcs_tr[:,0], pcs_tr[:,1], mah(pcs_tr)])
    Z_te_raw = np.column_stack([ent_te, gin_te, pcs_te[:,0], pcs_te[:,1], mah(pcs_te)])
    szZ = StandardScaler()
    return szZ.fit_transform(Z_tr_raw), szZ.transform(Z_te_raw), Z_tr_raw, Z_te_raw

def tune_and_fit_lr(X_tr, y_tr, inner_cv, C_GRID):
    gs = GridSearchCV(
        LogisticRegression(max_iter=2000, random_state=RANDOM_STATE, solver='lbfgs'),
        C_GRID, cv=inner_cv, scoring='roc_auc', n_jobs=-1)
    gs.fit(X_tr, y_tr)
    lr = LogisticRegression(C=gs.best_params_['C'], max_iter=2000,
                             random_state=RANDOM_STATE, solver='lbfgs')
    lr.fit(X_tr, y_tr)
    return lr, gs.best_params_['C']

def cliffs_delta(a, b):
    n1, n2 = len(a), len(b)
    more = sum(1 for x in a for y in b if x > y)
    less = sum(1 for x in a for y in b if x < y)
    return (more - less) / (n1 * n2)

def bootstrap_auc_ci(y_true, y_prob, n_boot=1000, ci=0.95, seed=42):
    rng  = np.random.default_rng(seed)
    aucs = []
    n    = len(y_true)
    for _ in range(n_boot):
        idx  = rng.integers(0, n, size=n)
        if len(np.unique(y_true[idx])) < 2: continue
        aucs.append(roc_auc_score(y_true[idx], y_prob[idx]))
    lo = np.percentile(aucs, (1-ci)/2*100)
    hi = np.percentile(aucs, (1-(1-ci)/2)*100)
    return lo, hi

def delong_test(y_true, prob_a, prob_b):
    def compute_Vxy(p_pos, p_neg):
        return np.array([np.mean(px > p_neg) for px in p_pos])
    def compute_Vyx(p_pos, p_neg):
        return np.array([np.mean(pn < p_pos) for pn in p_neg])
    y = np.asarray(y_true)
    pos_mask = y == 1;  neg_mask = y == 0
    n1, n0   = pos_mask.sum(), neg_mask.sum()
    def auc_vars(probs):
        pp, pn = probs[pos_mask], probs[neg_mask]
        V10    = compute_Vxy(pp, pn)
        V01    = compute_Vyx(pp, pn)
        auc    = V10.mean()
        s10    = np.var(V10, ddof=1) / n1
        s01    = np.var(V01, ddof=1) / n0
        return auc, s10, s01, V10, V01
    auc_a, s10a, s01a, V10a, V01a = auc_vars(prob_a)
    auc_b, s10b, s01b, V10b, V01b = auc_vars(prob_b)
    r_10 = np.corrcoef(V10a, V10b)[0,1] if n1 > 2 else 0
    r_01 = np.corrcoef(V01a, V01b)[0,1] if n0 > 2 else 0
    var_diff = s10a + s01a + s10b + s01b - 2*r_10*np.sqrt(s10a*s10b) - 2*r_01*np.sqrt(s01a*s01b)
    se = np.sqrt(max(var_diff, 1e-12))
    z  = (auc_a - auc_b) / se
    p  = 2 * (1 - norm.cdf(abs(z)))
    return float(z), float(p)

C_GRID   = {'C': [0.001, 0.01, 0.1, 1, 10, 100]}
outer_cv = StratifiedKFold(n_splits=5, shuffle=True, random_state=RANDOM_STATE)
inner_cv = StratifiedKFold(n_splits=5, shuffle=True, random_state=RANDOM_STATE)
TOP_N_PEPTIDES = 50

print('Helper functions defined')

# %% [markdown]
## STEP 3 — Model 1: Remodeling Classifier (on RRL only)

Remodeling Z-vector is now built **exclusively from 2,496 RRL peptides**.
This tests whether TB-related immune remodeling is detectable even in
a random/reference peptide library with no known TB epitopes.

# %% [code]
remod = dict(y_true=[], y_prob=[], fold_aucs=[], Z_train=[], Z_test=[],
             train_indices=[], test_indices=[])

for fold_i, (train_idx, test_idx) in enumerate(outer_cv.split(X_rrl, y_tb)):
    X_tr, X_te = X_rrl[train_idx], X_rrl[test_idx]
    y_tr, y_te = y_tb[train_idx],   y_tb[test_idx]

    Z_tr, Z_te, Z_tr_raw, _ = build_Z(X_tr, X_te, y_tr)
    lr, best_C = tune_and_fit_lr(Z_tr, y_tr, inner_cv, C_GRID)
    y_prob = lr.predict_proba(Z_te)[:, 1]

    remod['y_true'].extend(y_te.tolist())
    remod['y_prob'].extend(y_prob.tolist())
    remod['fold_aucs'].append(roc_auc_score(y_te, y_prob))
    remod['Z_train'].append(Z_tr_raw)
    remod['Z_test'].append(Z_te)
    remod['train_indices'].append(train_idx)
    remod['test_indices'].append(test_idx)
    print(f'  Remodeling Fold {fold_i+1}: AUC={remod["fold_aucs"][-1]:.3f}  C={best_C}')

print(f'\nRRL Remodeling AUC = {np.mean(remod["fold_aucs"]):.3f} +/- {np.std(remod["fold_aucs"]):.3f}')

# %% [markdown]
## STEP 4 — Model 2: Peptide-Level Classifier (RRL Only, Elastic Net → RF → LR)

### Two-stage feature reduction pipeline per fold:
1. **Standardize** intensities (training scaler only)
2. **Elastic Net pre-filter** (SGDClassifier, l1_ratio=0.5, α=0.001) → removes noise peptides
3. **Random Forest** feature importance on EN survivors → rank peptides
4. **Select top 50** peptides by RF importance
5. **Inner CV** to tune LR `C`
6. **Predict** on test set

### Why two-stage selection?
- With p >> n, direct RF ranking is unstable
- **Elastic Net** (L1+L2) sets uninformative coefficients to zero first
- RF then ranks only the surviving informative peptides
- Both EN and RF fitted on training fold only → no leakage

# %% [code]
peptide = dict(y_true=[], y_prob=[], fold_aucs=[], fold_top_peptides=[],
               fold_rf_importance=[], fold_best_C=[], fold_en_survivors=[])

# RF hyperparameters — fixed (no inner tuning for RF to save compute)
RF_PARAMS = dict(n_estimators=500, max_depth=None, min_samples_leaf=3,
                 class_weight='balanced', random_state=RANDOM_STATE, n_jobs=-1)

for fold_i, (train_idx, test_idx) in enumerate(outer_cv.split(X_rrl, y_tb)):
    X_tr_raw, X_te_raw = X_rrl[train_idx], X_rrl[test_idx]
    y_tr, y_te         = y_tb[train_idx],   y_tb[test_idx]

    # ── 4.1 Standardize ──────────────────────────────────────────────────
    scaler = StandardScaler()
    X_tr   = scaler.fit_transform(X_tr_raw)
    X_te   = scaler.transform(X_te_raw)

    # ── 4.2 Elastic Net pre-filter ───────────────────────────────────────
    en = SGDClassifier(loss='log_loss', penalty='elasticnet',
                       l1_ratio=0.5, alpha=0.001, max_iter=5000,
                       random_state=RANDOM_STATE, class_weight='balanced')
    en.fit(X_tr, y_tr)
    en_mask     = np.abs(en.coef_.ravel()) > 0
    n_survivors = en_mask.sum()
    peptide['fold_en_survivors'].append(n_survivors)

    X_tr_en = X_tr[:, en_mask]
    X_te_en = X_te[:, en_mask]

    # ── 4.3 Random Forest feature selection (on EN survivors) ─────
    rf = RandomForestClassifier(**RF_PARAMS)
    rf.fit(X_tr_en, y_tr)

    importance    = rf.feature_importances_
    ranked_idx    = np.argsort(importance)[::-1]
    top50_idx     = ranked_idx[:TOP_N_PEPTIDES]

    # Map back to original peptide names
    survivor_indices = np.where(en_mask)[0]
    top50_names = [rrl_peptide_names[survivor_indices[i]] for i in top50_idx]
    top50_imp   = importance[top50_idx]

    peptide['fold_top_peptides'].append(top50_names)
    peptide['fold_rf_importance'].append(
        {rrl_peptide_names[survivor_indices[i]]: importance[i]
         for i in range(len(survivor_indices))})

    # ── 4.4 Reduce to top 50 ─────────────────────────────────────────
    X_tr_sel = X_tr_en[:, top50_idx]
    X_te_sel = X_te_en[:, top50_idx]

    # ── 4.5 Inner CV + 4.6 Train LR ──────────────────────────────────
    lr, best_C = tune_and_fit_lr(X_tr_sel, y_tr, inner_cv, C_GRID)
    y_prob     = lr.predict_proba(X_te_sel)[:, 1]
    fold_auc   = roc_auc_score(y_te, y_prob)

    peptide['y_true'].extend(y_te.tolist())
    peptide['y_prob'].extend(y_prob.tolist())
    peptide['fold_aucs'].append(fold_auc)
    peptide['fold_best_C'].append(best_C)

    print(f'  Peptide  Fold {fold_i+1}: EN surv={n_survivors}/{X_tr.shape[1]} '
          f'({100*n_survivors/X_tr.shape[1]:.1f}%)  AUC={fold_auc:.3f}  '
          f'C={best_C}  top3={top50_names[:3]}')

print(f'\nPeptide AUC = {np.mean(peptide["fold_aucs"]):.3f} ± {np.std(peptide["fold_aucs"]):.3f}')
print(f'EN survivors per fold: {peptide["fold_en_survivors"]}')

# %% [markdown]
### 4B — RRL Peptide Selection Consensus

# %% [code]
all_selected = [p for fold in peptide['fold_top_peptides'] for p in fold]
selection_freq = Counter(all_selected)

df_consensus = pd.DataFrame([
    {'Peptide': pep,
     'Origin': seq2origin.get(pep, ''),
     'Folds_Selected': cnt,
     'Mean_RF_Importance': np.mean([
         peptide['fold_rf_importance'][f].get(pep, 0) for f in range(5)])}
    for pep, cnt in selection_freq.most_common()
]).sort_values(['Folds_Selected', 'Mean_RF_Importance'], ascending=[False, False])

core_peptides    = df_consensus[df_consensus['Folds_Selected'] == 5]['Peptide'].tolist()
robust_peptides  = df_consensus[df_consensus['Folds_Selected'] >= 4]['Peptide'].tolist()
stable_peptides  = df_consensus[df_consensus['Folds_Selected'] >= 3]['Peptide'].tolist()

print(f'Total unique RRL peptides ever selected: {len(df_consensus)}')
print(f'Core     (5/5 folds): {len(core_peptides)}')
print(f'Robust   (>=4/5 folds): {len(robust_peptides)}')
print(f'Stable   (>=3/5 folds): {len(stable_peptides)}')
print(f'\nTop 20 most consistently selected RRL peptides:')
print(df_consensus.head(20).to_string(index=False, float_format='%.5f'))

df_consensus.to_csv('rrl_peptide_selection_consensus.csv', index=False)
print('\nSaved: rrl_peptide_selection_consensus.csv')

# %% [markdown]
### 4C — Peptide Number Sweep (RRL Only)
How many top RRL peptides give the best AUC?

# %% [code]
SWEEP_NS = [1, 2, 3, 4, 5, 10, 15, 20, 30, 50, 75, 100, 150, 200]
sweep_results = {n: [] for n in SWEEP_NS}

print('Running peptide number sweep (EN → RF → LR)...')
for fold_i, (train_idx, test_idx) in enumerate(outer_cv.split(X_rrl, y_tb)):
    X_tr_raw, X_te_raw = X_rrl[train_idx], X_rrl[test_idx]
    y_tr, y_te         = y_tb[train_idx],   y_tb[test_idx]

    scaler = StandardScaler()
    X_tr   = scaler.fit_transform(X_tr_raw)
    X_te   = scaler.transform(X_te_raw)

    # Elastic Net pre-filter
    en = SGDClassifier(loss='log_loss', penalty='elasticnet',
                       l1_ratio=0.5, alpha=0.001, max_iter=5000,
                       random_state=RANDOM_STATE, class_weight='balanced')
    en.fit(X_tr, y_tr)
    en_mask = np.abs(en.coef_.ravel()) > 0
    X_tr_en = X_tr[:, en_mask]
    X_te_en = X_te[:, en_mask]

    # Fit RF once per fold on EN survivors
    rf = RandomForestClassifier(**RF_PARAMS)
    rf.fit(X_tr_en, y_tr)
    ranked = np.argsort(rf.feature_importances_)[::-1]

    for N in SWEEP_NS:
        top_idx  = ranked[:min(N, len(ranked))]
        gs = GridSearchCV(
            LogisticRegression(max_iter=2000, random_state=RANDOM_STATE, solver='lbfgs'),
            C_GRID, cv=inner_cv, scoring='roc_auc', n_jobs=-1)
        gs.fit(X_tr_en[:, top_idx], y_tr)
        lr = LogisticRegression(C=gs.best_params_['C'], max_iter=2000,
                                 random_state=RANDOM_STATE, solver='lbfgs')
        lr.fit(X_tr_en[:, top_idx], y_tr)
        auc = roc_auc_score(y_te, lr.predict_proba(X_te_en[:, top_idx])[:,1])
        sweep_results[N].append(auc)

    print(f'  Fold {fold_i+1} done')

sweep_means = [np.mean(sweep_results[n]) for n in SWEEP_NS]
sweep_stds  = [np.std(sweep_results[n])  for n in SWEEP_NS]
best_N      = SWEEP_NS[np.argmax(sweep_means)]
best_auc    = max(sweep_means)

print(f'\nOptimal N = {best_N} peptides (AUC = {best_auc:.3f})')
for n, m, s in zip(SWEEP_NS, sweep_means, sweep_stds):
    marker = ' ← BEST' if n == best_N else ''
    print(f'  N={n:4d}: {m:.3f} ± {s:.3f}{marker}')

# %% [markdown]
## STEP 5 — Meta-Model (Stacked Classifier)

Combine remodeling + peptide probabilities. Keep only if ΔAUC >= 0.02.

# %% [code]
meta_model = dict(y_true=[], y_prob=[], fold_aucs=[])

fold_splits = list(outer_cv.split(X_rrl, y_tb))

for fold_i, (train_idx, test_idx) in enumerate(fold_splits):
    y_tr, y_te = y_tb[train_idx], y_tb[test_idx]

    # Remodeling OOF on training
    Z_tr, Z_te, _, _ = build_Z(X_rrl[train_idx], X_rrl[test_idx], y_tr)
    oof_remod_tr = cross_val_predict(
        LogisticRegression(C=1, max_iter=2000, random_state=RANDOM_STATE, solver='lbfgs'),
        Z_tr, y_tr, cv=inner_cv, method='predict_proba')[:, 1]

    # Peptide OOF on training
    scaler = StandardScaler()
    X_tr_sc = scaler.fit_transform(X_rrl[train_idx])
    X_te_sc = scaler.transform(X_rrl[test_idx])
    en = SGDClassifier(loss='log_loss', penalty='elasticnet',
                       l1_ratio=0.5, alpha=0.001, max_iter=5000,
                       random_state=RANDOM_STATE, class_weight='balanced')
    en.fit(X_tr_sc, y_tr)
    en_mask = np.abs(en.coef_.ravel()) > 0
    X_tr_sc = X_tr_sc[:, en_mask]; X_te_sc = X_te_sc[:, en_mask]
    rf = RandomForestClassifier(**RF_PARAMS)
    rf.fit(X_tr_sc, y_tr)
    top_idx = np.argsort(rf.feature_importances_)[::-1][:TOP_N_PEPTIDES]
    oof_pep_tr = cross_val_predict(
        LogisticRegression(C=1, max_iter=2000, random_state=RANDOM_STATE, solver='lbfgs'),
        X_tr_sc[:, top_idx], y_tr, cv=inner_cv, method='predict_proba')[:, 1]

    # Stack
    S_tr = np.column_stack([oof_remod_tr, oof_pep_tr])
    meta_lr, best_C = tune_and_fit_lr(S_tr, y_tr, inner_cv,
                                       {'C': [0.01, 0.1, 1, 10]})

    # Test predictions
    lr_remod, _ = tune_and_fit_lr(Z_tr, y_tr, inner_cv, C_GRID)
    p_remod_te  = lr_remod.predict_proba(Z_te)[:, 1]

    lr_pep, _   = tune_and_fit_lr(X_tr_sc[:, top_idx], y_tr, inner_cv, C_GRID)
    p_pep_te    = lr_pep.predict_proba(X_te_sc[:, top_idx])[:, 1]

    S_te    = np.column_stack([p_remod_te, p_pep_te])
    p_meta  = meta_lr.predict_proba(S_te)[:, 1]
    fold_auc = roc_auc_score(y_te, p_meta)

    meta_model['y_true'].extend(y_te.tolist())
    meta_model['y_prob'].extend(p_meta.tolist())
    meta_model['fold_aucs'].append(fold_auc)
    print(f'  Meta Fold {fold_i+1}: AUC={fold_auc:.3f}')

auc_remod = np.mean(remod['fold_aucs'])
auc_pep   = np.mean(peptide['fold_aucs'])
auc_meta  = np.mean(meta_model['fold_aucs'])
best_base = max(auc_remod, auc_pep)
delta_meta = auc_meta - best_base

KEEP_META = delta_meta >= 0.02
print(f'\nRemodeling AUC : {auc_remod:.3f}')
print(f'Peptide    AUC : {auc_pep:.3f}')
print(f'Meta       AUC : {auc_meta:.3f}  (delta = {delta_meta:+.3f})')
print(f'Keep meta-model: {"YES" if KEEP_META else "NO -- improvement < 0.02"}')


# %% [markdown]
## STEP 6 — HIV-Stratified Analysis (KEY COMPARISON)

**Central question:** How does AUC change when restricting to HIV-negative samples
using only the RRL peptide library?

# %% [code]
# ── 6A: HIV- only pipeline on RRL ────────────────────────────────────────
hiv_neg_mask = (hiv_status == 0)
X_hiv_neg    = X_rrl[hiv_neg_mask]
y_hiv_neg    = y_tb[hiv_neg_mask]

print(f'HIV- cohort: {hiv_neg_mask.sum()} samples  '
      f'(TB+={y_hiv_neg.sum()}, TB-={(y_hiv_neg==0).sum()})')

hiv_neg_remod   = dict(fold_aucs=[], y_true=[], y_prob=[])
hiv_neg_peptide = dict(fold_aucs=[], y_true=[], y_prob=[])

cv_hiv = StratifiedKFold(n_splits=5, shuffle=True, random_state=RANDOM_STATE)

for fold_i, (tr_idx, te_idx) in enumerate(cv_hiv.split(X_hiv_neg, y_hiv_neg)):
    Xtr, Xte = X_hiv_neg[tr_idx], X_hiv_neg[te_idx]
    ytr, yte = y_hiv_neg[tr_idx], y_hiv_neg[te_idx]

    # Remodeling on RRL
    Ztr, Zte, _, _ = build_Z(Xtr, Xte, ytr)
    lr, _ = tune_and_fit_lr(Ztr, ytr, inner_cv, C_GRID)
    p = lr.predict_proba(Zte)[:, 1]
    hiv_neg_remod['fold_aucs'].append(roc_auc_score(yte, p))
    hiv_neg_remod['y_true'].extend(yte.tolist())
    hiv_neg_remod['y_prob'].extend(p.tolist())

    # Peptide on RRL
    sc = StandardScaler()
    Xtr_sc = sc.fit_transform(Xtr); Xte_sc = sc.transform(Xte)
    en = SGDClassifier(loss='log_loss', penalty='elasticnet',
                       l1_ratio=0.5, alpha=0.001, max_iter=5000,
                       random_state=RANDOM_STATE, class_weight='balanced')
    en.fit(Xtr_sc, ytr)
    en_mask = np.abs(en.coef_.ravel()) > 0
    Xtr_en = Xtr_sc[:, en_mask]; Xte_en = Xte_sc[:, en_mask]
    rf = RandomForestClassifier(**RF_PARAMS)
    rf.fit(Xtr_en, ytr)
    top_idx = np.argsort(rf.feature_importances_)[::-1][:TOP_N_PEPTIDES]
    lr2, _ = tune_and_fit_lr(Xtr_en[:, top_idx], ytr, inner_cv, C_GRID)
    p2 = lr2.predict_proba(Xte_en[:, top_idx])[:, 1]
    hiv_neg_peptide['fold_aucs'].append(roc_auc_score(yte, p2))
    hiv_neg_peptide['y_true'].extend(yte.tolist())
    hiv_neg_peptide['y_prob'].extend(p2.tolist())

# ── 6B: HIV as covariate ──────────────────────────────────────────────────
covar_aucs = []
for fold_i, (train_idx, test_idx) in enumerate(outer_cv.split(X_rrl, y_tb)):
    y_tr, y_te = y_tb[train_idx], y_tb[test_idx]
    Z_tr, Z_te, _, _ = build_Z(X_rrl[train_idx], X_rrl[test_idx], y_tr)
    hiv_tr = hiv_status[train_idx].reshape(-1,1)
    hiv_te = hiv_status[test_idx].reshape(-1,1)
    Z_tr_h = np.hstack([Z_tr, hiv_tr])
    Z_te_h = np.hstack([Z_te, hiv_te])
    lr, _ = tune_and_fit_lr(Z_tr_h, y_tr, inner_cv, C_GRID)
    covar_aucs.append(roc_auc_score(y_te, lr.predict_proba(Z_te_h)[:,1]))

print('\n' + '='*65)
print('HIV ANALYSIS RESULTS — RRL LIBRARY')
print('='*65)
print(f'Full cohort   | Remodeling: {np.mean(remod["fold_aucs"]):.3f}  '
      f'Peptide: {np.mean(peptide["fold_aucs"]):.3f}')
print(f'HIV- only     | Remodeling: {np.mean(hiv_neg_remod["fold_aucs"]):.3f}  '
      f'Peptide: {np.mean(hiv_neg_peptide["fold_aucs"]):.3f}')
print(f'HIV covariate | Remodeling+HIV: {np.mean(covar_aucs):.3f}')
print(f'\nDelta AUC (HIV- vs Full) | Remodeling: '
      f'{np.mean(hiv_neg_remod["fold_aucs"]) - np.mean(remod["fold_aucs"]):+.3f}')
print(f'Delta AUC (HIV- vs Full) | Peptide   : '
      f'{np.mean(hiv_neg_peptide["fold_aucs"]) - np.mean(peptide["fold_aucs"]):+.3f}')


# %% [markdown]
## STEP 7 — Effect Size Analysis (RRL)

# %% [code]
Z_LABELS = ['Entropy', 'Gini', 'PC1', 'PC2', 'Mahalanobis']

fold_cliffs  = {z: [] for z in Z_LABELS}
fold_mwu_p   = {z: [] for z in Z_LABELS}

for fold_i, (train_idx, _) in enumerate(outer_cv.split(X_rrl, y_tb)):
    y_tr = y_tb[train_idx]
    _, _, Z_tr_raw, _ = build_Z(X_rrl[train_idx], X_rrl[train_idx[:5]], y_tr)

    for fi, fname in enumerate(Z_LABELS):
        vals_neg = Z_tr_raw[y_tr == 0, fi]
        vals_pos = Z_tr_raw[y_tr == 1, fi]
        stat, p = mannwhitneyu(vals_pos, vals_neg, alternative='two-sided')
        fold_mwu_p[fname].append(p)
        cd = cliffs_delta(vals_pos.tolist(), vals_neg.tolist())
        fold_cliffs[fname].append(cd)

def cliff_label(d):
    a = abs(d)
    if a < 0.15: return 'negligible'
    elif a < 0.33: return 'small'
    elif a < 0.47: return 'medium'
    else: return 'large'

rows = []
for fname in Z_LABELS:
    cd_mean = np.mean(fold_cliffs[fname])
    rows.append({'Feature': fname, "Cliff's delta (mean)": cd_mean,
                 "Cliff's delta (std)": np.std(fold_cliffs[fname]),
                 '|delta|': abs(cd_mean), 'Magnitude': cliff_label(cd_mean),
                 'MWU p (mean)': np.mean(fold_mwu_p[fname])})

df_effect = pd.DataFrame(rows).sort_values('|delta|', ascending=False)
print("Effect Sizes (Cliff's Delta, TB+ vs TB-, RRL only, training folds):")
print(df_effect.to_string(index=False, float_format='%.4f'))
df_effect.to_csv('rrl_results_effect_sizes.csv', index=False)

# %% [markdown]
## STEP 8 — Volcano Plot (RRL Peptides)

# %% [code]
fold_log2fc = []
fold_pvals  = []

for train_idx, _ in outer_cv.split(X_rrl, y_tb):
    X_tr = X_rrl[train_idx]
    y_tr = y_tb[train_idx]
    pos_tr = X_tr[y_tr == 1]
    neg_tr = X_tr[y_tr == 0]

    mean_pos = pos_tr.mean(axis=0) + 1e-9
    mean_neg = neg_tr.mean(axis=0) + 1e-9
    log2fc   = np.log2(mean_pos / mean_neg)
    fold_log2fc.append(log2fc)

    pvals = []
    for j in range(X_tr.shape[1]):
        _, p = mannwhitneyu(pos_tr[:, j], neg_tr[:, j], alternative='two-sided')
        pvals.append(p)
    fold_pvals.append(pvals)

mean_log2fc = np.mean(fold_log2fc, axis=0)
mean_pvals  = np.mean(fold_pvals, axis=0)

_, fdr_pvals, _, _ = multipletests(mean_pvals, alpha=0.05, method='fdr_bh')
neg_log10_fdr = -np.log10(fdr_pvals + 1e-300)

df_volcano = pd.DataFrame({
    'Peptide': rrl_peptide_names,
    'Origin': [seq2origin.get(p, '') for p in rrl_peptide_names],
    'log2FC': mean_log2fc,
    'FDR_p': fdr_pvals,
    'neg_log10_FDR': neg_log10_fdr,
})
df_volcano['Significant'] = (fdr_pvals < 0.05) & (np.abs(mean_log2fc) > 0.5)
df_volcano['Direction']   = np.where(mean_log2fc > 0, 'Up in TB+', 'Up in TB-')

n_sig = df_volcano['Significant'].sum()
n_up  = ((df_volcano['Significant']) & (mean_log2fc > 0)).sum()
n_dn  = ((df_volcano['Significant']) & (mean_log2fc < 0)).sum()

print(f'RRL significant peptides (FDR<0.05, |log2FC|>0.5): {n_sig}')
print(f'  Up in TB+: {n_up}  |  Up in TB-: {n_dn}')
print(f'\nTop 15 upregulated in TB+ (RRL):')
print(df_volcano[df_volcano['log2FC']>0].sort_values('neg_log10_FDR',
    ascending=False).head(15)[['Peptide','Origin','log2FC','FDR_p']].to_string(
    index=False, float_format='%.4f'))
df_volcano.to_csv('rrl_peptide_volcano_data.csv', index=False)
print('\nSaved: rrl_peptide_volcano_data.csv')

# %% [markdown]
## STEP 9 — Bootstrap CI + DeLong Test

# %% [code]
y_true_arr = np.array(remod['y_true'])
p_remod    = np.array(remod['y_prob'])
p_peptide  = np.array(peptide['y_prob'])
p_meta_arr = np.array(meta_model['y_prob'])

ci_remod   = bootstrap_auc_ci(y_true_arr, p_remod)
ci_peptide = bootstrap_auc_ci(y_true_arr, p_peptide)
ci_meta    = bootstrap_auc_ci(y_true_arr, p_meta_arr)

z_rp, p_rp = delong_test(y_true_arr, p_remod, p_peptide)
best_base_prob = p_remod if auc_remod >= auc_pep else p_peptide
z_mb, p_mb     = delong_test(y_true_arr, p_meta_arr, best_base_prob)

y_hivneg_arr    = np.array(hiv_neg_remod['y_true'])
p_hivneg_remod  = np.array(hiv_neg_remod['y_prob'])
p_hivneg_pep    = np.array(hiv_neg_peptide['y_prob'])
ci_hivneg_remod = bootstrap_auc_ci(y_hivneg_arr, p_hivneg_remod)
ci_hivneg_pep   = bootstrap_auc_ci(y_hivneg_arr, p_hivneg_pep)

print('='*65)
print('RRL LIBRARY — FINAL RESULTS SUMMARY')
print('='*65)
print("\n{:<30} {:>10} {:>20} {:>10}".format("Model", "Mean AUC", "95% CI", "Fold SD"))
print('-'*65)

rows_out = [
    ('Remodeling (full)',       np.mean(remod['fold_aucs']),         ci_remod,   np.std(remod['fold_aucs'])),
    ('Peptide LR (full)',       np.mean(peptide['fold_aucs']),       ci_peptide, np.std(peptide['fold_aucs'])),
    ('Meta-model',              np.mean(meta_model['fold_aucs']),    ci_meta,    np.std(meta_model['fold_aucs'])),
    ('Remodeling (HIV- only)',  np.mean(hiv_neg_remod['fold_aucs']), ci_hivneg_remod, np.std(hiv_neg_remod['fold_aucs'])),
    ('Peptide LR (HIV- only)',  np.mean(hiv_neg_peptide['fold_aucs']),ci_hivneg_pep,  np.std(hiv_neg_peptide['fold_aucs'])),
    ('Remodeling + HIV covar.', np.mean(covar_aucs), ('n/a','n/a'), np.std(covar_aucs)),
]

df_final = []
for name, auc, ci, sd in rows_out:
    ci_str = f'{ci[0]:.3f}-{ci[1]:.3f}' if ci[0] != 'n/a' else 'n/a'
    print(f'{name:<30} {auc:>10.3f} {ci_str:>20} {sd:>10.3f}')
    df_final.append({'Model': name, 'AUC': auc,
                     'CI_lo': ci[0] if ci[0]!='n/a' else np.nan,
                     'CI_hi': ci[1] if ci[1]!='n/a' else np.nan, 'SD': sd})

print(f'\nDeLong: Remodeling vs Peptide  z={z_rp:.3f}  p={p_rp:.4f}')
print(f'DeLong: Meta vs Best-base      z={z_mb:.3f}  p={p_mb:.4f}')
print(f'Meta-model decision: {"KEEP" if KEEP_META else "DISCARD (delta < 0.02)"}')

pd.DataFrame(df_final).to_csv('rrl_results_final_summary.csv', index=False)
print('\nSaved: rrl_results_final_summary.csv')

# %% [markdown]
## STEP 10 — Publication Figures

# %% [code]
# ═══════════════════════════════════════════════════════════════════════════
# FIGURE 1 — AUC Comparison: Remodeling vs Peptide vs Meta (RRL)
# ═══════════════════════════════════════════════════════════════════════════
fig, axes = plt.subplots(1, 3, figsize=(17, 6))

COLORS_MODEL = {'Remodeling': '#2196F3', 'Peptide LR': '#4CAF50', 'Meta-model': '#FF9800'}

# Left: Fold AUC box + swarm
ax = axes[0]
models_data = [
    ('Remodeling',  remod['fold_aucs']),
    ('Peptide LR',  peptide['fold_aucs']),
    ('Meta-model',  meta_model['fold_aucs']),
]
for xi, (name, aucs) in enumerate(models_data):
    color = COLORS_MODEL[name]
    bp = ax.boxplot(aucs, positions=[xi], patch_artist=True, widths=0.4,
                    medianprops={'color':'black','linewidth':2})
    bp['boxes'][0].set_facecolor(color); bp['boxes'][0].set_alpha(0.6)
    jitter = np.random.normal(0, 0.04, len(aucs))
    ax.scatter(np.full(5, xi)+jitter, aucs, color=color, s=50, zorder=5,
               edgecolors='white')
ax.axhline(0.5, color='red', ls='--', alpha=0.4, lw=1.5)
ax.set_xticks([0,1,2]); ax.set_xticklabels([m[0] for m in models_data])
ax.set_ylabel('ROC-AUC'); ax.set_ylim(0.2, 1.0)
ax.set_title('A  |  Fold AUC Distribution (RRL)', fontweight='bold')

# Middle: ROC curves
ax2 = axes[1]
for name, y_t, y_p, color in [
    ('Remodeling', remod['y_true'],      remod['y_prob'],       '#2196F3'),
    ('Peptide LR', peptide['y_true'],    peptide['y_prob'],     '#4CAF50'),
    ('Meta-model', meta_model['y_true'], meta_model['y_prob'],  '#FF9800'),
]:
    yt = np.array(y_t); yp = np.array(y_p)
    fpr, tpr, _ = roc_curve(yt, yp)
    auc_val = roc_auc_score(yt, yp)
    ci_lo, ci_hi = bootstrap_auc_ci(yt, yp, n_boot=500)
    ax2.plot(fpr, tpr, color=color, lw=2.5,
             label=f'{name}\nAUC={auc_val:.3f} [{ci_lo:.3f}-{ci_hi:.3f}]')
ax2.plot([0,1],[0,1],'k--',alpha=0.3, lw=1.5)
ax2.set_xlabel('False Positive Rate'); ax2.set_ylabel('True Positive Rate')
ax2.set_title('B  |  ROC Curves (RRL, bootstrap CI)', fontweight='bold')
ax2.legend(loc='lower right', fontsize=8)

# Right: Peptide sweep
ax3 = axes[2]
ax3.errorbar(SWEEP_NS, sweep_means, yerr=sweep_stds, fmt='o-',
             color='#4CAF50', linewidth=2, markersize=7, capsize=5,
             label='Peptide LR AUC')
ax3.axhline(np.mean(remod['fold_aucs']), color='#2196F3', ls='--',
            lw=2, label=f'Remodeling AUC = {np.mean(remod["fold_aucs"]):.3f}')
ax3.axvline(best_N, color='red', ls=':', lw=1.5, label=f'Optimal N={best_N}')
ax3.scatter([best_N], [best_auc], color='red', s=120, zorder=6)
ax3.set_xlabel('Number of Top RRL Peptides')
ax3.set_ylabel('ROC-AUC')
ax3.set_title(f'C  |  RRL Peptide Sweep\nOptimal N={best_N} (AUC={best_auc:.3f})',
              fontweight='bold')
ax3.legend(fontsize=9); ax3.set_ylim(0.35, 1.0)

fig.suptitle('Figure 1 — RRL Library: Model Performance Comparison',
             fontsize=13, fontweight='bold')
plt.tight_layout()
plt.savefig('RRL_Fig1_auc_comparison.png', bbox_inches='tight', dpi=180)
plt.show()
print('Fig 1 saved.')

# %% [code]
# ═══════════════════════════════════════════════════════════════════════════
# FIGURE 2 — HIV Effect on RRL-Based TB Classification
# ═══════════════════════════════════════════════════════════════════════════
fig, axes = plt.subplots(1, 3, figsize=(17, 6))

# Left: AUC bars — Full vs HIV-
ax = axes[0]
bar_data = [
    ('Remodeling\n(Full)',  np.mean(remod['fold_aucs']),          np.std(remod['fold_aucs']),          '#2196F3'),
    ('Remodeling\n(HIV-)',  np.mean(hiv_neg_remod['fold_aucs']),  np.std(hiv_neg_remod['fold_aucs']),  '#90CAF9'),
    ('Peptide LR\n(Full)',  np.mean(peptide['fold_aucs']),        np.std(peptide['fold_aucs']),        '#4CAF50'),
    ('Peptide LR\n(HIV-)',  np.mean(hiv_neg_peptide['fold_aucs']),np.std(hiv_neg_peptide['fold_aucs']),'#A5D6A7'),
    ('Remod\n+HIV covar',   np.mean(covar_aucs),                  np.std(covar_aucs),                  '#FF9800'),
]
for xi, (name, m, s, color) in enumerate(bar_data):
    ax.bar(xi, m, yerr=s, color=color, edgecolor='black', alpha=0.8, capsize=6)
    ax.text(xi, m+s+0.01, f'{m:.3f}', ha='center', va='bottom', fontsize=9, fontweight='bold')
ax.axhline(0.5, color='red', ls='--', alpha=0.4)
ax.set_xticks(range(len(bar_data)))
ax.set_xticklabels([b[0] for b in bar_data], fontsize=8)
ax.set_ylabel('Mean AUC +/- SD'); ax.set_ylim(0.3, 1.0)
ax.set_title('A  |  Full Cohort vs HIV- Only (RRL)', fontweight='bold')

# Middle: ROC overlay — Full vs HIV- (Remodeling)
ax2 = axes[1]
fpr_f, tpr_f, _ = roc_curve(remod['y_true'], remod['y_prob'])
auc_f = roc_auc_score(remod['y_true'], remod['y_prob'])
ax2.plot(fpr_f, tpr_f, color='#2196F3', lw=2.5, label=f'Full (AUC={auc_f:.3f})')
fpr_h, tpr_h, _ = roc_curve(hiv_neg_remod['y_true'], hiv_neg_remod['y_prob'])
auc_h = roc_auc_score(hiv_neg_remod['y_true'], hiv_neg_remod['y_prob'])
ax2.plot(fpr_h, tpr_h, color='#90CAF9', lw=2.5, ls='--', label=f'HIV- (AUC={auc_h:.3f})')
ax2.plot([0,1],[0,1],'k--',alpha=0.3)
ax2.set_xlabel('FPR'); ax2.set_ylabel('TPR')
ax2.set_title('B  |  ROC: Remodeling (RRL)\nFull vs HIV-', fontweight='bold')
ax2.legend(fontsize=9)

# Right: ROC overlay — Full vs HIV- (Peptide)
ax3 = axes[2]
fpr_pf, tpr_pf, _ = roc_curve(peptide['y_true'], peptide['y_prob'])
auc_pf = roc_auc_score(peptide['y_true'], peptide['y_prob'])
ax3.plot(fpr_pf, tpr_pf, color='#4CAF50', lw=2.5, label=f'Full (AUC={auc_pf:.3f})')
fpr_ph, tpr_ph, _ = roc_curve(hiv_neg_peptide['y_true'], hiv_neg_peptide['y_prob'])
auc_ph = roc_auc_score(hiv_neg_peptide['y_true'], hiv_neg_peptide['y_prob'])
ax3.plot(fpr_ph, tpr_ph, color='#A5D6A7', lw=2.5, ls='--', label=f'HIV- (AUC={auc_ph:.3f})')
ax3.plot([0,1],[0,1],'k--',alpha=0.3)
ax3.set_xlabel('FPR'); ax3.set_ylabel('TPR')
ax3.set_title('C  |  ROC: Peptide LR (RRL)\nFull vs HIV-', fontweight='bold')
ax3.legend(fontsize=9)

fig.suptitle('Figure 2 — HIV Effect on RRL-Based TB Classification',
             fontsize=13, fontweight='bold')
plt.tight_layout()
plt.savefig('RRL_Fig2_HIV_effect.png', bbox_inches='tight', dpi=180)
plt.show()
print('Fig 2 saved.')

# %% [code]
# ═══════════════════════════════════════════════════════════════════════════
# FIGURE 3 — Volcano + Peptide Consensus (RRL)
# ═══════════════════════════════════════════════════════════════════════════
fig, axes = plt.subplots(1, 2, figsize=(17, 7))

# Left: Volcano
ax = axes[0]
mask_ns = ~df_volcano['Significant']
ax.scatter(df_volcano.loc[mask_ns,'log2FC'], df_volcano.loc[mask_ns,'neg_log10_FDR'],
           c='#B0BEC5', s=12, alpha=0.4, label='Not significant')

for direction, color, label in [('Up in TB+','#F44336','Up in TB+'),
                                  ('Up in TB-','#2196F3','Up in TB-')]:
    mask = df_volcano['Significant'] & (df_volcano['Direction'] == direction)
    ax.scatter(df_volcano.loc[mask,'log2FC'], df_volcano.loc[mask,'neg_log10_FDR'],
               c=color, s=30, alpha=0.75, label=label)

# Highlight core consensus
df_volcano['selected_consensus'] = df_volcano['Peptide'].isin(core_peptides)
mask_core = df_volcano['Significant'] & df_volcano['selected_consensus']
if mask_core.any():
    ax.scatter(df_volcano.loc[mask_core,'log2FC'], df_volcano.loc[mask_core,'neg_log10_FDR'],
               c='gold', s=80, edgecolors='black', linewidths=0.8, zorder=6,
               label=f'Core consensus (5/5, n={len(core_peptides)})')

fdr_thresh = -np.log10(0.05)
ax.axhline(fdr_thresh, color='gray', ls='--', lw=1, alpha=0.7)
ax.axvline(0.5, color='gray', ls=':', lw=1, alpha=0.7)
ax.axvline(-0.5, color='gray', ls=':', lw=1, alpha=0.7)

top_label = df_volcano[df_volcano['Significant']].nlargest(10,'neg_log10_FDR')
for _, row in top_label.iterrows():
    ax.annotate(row['Peptide'], (row['log2FC'], row['neg_log10_FDR']),
                xytext=(5, 3), textcoords='offset points', fontsize=6.5,
                arrowprops=dict(arrowstyle='-', lw=0.5, color='gray'))

ax.set_xlabel('log2 Fold Change (TB+ / TB-)'); ax.set_ylabel('-log10(FDR)')
ax.set_title(f'A  |  RRL Volcano: {n_sig} significant\n(FDR<0.05, |log2FC|>0.5)',
             fontweight='bold')
ax.legend(fontsize=8, loc='upper left')

# Right: Consensus bar chart
ax2 = axes[1]
df_top_con = df_consensus.head(30).copy()
bar_colors_con = []
for _, row in df_top_con.iterrows():
    n_folds = row['Folds_Selected']
    if n_folds == 5:   bar_colors_con.append('#F44336')
    elif n_folds == 4: bar_colors_con.append('#FF9800')
    elif n_folds == 3: bar_colors_con.append('#FFC107')
    else:              bar_colors_con.append('#90CAF9')

ax2.barh(range(len(df_top_con)), df_top_con['Mean_RF_Importance'],
         color=bar_colors_con, edgecolor='black', alpha=0.85)
ax2.set_yticks(range(len(df_top_con)))
ax2.set_yticklabels(df_top_con['Peptide'], fontsize=7)
ax2.invert_yaxis()
ax2.set_xlabel('Mean RF Importance')
ax2.set_title('B  |  Top 30 RRL Peptides by RF Consensus', fontweight='bold')

for i, (_, row) in enumerate(df_top_con.iterrows()):
    ax2.text(row['Mean_RF_Importance'] * 1.01, i,
             f"{int(row['Folds_Selected'])}/5", va='center', fontsize=7.5)

from matplotlib.patches import Patch
legend_els = [Patch(facecolor='#F44336', label='5/5 folds (core)'),
              Patch(facecolor='#FF9800', label='4/5 folds (robust)'),
              Patch(facecolor='#FFC107', label='3/5 folds (stable)'),
              Patch(facecolor='#90CAF9', label='<=2/5 folds')]
ax2.legend(handles=legend_els, fontsize=8, loc='lower right')

fig.suptitle('Figure 3 — RRL Peptide Signatures: Volcano & Selection Consensus',
             fontsize=13, fontweight='bold')
plt.tight_layout()
plt.savefig('RRL_Fig3_volcano_consensus.png', bbox_inches='tight', dpi=180)
plt.show()
print('Fig 3 saved.')

# %% [markdown]
## Final Summary & Export

# %% [code]
print('='*70)
print('RRL LIBRARY PIPELINE -- FINAL SUMMARY')
print('='*70)

print(f"""
COHORT
  Samples  : {n_samples}  (TB+={y_tb.sum()}, TB-={(y_tb==0).sum()})
  HIV+     : {hiv_status.sum()}  |  HIV-: {(hiv_status==0).sum()}
  Peptides : {n_rrl} RRL (Random Reference Library)

MODEL PERFORMANCE (nested 5-fold CV, mean AUC +/- SD)
  Remodeling (RRL)     : {np.mean(remod['fold_aucs']):.3f} +/- {np.std(remod['fold_aucs']):.3f}
  Peptide LR (RRL)     : {np.mean(peptide['fold_aucs']):.3f} +/- {np.std(peptide['fold_aucs']):.3f}
  Meta-model (RRL)     : {np.mean(meta_model['fold_aucs']):.3f} +/- {np.std(meta_model['fold_aucs']):.3f}
  Meta kept            : {'YES' if KEEP_META else 'NO (delta < 0.02)'}

PEPTIDE SELECTION (top {TOP_N_PEPTIDES} per fold via RF)
  Core (5/5 folds)     : {len(core_peptides)}
  Robust (>=4/5)       : {len(robust_peptides)}
  Stable (>=3/5)       : {len(stable_peptides)}
  Optimal N (sweep)    : {best_N} peptides (AUC={best_auc:.3f})

HIV ANALYSIS -- KEY RESULTS FOR RRL
  Remodeling Full      : {np.mean(remod['fold_aucs']):.3f}
  Remodeling HIV-      : {np.mean(hiv_neg_remod['fold_aucs']):.3f}  (delta={np.mean(hiv_neg_remod['fold_aucs'])-np.mean(remod['fold_aucs']):+.3f})
  Peptide LR Full      : {np.mean(peptide['fold_aucs']):.3f}
  Peptide LR HIV-      : {np.mean(hiv_neg_peptide['fold_aucs']):.3f}  (delta={np.mean(hiv_neg_peptide['fold_aucs'])-np.mean(peptide['fold_aucs']):+.3f})
  Remodeling + HIV cov.: {np.mean(covar_aucs):.3f}

STATISTICAL COMPARISONS
  DeLong: Remodeling vs Peptide  z={z_rp:.3f}  p={p_rp:.4f}
  DeLong: Meta vs Best-base      z={z_mb:.3f}  p={p_mb:.4f}
""")

print("EFFECT SIZES (Cliff's delta, RRL, training folds):")
print(df_effect[['Feature',"Cliff's delta (mean)",'Magnitude','MWU p (mean)']].to_string(
    index=False, float_format='%.4f'))

print(f"""
VOLCANO SUMMARY (RRL)
  Significant peptides : {n_sig}  (FDR<0.05, |log2FC|>0.5)
  Up in TB+            : {n_up}
  Up in TB-            : {n_dn}

FILES SAVED
  rrl_peptide_selection_consensus.csv
  rrl_peptide_volcano_data.csv
  rrl_results_effect_sizes.csv
  rrl_results_final_summary.csv
  RRL_Fig1_auc_comparison.png
  RRL_Fig2_HIV_effect.png
  RRL_Fig3_volcano_consensus.png
""")

print('='*70)
print('Pipeline complete.')

# %% [code]


# %% [code]
```
