## Supplementary material for "High-density peptide arrays detect tuberculosis through immune remodeling, not only antigen recognition alone": code: TB_Publication_Figures.ipynb

```
# %% [markdown]
# Publication Figures — All Main + Supplementary
## For: TB Immune Remodeling Paper

Run each cell to generate one figure. Adjust colors, sizes, labels as needed.

### Main Figures (2–8)
- Fig 2: Entropy × Gini phase space (HIV effect, 3 libraries)
- Fig 3: TB severity AUC + sweep
- Fig 4: Severity in phase space
- Fig 5: Top peptides TBC vs RRL + fold-change
- Fig 6: Sensitivity / Specificity with CI
- Fig 7: Permutation test histograms
- Fig 8: LOCO bar chart

### Supplementary Figures (S1–S4)
- Fig S1: Peptide number sweep curves
- Fig S2: Combined model comparison
- Fig S3: PC2 × Mahalanobis phase space
- Fig S4: Effect size bars by severity

*(Figure 1 = study design schematic, not generated here)*

# %% [markdown]
## Step 0 — Imports & Data Loading
Run this cell first. All figures depend on it.

# %% [code]
import numpy as np
import pandas as pd
import matplotlib.pyplot as plt
import matplotlib.patches as mpatches
from matplotlib.patches import Ellipse
import seaborn as sns
import warnings
warnings.filterwarnings('ignore')
from collections import Counter
from scipy.stats import mannwhitneyu
from sklearn.ensemble import RandomForestClassifier
from sklearn.linear_model import LogisticRegression, SGDClassifier
from sklearn.preprocessing import StandardScaler
from sklearn.decomposition import PCA
from sklearn.model_selection import StratifiedKFold, GridSearchCV
from sklearn.metrics import roc_auc_score, roc_curve

RANDOM_STATE = 42
np.random.seed(RANDOM_STATE)

# ── Style settings (adjust here for all figures) ──
plt.rcParams.update({
    'figure.dpi': 150,
    'font.size': 11,
    'axes.titlesize': 12,
    'axes.labelsize': 11,
    'savefig.dpi': 300,       # publication quality
    'savefig.bbox': 'tight',
})
sns.set_style('whitegrid')

# ── Color palette (adjust here for all figures) ──
COL_TB_NEG   = '#2196F3'    # blue
COL_TB_POS   = '#E53935'    # red
COL_SC_POS   = '#F44336'    # red (S+C+)
COL_SC_NEG   = '#FF9800'    # orange (S-C+)
COL_NEG      = '#2196F3'    # blue (Negative)
COL_HIV_NEG  = '#4CAF50'    # green
COL_HIV_POS  = '#FF9800'    # orange
COL_REMOD    = '#2196F3'    # blue
COL_PEPTIDE  = '#FF9800'    # orange
COL_COMBINED = '#4CAF50'    # green
COL_TBC      = '#F44336'    # red
COL_RRL      = '#2196F3'    # blue
FILL_TB_NEG  = '#BBDEFB'
FILL_TB_POS  = '#FFCDD2'
FILL_SC_POS  = '#FFCDD2'
FILL_SC_NEG  = '#FFE0B2'
FILL_NEG     = '#BBDEFB'

# ── Load data ──
DATA_PATH = r"D:/Users/ha6057/Desktop/TBC_AI/Files for Simulation/all_signals_TBC.csv"      # peptide signals
META_PATH = r"D:/Users/ha6057/Desktop/TBC_AI/Files for Simulation/df_samples_all.xls"
PEP_CAT_PATH = r"D:/Users/ha6057/Desktop/TBC_AI/Files for Simulation/df_peptide_origin_with_Category.xlsx"

df_raw = pd.read_csv(DATA_PATH, index_col=0)
all_peptides = df_raw.columns.tolist()
X_all = df_raw.values.astype(np.float64)
sample_ids = df_raw.index.tolist()
n_samples, n_peptides = X_all.shape

meta = pd.read_csv(META_PATH).set_index('ID').loc[sample_ids]
tb_status = meta['TB Status'].values
y_tb = np.array([0 if v == 'Negative' else 1 for v in tb_status], dtype=int)
hiv = np.array([1 if v == 'HIV+' else 0 for v in meta['HIV_status']], dtype=int)
country = meta['Country'].apply(lambda x: 'Vietnam' if 'iet' in str(x) else x).values
hn = (hiv == 0)

pep_cat = pd.read_excel(PEP_CAT_PATH)
seq2cat = dict(zip(pep_cat['Sequence'], pep_cat['Category']))
seq2origin = dict(zip(pep_cat['Sequence'], pep_cat['Origin']))
tbc_seqs = [s for s in pep_cat[pep_cat['Category']=='TBC (Mycobacterium tuberculosis complex)']['Sequence'] if s in all_peptides]
rrl_seqs = [s for s in pep_cat[pep_cat['Category']=='Random Reference Library (RRL2)']['Sequence'] if s in all_peptides]
X_tbc = df_raw[tbc_seqs].values.astype(np.float64)
X_rrl = df_raw[rrl_seqs].values.astype(np.float64)

# ── Helpers ──
def entropy(X):
    s = X.sum(1, keepdims=True); s = np.where(s==0, 1, s)
    p = np.clip(X/s, 1e-12, None); return -np.sum(p * np.log(p), 1)

def gini(X):
    g = []
    for r in X:
        x = np.sort(r); c = np.cumsum(x)
        g.append((len(x)+1-2*c.sum()/c[-1])/len(x) if c[-1]>0 else 0)
    return np.array(g)

def compute_Z(X):
    e = entropy(X); g = gini(X)
    sc = StandardScaler(); Xsc = sc.fit_transform(X)
    pca = PCA(n_components=2, random_state=RANDOM_STATE)
    pcs = pca.fit_transform(Xsc)
    ref = pcs[y_tb==0]; mu = ref.mean(0)
    cov = np.cov(ref, rowvar=False) + 1e-3*np.eye(2); ci = np.linalg.inv(cov)
    mah = np.array([np.sqrt((x-mu)@ci@(x-mu)) for x in pcs])
    return np.column_stack([e, g, pcs[:,0], pcs[:,1], mah])

def confidence_ellipse(x, y, ax, n_std=1.5, facecolor='none', **kwargs):
    if len(x) < 3: return
    cov = np.cov(x, y)
    vals, vecs = np.linalg.eigh(cov)
    vals = np.maximum(vals, 1e-6)
    order = vals.argsort()[::-1]
    vals = vals[order]; vecs = vecs[:, order]
    angle = np.degrees(np.arctan2(vecs[1,0], vecs[0,0]))
    w, h = 2 * n_std * np.sqrt(vals)
    ell = Ellipse(xy=(np.mean(x), np.mean(y)), width=w, height=h, angle=angle,
                  facecolor=facecolor, **kwargs)
    ax.add_patch(ell)

def build_Z_cv(X_tr, X_te, y_tr):
    sc = StandardScaler(); xs = sc.fit_transform(X_tr); xt = sc.transform(X_te)
    pca = PCA(n_components=2, random_state=RANDOM_STATE)
    pt = pca.fit_transform(xs); pe = pca.transform(xt)
    ref = pt[y_tr==0]; mu = ref.mean(0)
    cov = np.cov(ref, rowvar=False) + 1e-3*np.eye(2); ci = np.linalg.inv(cov)
    mt = np.array([np.sqrt((x-mu)@ci@(x-mu)) for x in pt])
    me = np.array([np.sqrt((x-mu)@ci@(x-mu)) for x in pe])
    Zt = np.column_stack([entropy(X_tr),gini(X_tr),pt[:,0],pt[:,1],mt])
    Ze = np.column_stack([entropy(X_te),gini(X_te),pe[:,0],pe[:,1],me])
    sz = StandardScaler(); return sz.fit_transform(Zt), sz.transform(Ze)

Z_LABELS = ['Entropy', 'Gini', 'PC1', 'PC2', 'Mahalanobis']
cv5 = StratifiedKFold(n_splits=5, shuffle=True, random_state=RANDOM_STATE)
inner_cv = StratifiedKFold(n_splits=5, shuffle=True, random_state=RANDOM_STATE)
C_GRID = {'C': [0.001, 0.01, 0.1, 1, 10, 100]}
RF_P = dict(n_estimators=500, max_depth=None, min_samples_leaf=3,
            class_weight='balanced', random_state=RANDOM_STATE, n_jobs=-1)

# Precompute Z for all libraries
Z_all = compute_Z(X_all)
Z_tbc = compute_Z(X_tbc)
Z_rrl = compute_Z(X_rrl)

print(f'Data loaded: {n_samples} samples × {n_peptides} peptides')
print(f'TBC: {len(tbc_seqs)}, RRL: {len(rrl_seqs)}')
print(f'TB+: {y_tb.sum()}, TB-: {(y_tb==0).sum()}, HIV+: {hiv.sum()}, HIV-: {hn.sum()}')
print('Z-vectors computed for ALL, TBC, RRL')
print('\nReady — run any figure cell below.')

# %% [markdown]
## Figure 2 — Entropy × Gini Phase Space: HIV Effect
**Main text.** Shows TB+/TB- separation across 3 libraries, Full vs HIV-.  
Top row: Full cohort. Bottom row: HIV- only. Columns: ALL, TBC, RRL.

# %% [code]
fig, axes = plt.subplots(2, 3, figsize=(21, 14))

TB_COL = {0: COL_TB_NEG, 1: COL_TB_POS}
TB_FILL = {0: FILL_TB_NEG, 1: FILL_TB_POS}
TB_NAME = {0: 'TB\u2212', 1: 'TB+'}
HIV_MARKER = {0: 'o', 1: '^'}

for ci, (lib_name, Z) in enumerate([
    ('ALL (6,936 peptides)', Z_all),
    ('TBC (4,317 peptides)', Z_tbc),
    ('RRL (2,496 peptides)', Z_rrl)]):

    for ri, (cohort, mask, ax) in enumerate([
        ('Full cohort (n=105)', np.ones(len(y_tb), dtype=bool), axes[0, ci]),
        ('HIV\u2212 only (n=70)', hn, axes[1, ci])]):

        xi, yi = 0, 1  # Entropy, Gini

        for tb in [0, 1]:
            m = (y_tb == tb) & mask
            x_pts = Z[m, xi]; y_pts = Z[m, yi]

            ax.scatter(x_pts, y_pts, c=TB_COL[tb], s=55, alpha=0.65,
                      edgecolors='white', linewidths=0.5, zorder=3,
                      label=f'{TB_NAME[tb]} (n={m.sum()})')

            cx, cy = np.mean(x_pts), np.mean(y_pts)
            ax.scatter(cx, cy, marker='*', c=TB_COL[tb], s=400,
                      edgecolors='black', linewidths=1.5, zorder=5)

            confidence_ellipse(x_pts, y_pts, ax, n_std=1.5,
                             facecolor=TB_FILL[tb], edgecolor=TB_COL[tb],
                             linewidth=2.5, alpha=0.25, zorder=1)
            confidence_ellipse(x_pts, y_pts, ax, n_std=1.0,
                             facecolor='none', edgecolor=TB_COL[tb],
                             linewidth=1.5, linestyle='--', alpha=0.6, zorder=2)

        # HIV markers (full cohort only)
        if 'Full' in cohort:
            hiv_m = (hiv == 1) & mask
            ax.scatter(Z[hiv_m, xi], Z[hiv_m, yi], c='none',
                      edgecolors='black', s=80, marker='x', linewidths=1.5,
                      alpha=0.6, zorder=4, label='HIV+')

        # Arrow TB- → TB+
        cx0 = np.mean(Z[(y_tb==0)&mask, xi]); cy0 = np.mean(Z[(y_tb==0)&mask, yi])
        cx1 = np.mean(Z[(y_tb==1)&mask, xi]); cy1 = np.mean(Z[(y_tb==1)&mask, yi])
        ax.annotate('', xy=(cx1, cy1), xytext=(cx0, cy0),
                   arrowprops=dict(arrowstyle='->', color='black', lw=2.5,
                                   connectionstyle='arc3,rad=0.1'))

        ax.set_xlabel('Entropy (antibody diversity)')
        ax.set_ylabel('Gini coefficient (concentration)')
        ax.legend(fontsize=8, loc='lower left')
        ax.set_title(f'{lib_name}\n{cohort}', fontweight='bold')

fig.suptitle('Figure 2 — Immune Remodeling Phase Space: Entropy × Gini\n'
             '\u2605 = centroid, ellipses = 1\u03c3 (dashed) / 1.5\u03c3 (filled), '
             'arrow = TB\u2212 \u2192 TB+ shift',
             fontweight='bold', fontsize=14)
plt.tight_layout()
plt.savefig('Fig2_phase_entropy_gini.png', dpi=300)
plt.show()
print('Saved: Fig2_phase_entropy_gini.png')

# %% [markdown]
## Figure 3 — TB Severity: AUC Comparison + Sweep
**Main text.** Shows AUC for All TB+ / S+C+ / S-C+, Full vs HIV-.  
⚠ This cell runs the full CV pipeline — takes ~2 minutes.

# %% [code]
tasks = {
    'All TB+ vs Neg': {'pos': ['S+C+','S-C+'], 'neg': ['Negative']},
    'S+C+ vs Neg':    {'pos': ['S+C+'],        'neg': ['Negative']},
    'S\u2212C+ vs Neg':   {'pos': ['S-C+'],        'neg': ['Negative']},
}
SWEEP_NS = [1, 2, 3, 5, 10, 20, 50]
sev_results = {}

for task_name, task_def in tasks.items():
    for coh_name, c_mask in [('Full', np.ones(n_samples, dtype=bool)), ('HIV\u2212', hn)]:
        key = f'{task_name} | {coh_name}'
        pm = np.isin(tb_status, task_def['pos']) & c_mask
        nm = np.isin(tb_status, task_def['neg']) & c_mask
        tm = pm | nm
        X_t = X_all[tm]; y_t = np.where(np.isin(tb_status[tm], task_def['pos']), 1, 0)
        n_pos = y_t.sum(); n_neg = (y_t==0).sum()
        if n_pos < 5 or n_neg < 5: continue

        # Remodeling
        ra = []
        for tri, tei in cv5.split(X_t, y_t):
            Ztr, Zte = build_Z_cv(X_t[tri], X_t[tei], y_t[tri])
            lr = LogisticRegression(C=1, max_iter=2000, random_state=RANDOM_STATE, solver='lbfgs')
            lr.fit(Ztr, y_t[tri])
            ra.append(roc_auc_score(y_t[tei], lr.predict_proba(Zte)[:,1]))

        # EN→RF sweep
        sw = {n: [] for n in SWEEP_NS}
        for tri, tei in cv5.split(X_t, y_t):
            sc = StandardScaler(); Xtr = sc.fit_transform(X_t[tri]); Xte = sc.transform(X_t[tei])
            en = SGDClassifier(loss='log_loss', penalty='elasticnet', l1_ratio=0.5,
                               alpha=0.001, max_iter=5000, random_state=RANDOM_STATE,
                               class_weight='balanced')
            en.fit(Xtr, y_t[tri]); mk = np.abs(en.coef_.ravel()) > 0
            Xtr_en = Xtr[:,mk]; Xte_en = Xte[:,mk]
            rf = RandomForestClassifier(**RF_P); rf.fit(Xtr_en, y_t[tri])
            ranked = np.argsort(rf.feature_importances_)[::-1]
            for N in SWEEP_NS:
                top = ranked[:min(N, len(ranked))]
                lr2 = LogisticRegression(C=1, max_iter=2000, random_state=RANDOM_STATE, solver='lbfgs')
                lr2.fit(Xtr_en[:,top], y_t[tri])
                sw[N].append(roc_auc_score(y_t[tei], lr2.predict_proba(Xte_en[:,top])[:,1]))

        sev_results[key] = {
            'remod': np.mean(ra), 'remod_sd': np.std(ra),
            'sweep': {n: np.mean(sw[n]) for n in SWEEP_NS},
            'n_pos': n_pos, 'n_neg': n_neg}
        print(f'{key}: Remod={np.mean(ra):.3f}')

# ── Plot ──
fig, axes = plt.subplots(2, 3, figsize=(20, 12))
task_list = list(tasks.keys())
task_colors = {'All TB+ vs Neg': COL_TB_POS, 'S+C+ vs Neg': COL_SC_POS, 'S\u2212C+ vs Neg': COL_SC_NEG}

for ti, task_name in enumerate(task_list):
    # Row 1: AUC bars
    ax = axes[0, ti]
    models = ['Remodeling', 'Peptide LR\n(EN\u2192RF)']
    for coh, alpha, hatch, label in [('Full', 0.5, '', 'Full'), ('HIV\u2212', 1.0, '///', 'HIV\u2212')]:
        key = f'{task_name} | {coh}'
        if key not in sev_results: continue
        r = sev_results[key]
        vals = [r['remod'], np.max([r['sweep'][n] for n in SWEEP_NS])]
        sds = [r['remod_sd'], 0]
        x = np.arange(2); w = 0.32
        offset = -w/2 if coh == 'Full' else w/2
        for i in range(2):
            ax.bar(x[i]+offset, vals[i], w, yerr=sds[i],
                   color=task_colors[task_name], edgecolor='black',
                   alpha=alpha, hatch=hatch, capsize=5)
            ax.text(x[i]+offset, vals[i]+sds[i]+0.02, f'{vals[i]:.3f}',
                    ha='center', fontsize=8, fontweight='bold',
                    color='darkred' if coh=='HIV\u2212' else 'black')

    ax.axhline(0.5, color='red', ls='--', lw=1.5, alpha=0.5)
    ax.set_xticks(x); ax.set_xticklabels(models)
    ax.set_ylim(0.25, 1.0); ax.set_ylabel('AUC')
    npos = sev_results.get(f'{task_name} | Full', {}).get('n_pos', '?')
    nneg = sev_results.get(f'{task_name} | Full', {}).get('n_neg', '?')
    ax.set_title(f'{task_name}\n(pos={npos}, neg={nneg})', fontweight='bold')

    # Row 2: Sweep
    ax2 = axes[1, ti]
    for coh, color, marker, ls in [('Full', COL_PEPTIDE, 'o', '-'),
                                     ('HIV\u2212', COL_HIV_NEG, 's', '--')]:
        key = f'{task_name} | {coh}'
        if key not in sev_results: continue
        r = sev_results[key]
        mn = [r['sweep'][n] for n in SWEEP_NS]
        ax2.plot(SWEEP_NS, mn, marker=marker, color=color, lw=2, ls=ls,
                markersize=5, label=coh)
        bn = SWEEP_NS[np.argmax(mn)]; ba = max(mn)
        ax2.annotate(f'N={bn}\n{ba:.3f}', xy=(bn, ba), fontsize=8,
                    fontweight='bold', color=color)

    ax2.axhline(0.5, color='red', ls=':', lw=1, alpha=0.5)
    ax2.set_xscale('log'); ax2.set_xticks(SWEEP_NS)
    ax2.set_xticklabels([str(n) for n in SWEEP_NS])
    ax2.set_xlabel('N peptides'); ax2.set_ylabel('AUC')
    ax2.set_ylim(0.25, 1.0); ax2.legend(fontsize=9)
    ax2.set_title(f'{task_name} \u2014 Sweep', fontweight='bold')

leg = [mpatches.Patch(fc='gray', alpha=0.5, label='Full'),
       mpatches.Patch(fc='gray', hatch='///', label='HIV\u2212 only')]
axes[0,0].legend(handles=leg, fontsize=8, loc='upper left')

fig.suptitle('Figure 3 \u2014 TB Severity Analysis', fontweight='bold', fontsize=14)
plt.tight_layout()
plt.savefig('Fig3_severity.png', dpi=300)
plt.show()
print('Saved: Fig3_severity.png')

# %% [markdown]
## Figure 4 — Severity in Phase Space (HIV- only)
**Main text.** Neg / S+C+ / S-C+ with centroids and ellipses.

# %% [code]
SEV_COL = {'Negative': COL_NEG, 'S+C+': COL_SC_POS, 'S-C+': COL_SC_NEG}
SEV_FILL = {'Negative': FILL_NEG, 'S+C+': FILL_SC_POS, 'S-C+': FILL_SC_NEG}

fig, axes = plt.subplots(2, 3, figsize=(21, 14))

for ci, (lib_name, Z) in enumerate([
    ('ALL (6,936)', Z_all), ('TBC (4,317)', Z_tbc), ('RRL (2,496)', Z_rrl)]):

    for ri, (pair_name, xi, yi, xl, yl) in enumerate([
        ('Entropy × Gini', 0, 1, 'Entropy', 'Gini'),
        ('PC1 × PC2', 2, 3, 'PC1', 'PC2')]):

        ax = axes[ri, ci]
        for sev in ['Negative', 'S+C+', 'S-C+']:
            m = (tb_status == sev) & hn
            if m.sum() < 3: continue
            x_pts = Z[m, xi]; y_pts = Z[m, yi]

            ax.scatter(x_pts, y_pts, c=SEV_COL[sev], s=65, alpha=0.7,
                      edgecolors='white', linewidths=0.5, zorder=3,
                      label=f'{sev} (n={m.sum()})')
            cx, cy = np.mean(x_pts), np.mean(y_pts)
            ax.scatter(cx, cy, marker='*', c=SEV_COL[sev], s=450,
                      edgecolors='black', linewidths=1.5, zorder=5)
            confidence_ellipse(x_pts, y_pts, ax, n_std=1.5,
                             facecolor=SEV_FILL[sev], edgecolor=SEV_COL[sev],
                             linewidth=2.5, alpha=0.2, zorder=1)
            confidence_ellipse(x_pts, y_pts, ax, n_std=1.0,
                             facecolor='none', edgecolor=SEV_COL[sev],
                             linewidth=1.5, linestyle='--', alpha=0.5, zorder=2)

        ax.set_xlabel(xl); ax.set_ylabel(yl)
        ax.legend(fontsize=9)
        ax.set_title(f'{lib_name} \u2014 {pair_name}', fontweight='bold')

fig.suptitle('Figure 4 \u2014 TB Severity in Phase Space (HIV\u2212 Only)\n'
             '\u2605 = centroid, ellipses = 1\u03c3 / 1.5\u03c3',
             fontweight='bold', fontsize=14)
plt.tight_layout()
plt.savefig('Fig4_severity_phase.png', dpi=300)
plt.show()
print('Saved: Fig4_severity_phase.png')

# %% [markdown]
## Figure 5 — Diagnostic Peptides: TBC vs RRL + Fold-Change
**Main text.** ⚠ Runs EN→RF selection — takes ~1 minute.

# %% [code]
# ← INSERT THESE 3 LINES HERE (top of the cell)
import warnings
from sklearn.exceptions import ConvergenceWarning
warnings.filterwarnings('ignore', category=ConvergenceWarning)

# Run peptide selection for S-C+ HIV-
pm = (tb_status == 'S-C+') & hn; nm = (tb_status == 'Negative') & hn; tm = pm | nm
X_t = X_all[tm]; y_t = np.where(tb_status[tm] == 'S-C+', 1, 0)

all_tops = []; all_imp = []
for tri, tei in cv5.split(X_t, y_t):
    sc = StandardScaler(); Xtr = sc.fit_transform(X_t[tri])
    en = SGDClassifier(loss='log_loss', penalty='elasticnet', l1_ratio=0.5,
                       alpha=0.001, max_iter=5000, random_state=RANDOM_STATE,
                       class_weight='balanced')
    en.fit(Xtr, y_t[tri]); mk = np.abs(en.coef_.ravel()) > 0
    si = np.where(mk)[0]; Xtr_en = Xtr[:,mk]
    rf = RandomForestClassifier(**RF_P); rf.fit(Xtr_en, y_t[tri])
    ranked = np.argsort(rf.feature_importances_)[::-1][:20]
    names = [all_peptides[si[i]] for i in ranked]
    imp = {all_peptides[si[i]]: rf.feature_importances_[i] for i in ranked}
    all_tops.append(names); all_imp.append(imp)

freq = Counter([p for fold in all_tops for p in fold])
rows = []
for pep, count in freq.most_common(20):
    cat = seq2cat.get(pep, '?')
    cat_short = 'TBC' if 'TBC' in cat else ('RRL' if 'RRL' in cat else 'Other')
    pidx = all_peptides.index(pep)
    mp = X_t[y_t==1, pidx].mean(); mn = X_t[y_t==0, pidx].mean()
    fc = np.log2((mp+1e-9)/(mn+1e-9))
    _, p_mwu = mannwhitneyu(X_t[y_t==1,pidx], X_t[y_t==0,pidx], alternative='two-sided')
    rows.append({'Peptide':pep, 'Folds':count, 'Cat':cat_short,
                 'log2FC':fc, 'p':p_mwu, 'Origin':seq2origin.get(pep,'')})
df_pep = pd.DataFrame(rows)

# ── Plot: 2 panels ──
fig, (ax1, ax2) = plt.subplots(1, 2, figsize=(18, 9))

# Left: Folds selected
top20 = df_pep.head(15)
colors = [COL_TBC if c=='TBC' else COL_RRL for c in top20['Cat']]
ax1.barh(range(len(top20)), top20['Folds'], color=colors, edgecolor='black', alpha=0.85)
ax1.set_yticks(range(len(top20)))
ax1.set_yticklabels([f'{r["Peptide"]} ({r["Cat"]})' for _, r in top20.iterrows()], fontsize=8)
ax1.invert_yaxis()
ax1.set_xlabel('Folds selected (out of 5)')
#ax1.set_title('A | Top 20 Peptides for S\u2212C+ Detection (HIV\u2212)', fontweight='bold')
for i, (_, r) in enumerate(top20.iterrows()):
    ax1.text(r['Folds']+0.05, i, f'{r["Folds"]}/5', va='center', fontsize=7.5)

# Right: log2 fold change
top15 = df_pep.head(15)                   # ← removed .sort_values('log2FC')
colors_fc = [COL_TBC if c=='TBC' else COL_RRL for c in top15['Cat']]
ax2.barh(range(len(top15)), top15['log2FC'], color=colors_fc, edgecolor='black', alpha=0.8)
ax2.set_yticks(range(len(top15)))
labels = [r['Peptide'] for _, r in top15.iterrows()]
ax2.set_yticklabels(labels, fontsize=8)
ax2.axvline(0, color='black', lw=1)
ax2.invert_yaxis()                         # ← ADD: same top-to-bottom as left panel
ax2.set_xlabel('log2 Fold Change (S\u2212C+ / Negative)')

# ← REMOVED: the for-loop with sig = '***' / ax2.text(...)

leg = [mpatches.Patch(fc=COL_TBC, ec='black', label='TBC (M. tuberculosis)'),
       mpatches.Patch(fc=COL_RRL, ec='black', label='RRL (host proteome)')]
ax1.legend(handles=leg, fontsize=9, loc='lower right')

#fig.suptitle('Figure 5 \u2014 Diagnostic Peptides: Pathogen + Host-Proteome Converge',
#             fontweight='bold', fontsize=14)
plt.tight_layout()
plt.savefig('Fig5_peptides.png', dpi=300)
plt.show()
print('Saved: Fig5_peptides.png')

# %% [markdown]
## Figure 6 — Sensitivity / Specificity with 95% CI
**Main text.** From precomputed values (no CV needed).

# %% [code]
# Precomputed values from bootstrap analysis
data = {
    'Remodeling (5f)':  {'AUC':0.706,'AUC_lo':0.522,'AUC_hi':0.867,
                         'Sens':0.857,'Sens_lo':0.643,'Sens_hi':1.000,
                         'Spec':0.512,'Spec_lo':0.357,'Spec_hi':0.667},
    '3 peptides':       {'AUC':0.831,'AUC_lo':0.690,'AUC_hi':0.951,
                         'Sens':0.643,'Sens_lo':0.375,'Sens_hi':0.889,
                         'Spec':0.951,'Spec_lo':0.875,'Spec_hi':1.000},
    '5 peptides':       {'AUC':0.834,'AUC_lo':0.697,'AUC_hi':0.946,
                         'Sens':0.643,'Sens_lo':0.368,'Sens_hi':0.889,
                         'Spec':0.902,'Spec_lo':0.800,'Spec_hi':0.979},
    '10 peptides':      {'AUC':0.843,'AUC_lo':0.709,'AUC_hi':0.952,
                         'Sens':0.643,'Sens_lo':0.368,'Sens_hi':0.889,
                         'Spec':0.927,'Spec_lo':0.837,'Spec_hi':1.000},
}

fig, ax = plt.subplots(figsize=(12, 7))
mods = list(data.keys()); x = np.arange(len(mods)); w = 0.25

for mi, mn in enumerate(mods):
    d = data[mn]
    ax.bar(mi-w, d['AUC'], w, yerr=[[d['AUC']-d['AUC_lo']],[d['AUC_hi']-d['AUC']]],
           color=COL_REMOD, edgecolor='black', alpha=0.8, capsize=5,
           label='AUC' if mi==0 else '')
    ax.bar(mi, d['Sens'], w, yerr=[[d['Sens']-d['Sens_lo']],[d['Sens_hi']-d['Sens']]],
           color=COL_SC_POS, edgecolor='black', alpha=0.8, capsize=5,
           label='Sensitivity' if mi==0 else '')
    ax.bar(mi+w, d['Spec'], w, yerr=[[d['Spec']-d['Spec_lo']],[d['Spec_hi']-d['Spec']]],
           color=COL_HIV_NEG, edgecolor='black', alpha=0.8, capsize=5,
           label='Specificity' if mi==0 else '')

    ax.text(mi-w, d['AUC_hi']+0.03, f'{d["AUC"]:.3f}', ha='center', fontsize=9, fontweight='bold')
    ax.text(mi,   d['Sens_hi']+0.03, f'{d["Sens"]:.3f}', ha='center', fontsize=9, fontweight='bold')
    ax.text(mi+w, d['Spec_hi']+0.03, f'{d["Spec"]:.3f}', ha='center', fontsize=9, fontweight='bold')

ax.axhline(0.5, color='gray', ls=':', lw=1, alpha=0.5)
ax.set_xticks(x); ax.set_xticklabels(mods, fontsize=11)
ax.set_ylim(0, 1.2); ax.set_ylabel('Score')
ax.set_title('Figure 6 \u2014 S\u2212C+ vs Neg (HIV\u2212, n=55): Diagnostic Performance\n'
             '95% Bootstrap CI, Youden-optimal threshold', fontweight='bold')
ax.legend(fontsize=11)
plt.tight_layout()
plt.savefig('Fig6_sens_spec.png', dpi=300)
plt.show()
print('Saved: Fig6_sens_spec.png')

# %% [markdown]
## Figure 7 — Permutation Test Histograms
**Main text.** ⚠ Runs 200 permutations — takes ~3 minutes.

# %% [code]
N_PERM = 200

# Precompute static features for speed
e_pre = entropy(X_all); g_pre = gini(X_all)
sc_pre = StandardScaler(); pcs_pre = PCA(n_components=2, random_state=RANDOM_STATE).fit_transform(sc_pre.fit_transform(X_all))

def fast_cv(pcs, e, g, y, cv):
    aucs = []
    for tri, tei in cv.split(pcs, y):
        ref = pcs[tri][y[tri]==0]; mu = ref.mean(0)
        cov = np.cov(ref, rowvar=False) + 1e-3*np.eye(2); ci = np.linalg.inv(cov)
        mt = np.array([np.sqrt((x-mu)@ci@(x-mu)) for x in pcs[tri]])
        me = np.array([np.sqrt((x-mu)@ci@(x-mu)) for x in pcs[tei]])
        Zt = np.column_stack([e[tri],g[tri],pcs[tri,0],pcs[tri,1],mt])
        Ze = np.column_stack([e[tei],g[tei],pcs[tei,0],pcs[tei,1],me])
        sz = StandardScaler(); Zt = sz.fit_transform(Zt); Ze = sz.transform(Ze)
        lr = LogisticRegression(C=1, max_iter=2000, random_state=RANDOM_STATE, solver='lbfgs')
        lr.fit(Zt, y[tri])
        aucs.append(roc_auc_score(y[tei], lr.predict_proba(Ze)[:,1]))
    return np.mean(aucs)

# Full cohort
true_full = fast_cv(pcs_pre, e_pre, g_pre, y_tb, cv5)
rng = np.random.default_rng(RANDOM_STATE)
perm_full = np.array([fast_cv(pcs_pre, e_pre, g_pre, rng.permutation(y_tb), cv5) for _ in range(N_PERM)])
p_full = (perm_full >= true_full).sum() / N_PERM

# HIV- only
e_hn = entropy(X_all[hn]); g_hn = gini(X_all[hn])
sc2 = StandardScaler(); pcs_hn = PCA(n_components=2, random_state=RANDOM_STATE).fit_transform(sc2.fit_transform(X_all[hn]))
y_hn = y_tb[hn]
true_hn = fast_cv(pcs_hn, e_hn, g_hn, y_hn, cv5)
rng = np.random.default_rng(RANDOM_STATE)
perm_hn = np.array([fast_cv(pcs_hn, e_hn, g_hn, rng.permutation(y_hn), cv5) for _ in range(N_PERM)])
p_hn = (perm_hn >= true_hn).sum() / N_PERM

print(f'Full: AUC={true_full:.3f}, p={p_full:.4f}')
print(f'HIV-: AUC={true_hn:.3f}, p={p_hn:.4f}')

# ── Plot ──
fig, axes = plt.subplots(1, 2, figsize=(16, 6))
for ax, (name, true, perm, pv) in zip(axes, [
    ('Full Cohort (n=105)', true_full, perm_full, p_full),
    ('HIV\u2212 (n=70)', true_hn, perm_hn, p_hn)]):

    ax.hist(perm, bins=30, color='#B0BEC5', edgecolor='black', alpha=0.7, density=True,
            label=f'Null (n={N_PERM})')
    ax.axvline(true, color=COL_SC_POS, lw=3, label=f'Observed = {true:.3f}')
    ax.axvline(np.percentile(perm, 95), color=COL_PEPTIDE, lw=2, ls='--',
               label=f'95th pctl = {np.percentile(perm, 95):.3f}')
    ax.axvspan(true, ax.get_xlim()[1], alpha=0.1, color=COL_SC_POS)

    sig = '***' if pv<0.001 else '**' if pv<0.01 else '*' if pv<0.05 else 'ns'
    ax.set_xlabel('AUC'); ax.set_ylabel('Density')
    ax.set_title(f'Remodeling \u2014 {name}\np = {pv:.4f} {sig}', fontweight='bold')
    ax.legend(fontsize=10)

fig.suptitle('Figure 7 \u2014 Permutation Tests', fontweight='bold', fontsize=14)
plt.tight_layout()
plt.savefig('Fig7_permutation.png', dpi=300)
plt.show()
print('Saved: Fig7_permutation.png')

# %% [markdown]
## Figure 8 — Leave-One-Country-Out
**Main text.** ⚠ Runs 3 LOCO iterations — takes ~1 minute.

# %% [code]
# ← INSERT THESE 3 LINES HERE (top of the cell)
import warnings
from sklearn.exceptions import ConvergenceWarning
warnings.filterwarnings('ignore', category=ConvergenceWarning)

loco = {}
for tc in ['Peru', 'South Africa', 'Vietnam']:
    tr = (country != tc); te = (country == tc)
    Xtr = X_all[tr]; Xte = X_all[te]; ytr = y_tb[tr]; yte = y_tb[te]

    Ztr, Zte = build_Z_cv(Xtr, Xte, ytr)
    gs = GridSearchCV(LogisticRegression(max_iter=2000, random_state=RANDOM_STATE, solver='lbfgs'),
                      C_GRID, cv=inner_cv, scoring='roc_auc', n_jobs=-1)
    gs.fit(Ztr, ytr)
    lr = LogisticRegression(C=gs.best_params_['C'], max_iter=2000, random_state=RANDOM_STATE, solver='lbfgs')
    lr.fit(Ztr, ytr)
    auc_r = roc_auc_score(yte, lr.predict_proba(Zte)[:,1])

    sc = StandardScaler(); xtr = sc.fit_transform(Xtr); xte = sc.transform(Xte)
    en = SGDClassifier(loss='log_loss', penalty='elasticnet', l1_ratio=0.5,
                       alpha=0.001, max_iter=5000, random_state=RANDOM_STATE, class_weight='balanced')
    en.fit(xtr, ytr); mk = np.abs(en.coef_.ravel()) > 0
    xtr_en = xtr[:,mk]; xte_en = xte[:,mk]
    rf = RandomForestClassifier(**RF_P); rf.fit(xtr_en, ytr)
    ranked = np.argsort(rf.feature_importances_)[::-1]
    pa = {}
    for N in [3, 10, 50]:
        top = ranked[:min(N, len(ranked))]
        lr2 = LogisticRegression(C=1, max_iter=2000, random_state=RANDOM_STATE, solver='lbfgs')
        lr2.fit(xtr_en[:,top], ytr)
        pa[N] = roc_auc_score(yte, lr2.predict_proba(xte_en[:,top])[:,1])

    loco[tc] = {'remod': auc_r, 'pep': pa, 'n': int(te.sum()), 'tb+': int(yte.sum())}
    print(f'{tc}: Remod={auc_r:.3f} Pep50={pa[50]:.3f}')

# ── Plot ──
fig, ax = plt.subplots(figsize=(14, 7))
countries_list = ['Peru', 'South Africa', 'Vietnam']
x = np.arange(3); w = 0.18
for mi, (label, key, col) in enumerate([
    ('Remodeling', 'remod', COL_REMOD), ('3 pep', 3, COL_SC_POS),
    ('10 pep', 10, COL_HIV_NEG), ('50 pep', 50, '#9C27B0')]):
    vals = [loco[c]['remod'] if key=='remod' else loco[c]['pep'][key] for c in countries_list]
    ax.bar(x+mi*w, vals, w, color=col, edgecolor='black', alpha=0.8, label=label)
    for i, v in enumerate(vals):
        ax.text(x[i]+mi*w, v+0.015, f'{v:.2f}', ha='center', fontsize=8, fontweight='bold')

ax.axhline(0.5, color='red', ls='--', lw=2, alpha=0.5, label='Chance')
ax.set_xticks(x + 1.5*w)
ax.set_xticklabels([f'{c}\n(n={loco[c]["n"]}, TB+={loco[c]["tb+"]})' for c in countries_list])
ax.set_ylabel('AUC'); ax.set_ylim(0.1, 1.0)
ax.set_title('Figure 8 \u2014 Leave-One-Country-Out Validation', fontweight='bold', fontsize=13)
ax.legend(fontsize=9, ncol=2)
plt.tight_layout()
plt.savefig('Fig8_loco.png', dpi=300)
plt.show()
print('Saved: Fig8_loco.png')

# %% [markdown]
---
# Supplementary Figures

# %% [markdown]
## Figure S1 — Peptide Number Sweep (all libraries)

# %% [code]
# Uses sev_results from Figure 3 — run that cell first
# If not available, this shows ALL library only

fig, ax = plt.subplots(figsize=(10, 6))
for coh, ls in [('Full', '-'), ('HIV\u2212', '--')]:
    key = f'All TB+ vs Neg | {coh}'
    if key in sev_results:
        r = sev_results[key]
        mn = [r['sweep'][n] for n in SWEEP_NS]
        ax.plot(SWEEP_NS, mn, marker='o', lw=2, ls=ls, label=f'{coh}')
        ax.axhline(r['remod'], ls=':', lw=1, alpha=0.5)

ax.axhline(0.5, color='red', ls='--', lw=1, alpha=0.5)
ax.set_xscale('log'); ax.set_xticks(SWEEP_NS)
ax.set_xticklabels([str(n) for n in SWEEP_NS])
ax.set_xlabel('N peptides'); ax.set_ylabel('AUC')
ax.set_title('Fig S1 \u2014 Peptide Number Sweep (ALL library)', fontweight='bold')
ax.legend(); ax.grid(True, alpha=0.3)
plt.tight_layout()
plt.savefig('FigS1_sweep.png', dpi=300)
plt.show()

# %% [markdown]
## Figure S2 — Combined Model (Z + Peptides)

# %% [code]
# Precomputed values
combo = {
    'All TB+':  {'remod':0.711, 'pep3':0.653, 'combo3':0.650},
    'S+C+':     {'remod':0.733, 'pep3':0.605, 'combo3':0.706},
    'S\u2212C+':    {'remod':0.728, 'pep3':0.872, 'combo3':0.758},
}

fig, ax = plt.subplots(figsize=(12, 7))
tasks_c = list(combo.keys()); x = np.arange(len(tasks_c)); w = 0.25

for i, tn in enumerate(tasks_c):
    r = combo[tn]
    ax.bar(i-w, r['remod'], w, color=COL_REMOD, edgecolor='black', alpha=0.8,
           label='Remodeling (5f)' if i==0 else '')
    ax.bar(i,   r['pep3'],  w, color=COL_PEPTIDE, edgecolor='black', alpha=0.8,
           label='Peptide (3f)' if i==0 else '')
    ax.bar(i+w, r['combo3'], w, color=COL_COMBINED, edgecolor='black', alpha=0.8,
           label='Z + 3 pep (8f)' if i==0 else '')
    for j, v in enumerate([r['remod'], r['pep3'], r['combo3']]):
        ax.text(i+(j-1)*w, v+0.015, f'{v:.3f}', ha='center', fontsize=9, fontweight='bold')

ax.axhline(0.5, color='red', ls='--', lw=1, alpha=0.5)
ax.set_xticks(x); ax.set_xticklabels(tasks_c, fontsize=11)
ax.set_ylim(0.4, 1.0); ax.set_ylabel('AUC')
ax.set_title('Fig S2 \u2014 Combined Model: Z + Peptides (HIV\u2212)', fontweight='bold')
ax.legend(fontsize=10)
plt.tight_layout()
plt.savefig('FigS2_combined.png', dpi=300)
plt.show()

# %% [markdown]
## Figure S3 — PC2 × Mahalanobis Phase Space

# %% [code]
fig, axes = plt.subplots(2, 3, figsize=(21, 14))

for ci, (lib_name, Z) in enumerate([
    ('ALL', Z_all), ('TBC', Z_tbc), ('RRL', Z_rrl)]):

    for ri, (cohort, mask, ax) in enumerate([
        ('Full', np.ones(len(y_tb), dtype=bool), axes[0, ci]),
        ('HIV\u2212', hn, axes[1, ci])]):

        xi, yi = 3, 4  # PC2, Mahalanobis
        for tb in [0, 1]:
            m = (y_tb==tb) & mask
            ax.scatter(Z[m,xi], Z[m,yi], c=TB_COL[tb], s=55, alpha=0.65,
                      edgecolors='white', linewidths=0.5, zorder=3,
                      label=f'{TB_NAME[tb]} (n={m.sum()})')
            cx, cy = np.mean(Z[m,xi]), np.mean(Z[m,yi])
            ax.scatter(cx, cy, marker='*', c=TB_COL[tb], s=400,
                      edgecolors='black', linewidths=1.5, zorder=5)
            confidence_ellipse(Z[m,xi], Z[m,yi], ax, n_std=1.5,
                             facecolor=TB_FILL[tb], edgecolor=TB_COL[tb],
                             linewidth=2.5, alpha=0.25, zorder=1)

        if 'Full' in cohort:
            hiv_m = (hiv==1) & mask
            ax.scatter(Z[hiv_m,xi], Z[hiv_m,yi], c='none', edgecolors='black',
                      s=80, marker='x', linewidths=1.5, alpha=0.6, zorder=4, label='HIV+')

        ax.set_xlabel('PC2'); ax.set_ylabel('Mahalanobis')
        ax.legend(fontsize=8)
        ax.set_title(f'{lib_name} \u2014 {cohort}', fontweight='bold')

fig.suptitle('Fig S3 \u2014 PC2 × Mahalanobis Phase Space', fontweight='bold', fontsize=14)
plt.tight_layout()
plt.savefig('FigS3_pc2_mah.png', dpi=300)
plt.show()

# %% [markdown]
## Figure S4 — Effect Size Comparison by Severity

# %% [code]
# Precomputed Cliff's delta values (HIV- only)
effects = {
    'All TB+': {'Entropy':-0.435,'Gini':+0.422,'PC1':-0.334,'PC2':+0.229,'Mahalanobis':+0.201},
    'S+C+':    {'Entropy':-0.567,'Gini':+0.545,'PC1':-0.149,'PC2':+0.004,'Mahalanobis':+0.194},
    'S\u2212C+':   {'Entropy':-0.296,'Gini':+0.295,'PC1':-0.521,'PC2':+0.364,'Mahalanobis':+0.212},
}
task_colors_e = {'All TB+': COL_REMOD, 'S+C+': COL_SC_POS, 'S\u2212C+': COL_SC_NEG}

fig, ax = plt.subplots(figsize=(14, 7))
x = np.arange(len(Z_LABELS)); w = 0.25

for ti, (tn, color) in enumerate(task_colors_e.items()):
    vals = [effects[tn][z] for z in Z_LABELS]
    ax.bar(x + ti*w, vals, w, color=color, edgecolor='black', alpha=0.8, label=tn)

ax.axhline(0, color='black', lw=1)
ax.axhline(0.15, color='gray', ls=':', lw=0.8, alpha=0.5)
ax.axhline(-0.15, color='gray', ls=':', lw=0.8, alpha=0.5)
ax.set_xticks(x + w); ax.set_xticklabels(Z_LABELS, fontsize=11)
ax.set_ylabel("Cliff's delta (TB+ vs Neg)")
ax.set_title('Fig S4 \u2014 Effect Sizes by Severity (HIV\u2212 Only)', fontweight='bold')
ax.legend(fontsize=10); ax.set_ylim(-0.7, 0.7)
plt.tight_layout()
plt.savefig('FigS4_effects.png', dpi=300)
plt.show()

# %% [markdown]
---
## Summary of Generated Files

| File | Figure | Section |
|---|---|---|
| `Fig2_phase_entropy_gini.png` | Phase space HIV effect | Main Fig 2 |
| `Fig3_severity.png` | Severity AUC + sweep | Main Fig 3 |
| `Fig4_severity_phase.png` | Severity phase space | Main Fig 4 |
| `Fig5_peptides.png` | Top peptides TBC/RRL | Main Fig 5 |
| `Fig6_sens_spec.png` | Sensitivity / Specificity | Main Fig 6 |
| `Fig7_permutation.png` | Permutation histograms | Main Fig 7 |
| `Fig8_loco.png` | LOCO bars | Main Fig 8 |
| `FigS1_sweep.png` | Peptide sweep curves | Suppl S1 |
| `FigS2_combined.png` | Combined model | Suppl S2 |
| `FigS3_pc2_mah.png` | PC2 × Mahalanobis | Suppl S3 |
| `FigS4_effects.png` | Effect sizes by severity | Suppl S4 |

All figures saved at 300 DPI. Adjust colors, sizes, fonts in the color palette at the top.

# %% [code]
```
