## Supplementary material for "High-density peptide arrays detect tuberculosis through immune remodeling, not only antigen recognition alone": code: TB_Supplementary_Tables.ipynb

```
# %% [markdown]
# Supplementary Tables — All 4 Tables
## For: TB Immune Remodeling Paper

| Table | Content |
|---|---|
| Suppl. Table 1 | Top 15 peptides for S−C+ detection (sequence, category, protein, folds, log₂FC, p) |
| Suppl. Table 2 | Elastic Net feature reduction per fold per library |
| Suppl. Table 3 | Complete peptide number sweep (N=1–100 × 3 libraries × 2 cohorts) |
| Suppl. Table 4 | M. tuberculosis proteins identified by pipeline |

All values computed from data. Output: formatted display + CSV + Excel.

# %% [markdown]
## Step 0 — Setup

# %% [code]
import numpy as np
import pandas as pd
import warnings
warnings.filterwarnings('ignore')
from scipy.stats import mannwhitneyu
from statsmodels.stats.multitest import multipletests
from collections import Counter
from sklearn.ensemble import RandomForestClassifier
from sklearn.linear_model import LogisticRegression, SGDClassifier
from sklearn.preprocessing import StandardScaler
from sklearn.decomposition import PCA
from sklearn.model_selection import StratifiedKFold
from sklearn.metrics import roc_auc_score

RANDOM_STATE = 42
np.random.seed(RANDOM_STATE)

# ── File paths (adjust if needed) ──
DATA_PATH = r"D:/Users/ha6057/Desktop/TBC_AI/Files for Simulation/all_signals_TBC.csv"
META_PATH = r"D:/Users/ha6057/Desktop/TBC_AI/Files for Simulation/df_samples_all.xls"
PEP_CAT_PATH = r"D:/Users/ha6057/Desktop/TBC_AI/Files for Simulation/df_peptide_origin_with_Category.xlsx"

# ── Load ──
df_raw = pd.read_csv(DATA_PATH, index_col=0)
X_all = df_raw.values.astype(np.float64)
all_peptides = df_raw.columns.tolist()
n_samples, n_peptides = X_all.shape

meta = pd.read_csv(META_PATH).set_index('ID').loc[df_raw.index]
tb_status = meta['TB Status'].values
y_tb = np.array([0 if v == 'Negative' else 1 for v in tb_status], dtype=int)
hiv = np.array([1 if v == 'HIV+' else 0 for v in meta['HIV_status']], dtype=int)
hn = (hiv == 0)

pep_cat = pd.read_excel(PEP_CAT_PATH)
seq2cat = dict(zip(pep_cat['Sequence'], pep_cat['Category']))
seq2origin = dict(zip(pep_cat['Sequence'], pep_cat['Origin']))

tbc_seqs = [s for s in pep_cat[pep_cat['Category'].str.contains('TBC', na=False)]['Sequence'] if s in all_peptides]
rrl_seqs = [s for s in pep_cat[pep_cat['Category'].str.contains('RRL', na=False)]['Sequence'] if s in all_peptides]
X_tbc = df_raw[tbc_seqs].values.astype(np.float64)
X_rrl = df_raw[rrl_seqs].values.astype(np.float64)

cv5 = StratifiedKFold(n_splits=5, shuffle=True, random_state=RANDOM_STATE)
RF_P = dict(n_estimators=500, max_depth=None, min_samples_leaf=3,
            class_weight='balanced', random_state=RANDOM_STATE, n_jobs=-1)

# Known TB protein annotations (Rv → protein name + function)
TB_PROTEINS = {
    'Rv3804c': ('Ag85A (FbpB)', 'Major secreted antigen; mycolyl transferase; cell wall biosynthesis',
                'Vaccine candidate (MVA85A); serodiagnostic antigen'),
    'Rv1984c': ('Mpt64', 'Secreted immunogenic protein; culture filtrate protein',
                'Used in SD Bioline TB Ag MPT64 rapid test (WHO endorsed)'),
    'Rv3478':  ('PPE60', 'PE/PPE family protein; immune evasion; T-cell epitope',
                'Serodiagnostic candidate'),
    'Rv1827':  ('GarA', 'Glycogen accumulation regulator; immunodominant antigen',
                'T-cell and B-cell antigen'),
    'Rv2875':  ('MPT70', 'Secreted protein; homolog of MPT83',
                'Serodiagnostic antigen; species-specific marker'),
    'Rv0934':  ('PstS1', 'Phosphate-binding lipoprotein; 38-kDa antigen',
                'Classic serodiagnostic antigen (38-kDa test)'),
    'Rv1886c': ('Ag85B (FbpB)', 'Major secreted antigen; mycolyl transferase',
                'Vaccine candidate; serodiagnostic'),
    'Rv3875':  ('ESAT-6', 'Early secreted antigenic target 6 kDa; ESX-1 secretion',
                'IGRA diagnostic target (QuantiFERON)'),
    'Rv3874':  ('CFP-10', 'Culture filtrate protein 10 kDa; ESX-1 secretion',
                'IGRA diagnostic target (QuantiFERON)'),
}

print(f'Data: {n_samples} samples × {n_peptides} peptides')
print(f'TBC: {len(tbc_seqs)}, RRL: {len(rrl_seqs)}')
print(f'HIV-: {hn.sum()}, S-C+ HIV-: {((tb_status=="S-C+") & hn).sum()}')
print('\nReady.')

# %% [markdown]
## Supplementary Table 1 — Top Peptides for S−C+ Detection
Runs EN→RF selection for S−C+ vs Negative (HIV−), computes log₂FC and FDR-corrected p-values.  
⚠ Takes ~2 minutes.

# %% [code]
# Define task: S-C+ vs Negative, HIV-negative only
pm = (tb_status == 'S-C+') & hn
nm = (tb_status == 'Negative') & hn
tm = pm | nm
X_t = X_all[tm]
y_t = np.where(tb_status[tm] == 'S-C+', 1, 0)
n_pos = y_t.sum(); n_neg = (y_t==0).sum()
print(f'Task: S-C+ (n={n_pos}) vs Negative (n={n_neg}), HIV- only')

# Run EN→RF selection across 5 folds, track top 20 per fold
all_tops = []
all_importances = {}

for fold_i, (tri, tei) in enumerate(cv5.split(X_t, y_t)):
    sc = StandardScaler()
    Xtr = sc.fit_transform(X_t[tri])
    
    # Elastic Net
    en = SGDClassifier(loss='log_loss', penalty='elasticnet', l1_ratio=0.5,
                       alpha=0.001, max_iter=5000, random_state=RANDOM_STATE,
                       class_weight='balanced')
    en.fit(Xtr, y_t[tri])
    mk = np.abs(en.coef_.ravel()) > 0
    surv_idx = np.where(mk)[0]
    Xtr_en = Xtr[:, mk]
    
    # Random Forest
    rf = RandomForestClassifier(**RF_P)
    rf.fit(Xtr_en, y_t[tri])
    ranked = np.argsort(rf.feature_importances_)[::-1][:20]
    
    fold_top = []
    for r in ranked:
        pep_name = all_peptides[surv_idx[r]]
        fold_top.append(pep_name)
        if pep_name not in all_importances:
            all_importances[pep_name] = []
        all_importances[pep_name].append(rf.feature_importances_[r])
    
    all_tops.append(fold_top)
    print(f'  Fold {fold_i+1}: EN survivors={mk.sum()}, top 3: {fold_top[:3]}')

# Count frequency across folds
freq = Counter([p for fold in all_tops for p in fold])

# Compute log2FC and p-values for all selected peptides
rows = []
for pep, count in freq.most_common(30):
    pidx = all_peptides.index(pep)
    vals_pos = X_t[y_t==1, pidx]
    vals_neg = X_t[y_t==0, pidx]
    
    mean_pos = vals_pos.mean()
    mean_neg = vals_neg.mean()
    log2fc = np.log2((mean_pos + 1e-9) / (mean_neg + 1e-9))
    
    _, p_mwu = mannwhitneyu(vals_pos, vals_neg, alternative='two-sided')
    
    # Category
    cat_full = seq2cat.get(pep, 'Unknown')
    if 'TBC' in cat_full:
        cat_short = 'TBC'
    elif 'RRL' in cat_full:
        cat_short = 'RRL'
    else:
        cat_short = 'Other'
    
    # Origin / protein
    origin = seq2origin.get(pep, '')
    protein = ''
    for rv, (pname, _, _) in TB_PROTEINS.items():
        if rv in str(origin):
            protein = f'{rv} / {pname}'
            break
    if not protein and cat_short == 'RRL':
        protein = f'{origin} / host proteome'
    elif not protein and cat_short == 'TBC':
        protein = f'{origin}'
    
    # Mean RF importance
    mean_imp = np.mean(all_importances.get(pep, [0]))
    
    # Amino acid composition
    hydro = sum(1 for aa in pep if aa in 'AILMFWVP') / len(pep) * 100
    
    rows.append({
        'Rank': len(rows) + 1,
        'Peptide': pep,
        'Length': len(pep),
        'Category': cat_short,
        'Origin / Protein': protein,
        'Folds (of 5)': count,
        'Mean RF importance': mean_imp,
        'Mean intensity (S-C+)': mean_pos,
        'Mean intensity (Neg)': mean_neg,
        'log2FC': log2fc,
        'MWU p-value': p_mwu,
        'Hydrophobic %': hydro,
    })

df_st1 = pd.DataFrame(rows)

# FDR correction
reject, pvals_corrected, _, _ = multipletests(df_st1['MWU p-value'], method='fdr_bh')
df_st1['FDR p-value'] = pvals_corrected
df_st1['Significant'] = ['***' if p<0.001 else '**' if p<0.01 else '*' if p<0.05 else 'ns'
                          for p in pvals_corrected]

# Display top 15
print('\n' + '='*90)
print('SUPPLEMENTARY TABLE 1. Top 15 peptides for S-C+ detection (HIV-, n=55)')
print('='*90)
display_cols = ['Rank', 'Peptide', 'Category', 'Origin / Protein', 'Folds (of 5)',
                'log2FC', 'FDR p-value', 'Significant', 'Hydrophobic %']
print(df_st1[display_cols].head(15).to_string(index=False, float_format='{:.3f}'.format))

# Summary statistics
n_tbc = sum(1 for _, r in df_st1.head(15).iterrows() if r['Category'] == 'TBC')
n_rrl = sum(1 for _, r in df_st1.head(15).iterrows() if r['Category'] == 'RRL')
all_negative = all(r['log2FC'] < 0 for _, r in df_st1.head(15).iterrows())
mean_hydro_tbc = df_st1[df_st1['Category']=='TBC'].head(9)['Hydrophobic %'].mean()
mean_hydro_rrl = df_st1[df_st1['Category']=='RRL'].head(6)['Hydrophobic %'].mean()

print(f'\nSummary:')
print(f'  TBC peptides in top 15: {n_tbc}')
print(f'  RRL peptides in top 15: {n_rrl}')
print(f'  All log2FC negative: {all_negative}')
print(f'  Mean hydrophobic %: TBC={mean_hydro_tbc:.1f}%, RRL={mean_hydro_rrl:.1f}%')

# Save
df_st1.to_csv('SuppTable1_top_peptides.csv', index=False)
df_st1.to_excel('SuppTable1_top_peptides.xlsx', index=False)
print('\nSaved: SuppTable1_top_peptides.csv + .xlsx')

# %% [markdown]
## Supplementary Table 2 — Elastic Net Feature Reduction per Fold
Shows how many peptides survive EN filtering in each CV fold, for all 3 libraries.

# %% [code]
rows_st2 = []

for lib_name, X_lib, pep_list in [
    ('ALL', X_all, all_peptides),
    ('TBC', X_tbc, tbc_seqs),
    ('RRL', X_rrl, rrl_seqs)]:
    
    n_start = X_lib.shape[1]
    fold_survivors = []
    
    for fold_i, (tri, tei) in enumerate(cv5.split(X_lib, y_tb)):
        sc = StandardScaler()
        Xtr = sc.fit_transform(X_lib[tri])
        
        en = SGDClassifier(loss='log_loss', penalty='elasticnet', l1_ratio=0.5,
                           alpha=0.001, max_iter=5000, random_state=RANDOM_STATE,
                           class_weight='balanced')
        en.fit(Xtr, y_tb[tri])
        n_surv = (np.abs(en.coef_.ravel()) > 0).sum()
        fold_survivors.append(n_surv)
    
    mean_surv = np.mean(fold_survivors)
    pct = mean_surv / n_start * 100
    
    rows_st2.append({
        'Library': lib_name,
        'Start': n_start,
        'Fold 1': fold_survivors[0],
        'Fold 2': fold_survivors[1],
        'Fold 3': fold_survivors[2],
        'Fold 4': fold_survivors[3],
        'Fold 5': fold_survivors[4],
        'Mean': f'{mean_surv:.0f}',
        'SD': f'{np.std(fold_survivors):.0f}',
        '% retained': f'{pct:.1f}%',
    })
    print(f'{lib_name}: {n_start} → {mean_surv:.0f} ± {np.std(fold_survivors):.0f} ({pct:.1f}%)')

df_st2 = pd.DataFrame(rows_st2)

print('\n' + '='*90)
print('SUPPLEMENTARY TABLE 2. Elastic Net feature reduction across peptide libraries')
print('='*90)
print(df_st2.to_string(index=False))

df_st2.to_csv('SuppTable2_EN_survivors.csv', index=False)
df_st2.to_excel('SuppTable2_EN_survivors.xlsx', index=False)
print('\nSaved: SuppTable2_EN_survivors.csv + .xlsx')

# %% [markdown]
## Supplementary Table 3 — Complete Peptide Number Sweep
AUC for N = 1, 2, 3, 5, 10, 20, 50, 100 × 3 libraries × Full/HIV−.  
⚠ Takes ~5 minutes.

# %% [code]
def entropy(X):
    s = X.sum(1, keepdims=True); s = np.where(s==0, 1, s)
    p = np.clip(X/s, 1e-12, None); return -np.sum(p * np.log(p), 1)

def gini(X):
    g = []
    for r in X:
        x = np.sort(r); c = np.cumsum(x)
        g.append((len(x)+1-2*c.sum()/c[-1])/len(x) if c[-1]>0 else 0)
    return np.array(g)

def build_Z_cv(X_tr, X_te, y_tr):
    sc = StandardScaler(); xs = sc.fit_transform(X_tr); xt = sc.transform(X_te)
    pca = PCA(n_components=2, random_state=RANDOM_STATE)
    pt = pca.fit_transform(xs); pe = pca.transform(xt)
    ref = pt[y_tr==0]; mu = ref.mean(0)
    cov = np.cov(ref, rowvar=False) + 1e-3*np.eye(2); ci = np.linalg.inv(cov)
    mt = np.array([np.sqrt((x-mu)@ci@(x-mu)) for x in pt])
    me = np.array([np.sqrt((x-mu)@ci@(x-mu)) for x in pe])
    Zt = np.column_stack([entropy(X_tr),gini(X_tr),pt[:,0],pt[:,1],mt])
    Ze = np.column_stack([entropy(X_te),gini(X_te),pe[:,0],pe[:,1],me])
    sz = StandardScaler(); return sz.fit_transform(Zt), sz.transform(Ze)

SWEEP_NS = [1, 2, 3, 5, 10, 20, 50, 100]
rows_st3 = []

for lib_name, X_lib in [('ALL', X_all), ('TBC', X_tbc), ('RRL', X_rrl)]:
    for coh_name, mask in [('Full', np.ones(n_samples, bool)), ('HIV-', hn)]:
        Xc = X_lib[mask]; yc = y_tb[mask]
        
        # Remodeling AUC (reference)
        remod_aucs = []
        for tri, tei in cv5.split(Xc, yc):
            Ztr, Zte = build_Z_cv(Xc[tri], Xc[tei], yc[tri])
            lr = LogisticRegression(C=1, max_iter=2000, random_state=RANDOM_STATE, solver='lbfgs')
            lr.fit(Ztr, yc[tri])
            remod_aucs.append(roc_auc_score(yc[tei], lr.predict_proba(Zte)[:,1]))
        remod_auc = np.mean(remod_aucs)
        
        # Sweep
        sweep_results = {}
        for tri, tei in cv5.split(Xc, yc):
            sc = StandardScaler(); Xtr = sc.fit_transform(Xc[tri]); Xte = sc.transform(Xc[tei])
            en = SGDClassifier(loss='log_loss', penalty='elasticnet', l1_ratio=0.5,
                 alpha=0.001, max_iter=5000, random_state=RANDOM_STATE, class_weight='balanced')
            en.fit(Xtr, yc[tri]); mk = np.abs(en.coef_.ravel()) > 0
            Xtr_en = Xtr[:,mk]; Xte_en = Xte[:,mk]
            rf = RandomForestClassifier(**RF_P); rf.fit(Xtr_en, yc[tri])
            ranked = np.argsort(rf.feature_importances_)[::-1]
            
            for N in SWEEP_NS:
                top = ranked[:min(N, len(ranked))]
                lr2 = LogisticRegression(C=1, max_iter=2000, random_state=RANDOM_STATE, solver='lbfgs')
                lr2.fit(Xtr_en[:,top], yc[tri])
                auc = roc_auc_score(yc[tei], lr2.predict_proba(Xte_en[:,top])[:,1])
                sweep_results.setdefault(N, []).append(auc)
        
        row = {'Library': lib_name, 'Cohort': coh_name, 'n': mask.sum(),
               'Remod AUC': f'{remod_auc:.3f}'}
        for N in SWEEP_NS:
            m = np.mean(sweep_results[N]); s = np.std(sweep_results[N])
            row[f'N={N}'] = f'{m:.3f}'
            row[f'N={N}_sd'] = f'{s:.3f}'
        
        # Best N
        best_N = max(SWEEP_NS, key=lambda n: np.mean(sweep_results[n]))
        best_auc = np.mean(sweep_results[best_N])
        row['Best N'] = best_N
        row['Best AUC'] = f'{best_auc:.3f}'
        
        rows_st3.append(row)
        print(f'{lib_name} {coh_name}: Remod={remod_auc:.3f}, Best Pep N={best_N} AUC={best_auc:.3f}')

df_st3 = pd.DataFrame(rows_st3)

# Display: AUC only (no SD)
display_cols = ['Library', 'Cohort', 'n', 'Remod AUC'] + [f'N={N}' for N in SWEEP_NS] + ['Best N', 'Best AUC']
print('\n' + '='*120)
print('SUPPLEMENTARY TABLE 3. Peptide number sweep: AUC by panel size, library, and cohort')
print('='*120)
print(df_st3[display_cols].to_string(index=False))

df_st3.to_csv('SuppTable3_sweep.csv', index=False)
df_st3.to_excel('SuppTable3_sweep.xlsx', index=False)
print('\nSaved: SuppTable3_sweep.csv + .xlsx')

# %% [markdown]
## Supplementary Table 4 — M. tuberculosis Proteins Identified
Maps selected TBC peptides to their protein of origin. Uses annotations from TB_PROTEINS dict + peptide category file.

# %% [code]
# Collect all TBC peptides from Suppl Table 1
tbc_selected = df_st1[df_st1['Category'] == 'TBC'].head(15)

# Map to Rv numbers
protein_rows = {}
for _, row in tbc_selected.iterrows():
    origin = seq2origin.get(row['Peptide'], '')
    
    # Try to match Rv number
    matched_rv = None
    for rv in TB_PROTEINS:
        if rv in str(origin):
            matched_rv = rv
            break
    
    if matched_rv:
        if matched_rv not in protein_rows:
            pname, func, diag = TB_PROTEINS[matched_rv]
            protein_rows[matched_rv] = {
                'Rv number': matched_rv,
                'Protein': pname,
                'Function': func,
                'Diagnostic relevance': diag,
                'Peptides selected': [],
                'Ranks': [],
            }
        protein_rows[matched_rv]['Peptides selected'].append(row['Peptide'])
        protein_rows[matched_rv]['Ranks'].append(row['Rank'])
    else:
        # Unknown protein
        key = str(origin)[:20]
        if key not in protein_rows:
            protein_rows[key] = {
                'Rv number': str(origin),
                'Protein': 'Unknown',
                'Function': 'Not annotated',
                'Diagnostic relevance': 'Not established',
                'Peptides selected': [],
                'Ranks': [],
            }
        protein_rows[key]['Peptides selected'].append(row['Peptide'])
        protein_rows[key]['Ranks'].append(row['Rank'])

# Format
rows_st4 = []
for key, info in protein_rows.items():
    n_pep = len(info['Peptides selected'])
    rank_str = ', '.join([str(r) for r in sorted(info['Ranks'])])
    rows_st4.append({
        'Rv number': info['Rv number'],
        'Protein': info['Protein'],
        'Function': info['Function'],
        'Diagnostic relevance': info['Diagnostic relevance'],
        'N peptides': n_pep,
        'Ranks': rank_str,
        'Peptide sequences': ', '.join(info['Peptides selected']),
    })

df_st4 = pd.DataFrame(rows_st4)

print('='*100)
print('SUPPLEMENTARY TABLE 4. M. tuberculosis proteins identified by EN→RF pipeline')
print('='*100)
for _, row in df_st4.iterrows():
    print(f"\n{row['Rv number']} — {row['Protein']}")
    print(f"  Function: {row['Function']}")
    print(f"  Diagnostic: {row['Diagnostic relevance']}")
    print(f"  Peptides: {row['N peptides']} (ranks {row['Ranks']})")
    print(f"  Sequences: {row['Peptide sequences']}")

# Also create RRL summary
rrl_selected = df_st1[df_st1['Category'] == 'RRL'].head(10)
print(f'\n\nRRL (host proteome) peptides in top 15:')
for _, row in rrl_selected.iterrows():
    print(f"  Rank {row['Rank']}: {row['Peptide']} (hydrophobic {row['Hydrophobic %']:.0f}%, log2FC={row['log2FC']:.2f})")

df_st4.to_csv('SuppTable4_TB_proteins.csv', index=False)
df_st4.to_excel('SuppTable4_TB_proteins.xlsx', index=False)
print('\nSaved: SuppTable4_TB_proteins.csv + .xlsx')

# %% [markdown]
## Export All Tables to One Excel File
Creates a single Excel file with one sheet per supplementary table.

# %% [code]
with pd.ExcelWriter('Supplementary_Tables_ALL.xlsx', engine='openpyxl') as writer:
    df_st1.to_excel(writer, sheet_name='SuppTable1_Peptides', index=False)
    df_st2.to_excel(writer, sheet_name='SuppTable2_EN_Survivors', index=False)
    df_st3[display_cols].to_excel(writer, sheet_name='SuppTable3_Sweep', index=False)
    df_st4.to_excel(writer, sheet_name='SuppTable4_TB_Proteins', index=False)

print('Saved: Supplementary_Tables_ALL.xlsx')
print('  Sheet 1: SuppTable1_Peptides (top 30 peptides with all annotations)')
print('  Sheet 2: SuppTable2_EN_Survivors (EN reduction per fold)')
print('  Sheet 3: SuppTable3_Sweep (complete N sweep)')
print('  Sheet 4: SuppTable4_TB_Proteins (TB protein mapping)')
print('\nOpen in Excel to format for submission.')

# %% [markdown]
---
## Summary

| Output File | Content |
|---|---|
| `SuppTable1_top_peptides.csv/.xlsx` | Top 30 peptides: sequence, category, protein, folds, RF importance, intensities, log₂FC, FDR p |
| `SuppTable2_EN_survivors.csv/.xlsx` | EN survivors per fold (ALL/TBC/RRL × 5 folds) |
| `SuppTable3_sweep.csv/.xlsx` | Complete sweep: N=1–100 × 3 libraries × 2 cohorts, with SD |
| `SuppTable4_TB_proteins.csv/.xlsx` | TB proteins: Rv number, name, function, diagnostic relevance, peptide sequences |
| **`Supplementary_Tables_ALL.xlsx`** | **All 4 tables in one Excel file (one sheet each)** |

### Estimated run time
| Table | Time |
|---|---|
| Table 1 (Peptide selection) | ~2 min |
| Table 2 (EN survivors) | ~30 sec |
| Table 3 (Complete sweep) | ~5 min |
| Table 4 (TB proteins) | instant (uses Table 1 data) |
| **Total** | **~8 min** |
```
