## Supplementary material for "High-density peptide arrays detect tuberculosis through immune remodeling, not only antigen recognition alone": readme to use code

README – Data and Code
for
High-density peptide arrays detect tuberculosis through immune remodeling not only antigen recognition

Dimitry Schmidt, Sergey Biniaminov, Nathalie Biniaminov, Clemens von Bojnicic-Kninski, Roman Popov, Josef Maier, Hubert Bernauer, Johanna Griesbaum, Nicole Schneiderhan-Marra, Alex Dulovic*, Alexander Nesterov-Mueller*

---------------------------------------------------------------------

OVERVIEW

This archive contains all data and code required to reproduce the analysis presented in the manuscript.

The study is based on high-density peptide array measurements of antibody binding profiles, combined with machine learning analysis for disease classification and immune system remodeling.

The framework enables both peptide-level biomarker identification and global characterization of immune repertoire structure.

---------------------------------------------------------------------

DATA DESCRIPTION

1. all_signals_TBC.csv
   Raw signal intensities from peptide arrays for all samples

2. df_samples_all.xls
   Sample metadata
   - Includes:
     - sample ID
     - disease status (e.g. TBC+, TBC-, HIV+)
     - cohort information (e.g. country)

3. df_peptide_origin_with_Category.xlsx
   Peptide annotation table includes:
     - peptide sequences
     - origin (e.g. Mycobacterium tuberculosis, HIV, RRL)
     - peptide category classification

---------------------------------------------------------------------

CODE DESCRIPTION

The analysis is implemented using Python in Jupyter notebooks.

Main notebooks:

1. TB_Pipeline_All_ElasticNet.ipynb
   - Full dataset analysis pipeline
   - Includes preprocessing, feature selection, model training, and evaluation

2. TB_Pipeline_TBC_ElasticNet.ipynb
   - Analysis focused on tuberculosis-associated peptides

3. TB_Pipeline_RRL_ElasticNet.ipynb
   - Analysis based on RRL (Resemblance Ranking Library) peptides

4. TB_Publication_Figures.ipynb
   - Generates all main figures presented in the manuscript

5. TB_Publication_Tables.ipynb
   - Generates tables used in the manuscript

6. TB_Supplementary_Tables.ipynb
   - Generates supplementary tables

---------------------------------------------------------------------

ANALYSIS PIPELINE

The notebooks perform the following steps:

1. Data loading and preprocessing
   - Integration of signal intensities with sample metadata
   - Filtering and normalization

2. Feature selection
   - Elastic Net–based feature selection
   - Identification of informative peptide subsets

3. Model training
   - Classification models for disease prediction
   - Cross-validation

4. Evaluation
   - ROC curve analysis
   - AUC calculation
   - Performance metrics

5. Visualization
   - Generation of figures and tables for publication

---------------------------------------------------------------------

REPRODUCIBILITY

To reproduce the results:

1. Ensure that all files remain in the same directory structure
2. Open the relevant notebook (e.g. TB_Pipeline_All_ElasticNet.ipynb)
3. Run all cells sequentially

All file paths are relative and require no modification.

---------------------------------------------------------------------

REQUIREMENTS

Python 3.x

Required packages:
- numpy
- pandas
- scikit-learn
- matplotlib
- scipy

Jupyter Notebook environment is required.

---------------------------------------------------------------------

NOTES

- The dataset represents processed peptide array measurements used for analysis.
- The provided notebooks reproduce the main classification results, figures, and tables reported in the manuscript.
- The approach captures high-dimensional antibody binding patterns, enabling analysis beyond single antigen recognition towards global immune system remodeling.

---------------------------------------------------------------------

CONTACT

For questions regarding the data or analysis, please contact:

Alexander Nesterov-Mueller

---------------------------------------------------------------------
